# Replicated anthropogenic hybridisations reveal parallel patterns of admixture in marine mussels

**DOI:** 10.1101/590737

**Authors:** Alexis Simon, Christine Arbiol, Einar Eg Nielsen, Jérôme Couteau, Rossana Sussarellu, Thierry Burgeot, Ismaël Bernard, Joop W.P. Coolen, Jean-Baptiste Lamy, Stéphane Robert, Maria Skazina, Petr Strelkov, Henrique Queiroga, Ibon Cancio, John J. Welch, Frédérique Viard, Nicolas Bierne

## Abstract

Human-mediated transport creates secondary contacts between genetically differentiated lineages, bringing new opportunities for gene exchange. When similar introductions occur in different places, they provide informally replicated experiments for studying hybridisation. We here examined 4279 *Mytilus* mussels, sampled in Europe and genotyped with 77 ancestry informative markers. We identified a type of introduced mussels, called ‘dock mussels’, associated with port habitats and displaying a particular genetic signal of admixture between *M. edulis* and the Mediterranean lineage of *M. galloprovincialis*. These mussels exhibit similarities in their ancestry compositions, regardless of the local native genetic backgrounds and the distance separating colonised ports. We observed fine-scale genetic shifts at the port entrance, at scales below natural dispersal distance. Such sharp clines do not fit with migration-selection tension zone models, and instead suggest habitat choice and early stage adaptation to the port environment, possibly coupled with connectivity barriers. Variations in the spread and admixture patterns of dock mussels seem to be influenced by the local native genetic backgrounds encountered. We next examined departures from the average admixture rate at different loci, and compared human-mediated admixture events, to naturally admixed populations and experimental crosses. When the same *M. galloprovincialis* background was involved, positive correlations in the departures of loci across locations were found; but when different backgrounds were involved, no or negative correlations were observed. While some observed positive correlations might be best explained by a shared history and saltatory colonisation, others are likely produced by parallel selective events. Altogether, genome-wide effect of admixture seems repeatable, and more dependent on genetic background than environmental context. Our results pave the way towards further genomic analyses of admixture, and monitoring of the spread of dock mussels both at large and fine spacial scales.

## 1 Introduction

Biological introductions have evolutionary impacts on both native and introduced species, through ecological and genetic responses (Mooney & Cleland, 2001; Prentis, Wilson, Dormontt, Richardson, & Lowe, 2008; Strayer, Eviner, Jeschke, & Pace, 2006; Suarez & Tsutsui, 2008). This is especially so when ‘anthropogenic hybridisations’ lead to gene exchange (see McFarlane & Pemberton, 2019, for a recent review). Anthropogenic hybridisations have probably been underestimated, but have nevertheless been reported in diverse taxonomic groups, including plants, birds, fishes, mammals and invertebrates (Largiadèr, 2008, and references therein). For instance, in nineteen different fish families, half of the observed interspecific hybridisations have been attributed to human disturbances (Scribner, Page, & Bartron, 2000). The outcomes of these hybridisations could be similarly diverse. Hybridisation might favour the sustainable establishment of non-indigenous species (NIS) by facilitating adaptation to the local environment via the introgression of ‘ready-to-use’ alleles from native genomes. Immediate advantage could also be gained through heterosis at the initial stage of introduction (Ellstrand & Schierenbeck, 2000; Schierenbeck & Ellstrand, 2009; Suarez & Tsutsui, 2008). Conversely, hybridisation is often considered as ‘genetic pollution’ of the native species, raising concerns of ‘extinction by hybridization and introgression’ (Rhymer & Simberloff, 1996), although these concerns often neglect the possibility of genetic rescue (Fitzpatrick et al., 2019; Harris, Zhang, & Nielsen, 2019). Additionally, hybrid fitness depression might oppose introduction success, stopping the spread of the introduced lineage (Kovach et al., 2016), perhaps at a natural barrier (Barton, 1979b). Overall, the evolutionary consequences of anthropogenic hybridisation (i.e., gene flow, local introgression, reinforcement, or rescue) are likely to be strongly contingent on intrinsic and extrinsic factors, such as the accumulation of reproductive incompatibilities or local selection processes (Abbott et al., 2013).

Introductions with hybridisation can also shed light on the evolutionary process itself. Just like natural hybrid zones, human-induced hybrid zones can be seen as ‘natural laboratories for evolutionary studies’ (G. M. Hewitt, 1988, p. 158) (Abbott et al., 2013; Barton & Hewitt, 1989). Indeed, anthropogenic introductions have a special value, because they tend to be recent, informally replicated (taking place independently in different locations), and can often be compared to natural admixture events between the same lineages (Bouchemousse, Liautard-Haag, Bierne, & Viard, 2016). This is important because, even with genome-wide genetic data and powerful inferential methods, the traces of secondary contacts tend to erode over time, and can be confounded with other processes (Bertl, Ringbauer, & Blum, 2018; Bierne, Gagnaire, & David, 2013). Recent secondary contacts allow a unique window on the processes involved during the early phase of admixture, including the sorting of alleles in admixed populations (Schumer et al., 2018).

The blue mussel complex of species (*Mytilus edulis*) includes three species naturally distributed in temperate regions of the Northern hemisphere: *M. edulis* (Linnaeus 1758), *M. galloprovincialis* (Lamarck 1819) and *M. trossulus* (Gould 1850). It constitutes a model for investigating the genetic and evolutionary consequences of marine invasions (Popovic, Matias, Bierne, & Riginos, 2019; Saarman & Pogson, 2015). Despite divergences estimated at 2.5 million years (MY) between *M. edulis* and *M. galloprovincialis* (Roux et al., 2014) and 3.5 MY between these and *M. trossulus* (Rawson & Hilbish, 1995), they are incompletely reproductively isolated and readily hybridise where they meet.

Where found in sympatry, the distribution of *M. edulis* and *M. galloprovincialis* are correlated with salinity, tidal height and wave exposure (Bierne, David, Langlade, & Bonhomme, 2002; Gardner, 1994). In certain cases, *M. edulis* occupies sheltered, deeper or estuarine environments, while *M. galloprovincialis* is found on more wave-exposed parts of the coast. In regions with a single species, however, individuals can occupy all niches. It should also be noted that independent contacts can show reversed associations with the environment, in agreement with the coupling hypothesis (Bierne, Welch, Loire, Bonhomme, & David, 2011). *M. galloprovincialis*, though known as the Mediterranean mussel, has a large natural distribution – from the Black Sea to the North of the British Isles – and is divided into two main lineages, Atlantic (Atl.) and Mediterranean (Med.). (Fraïsse, Belkhir, Welch, & Bierne, 2016; Popovic et al., 2019; Quesada, Zapata, & Alvarez, 1995; Roux et al., 2014; Zbawicka, Drywa, Śmietanka, & Wenne, 2012). These two lineages form hybrid zones in the Almeria-Oran front region (El Ayari, Trigui El Menif, Hamer, Cahill, & Bierne, 2019; Quesada, Beynon, & Skibinski, 1995; Quesada, Zapata, & Alvarez, 1995).

Mussels of the family *Mytilidae* have several traits making them prone to transportation by humans. As bentho-pelagic molluscs, their planktonic feeding larval stage allows long distance spread through both marine currents (Bayne, 1976; Branch & Steffani, 2004) and anthropogenic vectors, mostly via ballast water (Geller, Carlton, & Powers, 1994) or fouling (e.g. on hulls: Apte, Holland, Godwin, and Gardner, 2000; Casoli et al., 2016; or marine litter: Miller, Carlton, Chapman, Geller, and Ruiz, 2017; Miralles, Gomez-Agenjo, Rayon-Viña, Gyraitė, and Garcia-Vazquez, 2018; Węsławski and Kotwicki, 2018). Mussels are also heavily cultivated on a global scale (287,958 tonnes in 2016, FAO, 2018); they can therefore follow the two main introduction pathways of marine species: international shipping and aquaculture (Molnar, Gamboa, Revenga, & Spalding, 2008; Nunes, Katsanevakis, Zenetos, & Cardoso, 2014). While larval dispersal might allow a post-introduction range expansion, initial establishment also relies on avoiding demographic and genetic Allee effects. As such, successful establishment depends on either large propagule pressure (likely to occur in many marine NIS: Rius, Turon, Bernardi, Volckaert, and Viard, 2015; Viard, David, and Darling, 2016), or on hybridisation with a native species (Mesgaran et al., 2016). In *Mytilus* mussels, this is facilitated by both high fecundity and high density traits, and by their incomplete reproductive isolation.

Among *Mytilus* species, *M. galloprovincialis* has been introduced many times across the globe, in both the northern and southern hemispheres, and notably, along the Pacific coast of North America, in South America, South Africa, Asia, and Oceania (Branch & Steffani, 2004; Daguin & Borsa, 2000; Han, Mao, Shui, Yanagimoto, & Gao, 2016; Kartavtsev, Chichvarkhin, Kijima, Hanzawa, & Park, 2005; Larraín, Zbawicka, Araneda, Gardner, & Wenne, 2018; McDonald, Seed, & Koehn, 1991; Saarman & Pogson, 2015; Zbawicka, Trucco, & Wenne, 2018). By contrast, we only know of a few cases of *M. edulis* introductions – either transient or successful – into non-native areas (Casoli et al., 2016; Crego-Prieto et al., 2015; Fraïsse, Haguenauer, et al., 2018). Branch and Steffani (2004) reported that observed introductions of *M. galloprovincialis* happened close to large shipping ports, with a secondary range expansion from these points. For instance in South Africa, *M. galloprovincialis* spread rapidly and had varying impacts on local communities, modulated by wave action (Branch, Odendaal, & Robinson, 2008; Branch & Steffani, 2004). Wherever *Mytilus* species are native, *M. galloprovincialis* has been shown to be highly competitive and has often displaced local mussels (James T. Carlton, Geller, Reaka-Kudla, & Norse, 1999). *M. galloprovincialis* has also been reported in the subarctic and Arctic, notably in Norway (Brooks & Farmen, 2013; Mathiesen et al., 2016). Given the low divergence between Atl. and Med. *M. galloprovincialis*, and their assignment to the same species, introduced *M. galloprovincialis* has often been reported without further investigation of its origin, and when markers are insufficiently informative, the origin is necessarily unresolved. Nevertheless, it is clear that both lineages have been successfully introduced in multiple places worldwide (Atl. in South Africa and Australia; Med. in the Eastern and Western Pacific Ocean; see Daguin and Borsa, 2000; Han et al., 2016; Popovic et al., 2019; Zardi et al., 2018).

Just as mussels are model organisms for studying the processes underlying successful introduction of alien species, ports are model locations (Bax, Hayes, Marshall, Parry, & Thresher, 2002). Because they are hubs of maritime traffic, with high connectivity, they are bridgeheads towards expansion at regional scales (Drake & Lodge, 2004). Vessels have been shown to be a major introduction pathway, through various vectors, including ballast water, sea-chest and hull (Katsanevakis, Zenetos, Belchior, & Cardoso, 2013; Sylvester et al., 2011). In addition, ports are often distinct from nearby natural habitats, with particular environmental features (Chapman & Underwood, 2011, and references therein). These new niches can be colonised by opportunistic species, such as many NIS (Bishop et al., 2017, and references therein). Mussels are likely to be introduced and become established in ports due to their aforementioned life history traits, their robustness to environmental pollution (Mlouka et al., 2019; Roberts, 1976), and tolerance to a large range of environmental conditions in terms of temperature, salinity and wave action (both through individual plasticity and interspecific variability; Braby and Somero, 2006; Fly and Hilbish, 2013; Lockwood and Somero, 2011).

In this study, using a population genomic dataset comprising 4279 mussels genotyped at 77 ancestry informative SNPs, we examined mussel populations established in ports in North-West France (located along the Atlantic and the English Channel coastlines), and compared these to mussel populations established in the vicinity. This genetic survey allows us to report, for the first time, an unexpected and extensive introduction of a non-indigenous lineage of *M. galloprovincialis* into five ports in our study area. We show that the introduced mussels have a distinctive genetic signature, originating from admixture between the Med. *M. galloprovincialis* and native *M. edulis*. We call these mussels, ‘dock mussels’, in recognition of their strong association with port environments. Dock mussel populations in ports appear to constitute stable admixed populations and form small-scale hybrid zones with native mussels at the port entrance, which can be either *M. edulis* or Atl. *M. galloprovincialis* depending on the region.

To place these populations in a wider context, we additionally analysed published and new samples of putative *M. galloprovincialis* in Norway (Mathiesen et al., 2016), and concluded that these are admixed mussels between Atl. *M. galloprovincialis* and the local North-European (North-Eu.) *M. edulis* lineage, resulting from an anthropogenic introduction. We also combined our data with multiple samples of admixed populations from natural hybrid zones, and laboratory crosses. This allowed us to compare multiple independent events of admixture, with a variety of ecological and genomic contexts.

The similarities and differences between these various admixed populations help to clarify the factors that determine the outcome of an introduction with hybridisation. In particular, we show that similar outcomes sometimes reflect shared colonisation history, but can also arise in genuinely independent colonisations. However, this predictability is highly background dependent, and replicated outcomes only appear when the same parental backgrounds are involved.

## 2 Methods

### 2.1 Sampling and genotyping

We aimed to examine mussel populations in ports, following the discovery of mussels with unexpected Med. *M. galloprovincialis* ancestry in the port of Cherbourg (France), as sampled in 2003 (Simon et al., 2019). Besides a new sampling in Cherbourg, we sampled seven additional ports and neighbouring natural populations. We also aimed to compare the admixture patterns observed in the ports to other admixed populations, involving different lineages of the same species. The sampling focused on populations where we had *a priori* expectations of admixture. Therefore, it should not be confused with a representative sample of the *M. edulis* complex, where populations are usually much closer to the reference parental populations. Most of the port sites were sampled between 2015 and 2017 and older samples were used as references or for temporal information. We either received samples from collaborators or directly sampled in the areas of interest (see Figure S1 and Table S1 for full details).

As part of our sampling process, we re-genotyped samples from several previous studies that reported the presence of *M. galloprovincialis* alleles, but had not assigned the samples to the Atl. or Med. *M. galloprovincialis* lineages. In particular, we used previously extracted DNA from the following studies: (i) Mathiesen et al. (2016) who studied the genetics of *Mytilus* spp. in the sub-Arctic and Arctic using 81 randomly ascertained SNPs. They identified *M. galloprovincialis* and putative hybrids with *M. edulis* in the Lofoten islands, Svalbard and Greenland. Their parental reference samples included only the Atl. *M. galloprovincialis* lineage (Galicia, Spain). Our aim was to further assess the origin of the *M. galloprovincialis* ancestry. (ii) Coolen (2017) studied connectivity between offshore energy installations in the North Sea, characterising samples with 6 microsatellite markers and the locus Me15/16. He identified populations containing individuals with *M. galloprovincialis* ancestry, using an Atl. *M. galloprovincialis* reference as well (Lisbon, Portugal).

Samples originating from another oil platform from the Norwegian Sea (Murchison oil station, MCH) and one Norwegian sample (Gåseid, GAS) were also included. We note that the MCH oil rig was free of settled mussels at the time of deployment.

These natural samples were compared to laboratory crosses between *M. edulis* and Med.*M. galloprovincialis*, produced in Bierne, Bonhomme, Boudry, Szulkin, and David (2006), and genotyped in Simon, Bierne, and Welch (2018). Briefly, F1 hybrids were first produced by crossing five males and five females of *M. edulis* from the North Sea (Grand-Fort-Philippe, France) and *M. galloprovincialis* from the western Mediterranean Sea (Thau lagoon, France). F2s were produced by crossing one F1 female and five F1 males. Additionally, sex-reciprocal backcrosses to *M. galloprovincialis* were made, they are named BCG when the females were *M. galloprovincialis* and BCF1 when the female was F1 (Table 1). Production of crosses are described in full detail in Bierne, David, Boudry, and Bonhomme (2002), Bierne et al. (2006) and Simon et al. (2018).

**Table 1:**
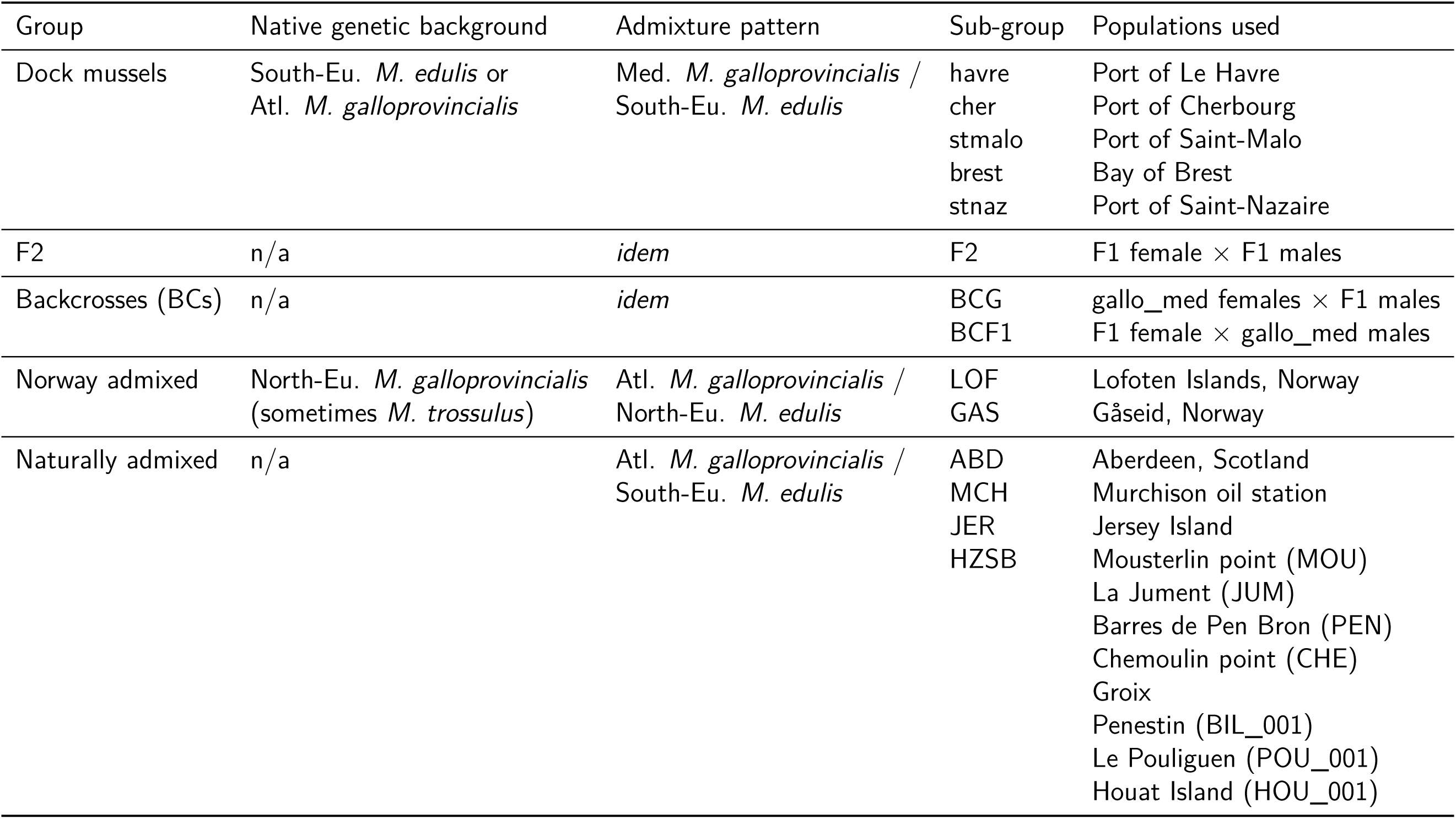
Groups used in the analyses of ancestry comparisons and correlations of distortion. The location and ancestry composition of sub-groups are indicated in Figure 2. The native genetic backgrounds possibly encountered is indicated for cases of introduction (n/a: not applicable).

| Group | Native genetic background | Admixture pattern | Sub-group | Populations used |
| --- | --- | --- | --- | --- |
| Dock mussels | South-Eu. *M. edulis* or | Med. *M. galloprovincialis* / | havre | Port of Le Havre |
|  | Atl. *M. galloprovincialis* | South-Eu. *M. edulis* | cher | Port of Cherbourg |
|  |  |  | stmalo | Port of Saint-Malo |
|  |  |  | brest | Bay of Brest |
|  |  |  | stnaz | Port of Saint-Nazaire |
| F2 | n/a | *idem* | F2 | F1 female × F1 males |
| Backcrosses (BCs) | n/a | *idem* | BCG | gallo_med females × F1 males |
|  |  |  | BCF1 | F1 female × gallo_med males |
| Norway admixed | North-Eu. *M. galloprovincialis*<br>(sometimes *M. trossulus*) | Atl. *M. galloprovincialis* / | LOF | Lofoten Islands, Norway |
|  |  | North-Eu. *M. edulis* | GAS | Gåseid, Norway |
| Naturally admixed | n/a | Atl. *M. galloprovincialis* /<br>South-Eu. *M. edulis* | ABD | Aberdeen, Scotland |
|  |  |  | MCH | Murchison oil station |
|  |  |  | JER | Jersey Island |
|  |  |  | HZSB | Mousterlin point (MOU) |
|  |  |  |  | La Jument (JUM) |
|  |  |  |  | Barres de Pen Bron (PEN) |
|  |  |  |  | Chemoulin point (CHE) |
|  |  |  |  | Groix |
|  |  |  |  | Penestin (BIL_001) |
|  |  |  |  | Le Pouliguen (POU_001) |
|  |  |  |  | Houat Island (HOU_001) |

We collected gill, mantle or hemolymph tissues from mussels either fixed in 96% ethanol or freshly collected for DNA extraction. We used the NucleoMag™ 96 Tissue kit (Macherey-Nagel) in combination with a Kingfisher Flex (serial number 711-920, ThermoFisher Scientific) extraction robot to extract DNA. We followed the kit protocol with modified volumes for the following reagents: 2*×* diluted magnetic beads, 200 µL of MB3 and MB4, 300 µL of MB5 and 100 µL of MB6. The extraction program is presented in Figure S2.

Genotyping was subcontracted to LGC genomics (Hoddesdon, UK) and performed with the KASP™ array method (Semagn, Babu, Hearne, & Olsen, 2014). We used a set of ancestry informative SNPs developed previously (Simon et al., 2018; Simon et al., 2019). For cost reduction, we used a subset of SNPs that were sufficient for species and population delineation. Multiple experiments of genotyping were performed. The results were pooled to obtain a dataset of 81 common markers.

### 2.2 Filtering

To obtain a clean starting dataset, we filtered loci and individuals for missing data. We then defined groups of individuals used as reference in downstream analyses and identified loci deviating from Hardy-Weinberg expectations, to filter used markers for analyses depending on equilibrium hypotheses.

Analyses were carried out using R (v3.5.3, R Core Team, 2019) and custom Python 3 scripts for format conversions. Software packages and versions used are listed in Table S2. Decision thresholds for all analyses and dataset selections are summarised in Table S3.

First, control individuals duplicated between genotyping experiments were removed by keeping the one having the least missing data. Over 81 markers, the maximum number of mismatches observed between two duplicated individuals was 2 (without considering missing data), showing that the genotyping method is mostly accurate. A few individuals identified as affected by a *M. trossulus* transmissible cancer were removed from the dataset (Metzger et al., 2016; Riquet, Simon, & Bierne, 2017).

The dataset was filtered for missing data with a maximum threshold of 10% for markers over all individuals and 30% for individuals over all markers. This filtering yielded 4279 individuals genotyped at 77 loci (from the initial dataset composed of 4495 individuals genotyped over 81 loci). We separated nuclear (76 loci) and mitochondrial (1 locus) markers for downstream analyses. The mitochondrial marker (named 601) is located on the female mitochondria.

Most analyses required reference population samples. A list of reference individuals and groups was set *a priori* using the literature and our knowledge of the *M. edulis* species complex (Figure 1c and Table S4). We defined three levels of structure that we call L1, L2 and L3. L1 is the species level comprising *M. edulis* (edu), *M. galloprovincialis* (gallo) and *M. trossulus* (tros). L2 defines allopatric lineages in each species: (i) American (edu_am, East coast) and European (edu_eu) *M. edulis*; (ii) Atl. (gallo_atl) and Med. (gallo_med) *M. galloprovincialis*; (iii) Pacific (tros_pac), American (tros_am, East coast) and European (tros_eu, Baltic Sea) *M. trossulus*. Finally, L3 defines sub-populations where the differentiation is mainly due to local introgression following historic contacts between lineages (Fraïsse et al., 2016): (i) North-Eu. populations of *M. edulis* (edu_eu_north) were included (Simon et al., 2019). This lineage is present along the coast of Norway and meet with the South-Eu. lineage (edu_eu_south) along the Danish coast; (ii) Atl. *M. galloprovincialis* from the Iberian peninsula (gallo_atl_iber) and mussels from Brittany (gallo_atl_brit); (iii) West (gallo_med_west) and East (gallo_med_east) Med. *M. galloprovincialis*, the limit being set at the Siculo-Tunisian strait.

**Figure 1:**
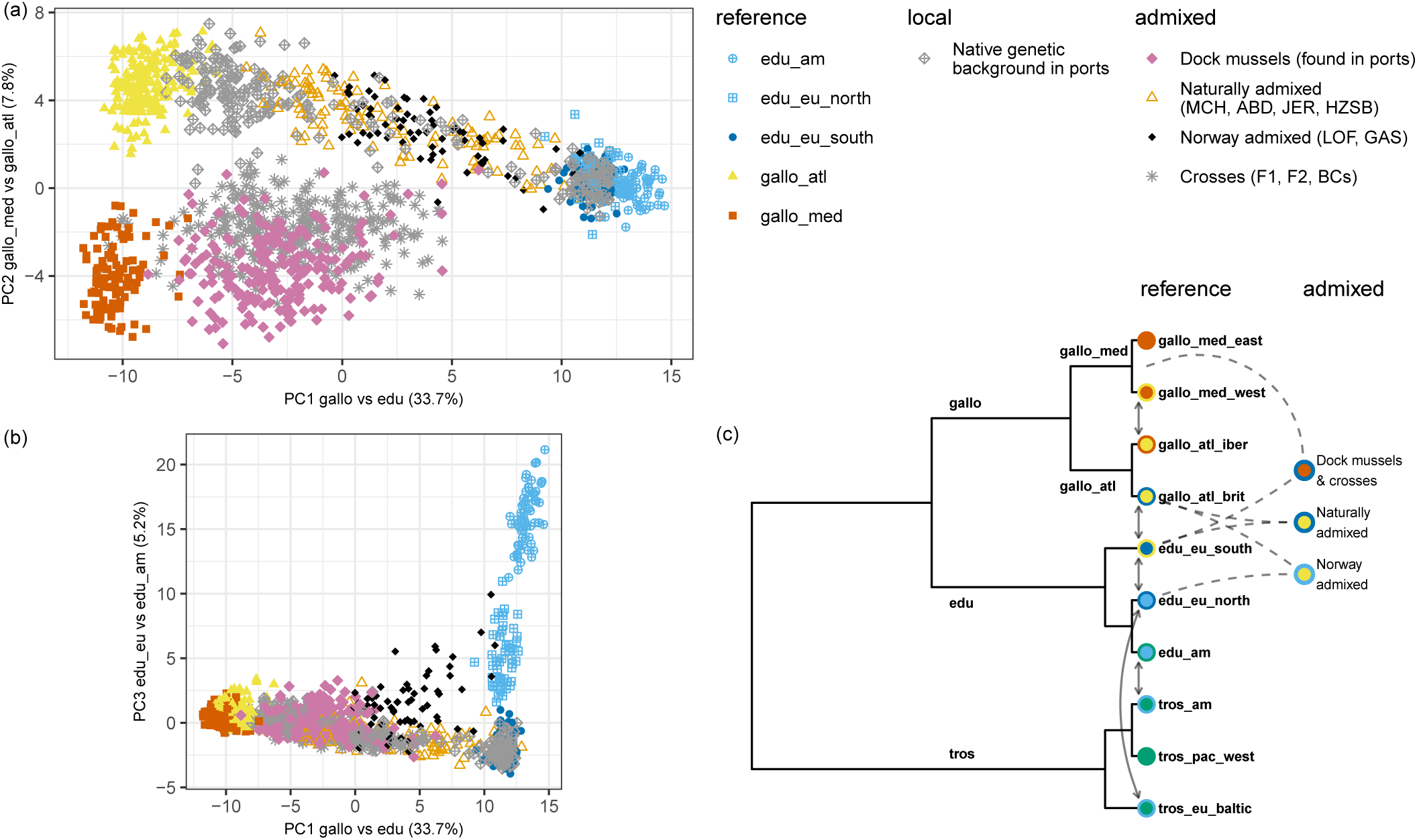
(a)-(b) Principal Component Analysis of reference samples and studied groups (*M. trossulus* samples were not considered). Locations in and around ports have been randomly sub-sampled for visual clarity (500 out of 1930 individuals retained) and individuals were classified as native genetic backgrounds (grey diamonds) or as dock mussels (pink diamonds) on the basis of a Structure analysis. The ports of interest are Le Havre, Cherbourg, Saint-Malo, Brest and Saint-Nazaire; see Figures 2 and 3 for details. (c) Schematic tree of lineage relationships presenting group names and colour schemes. External circle colours and arrows represent known local introgression between *Mytilus* spp. lineages. The three admixture types studied are presented in the right column.

**Figure 2:**
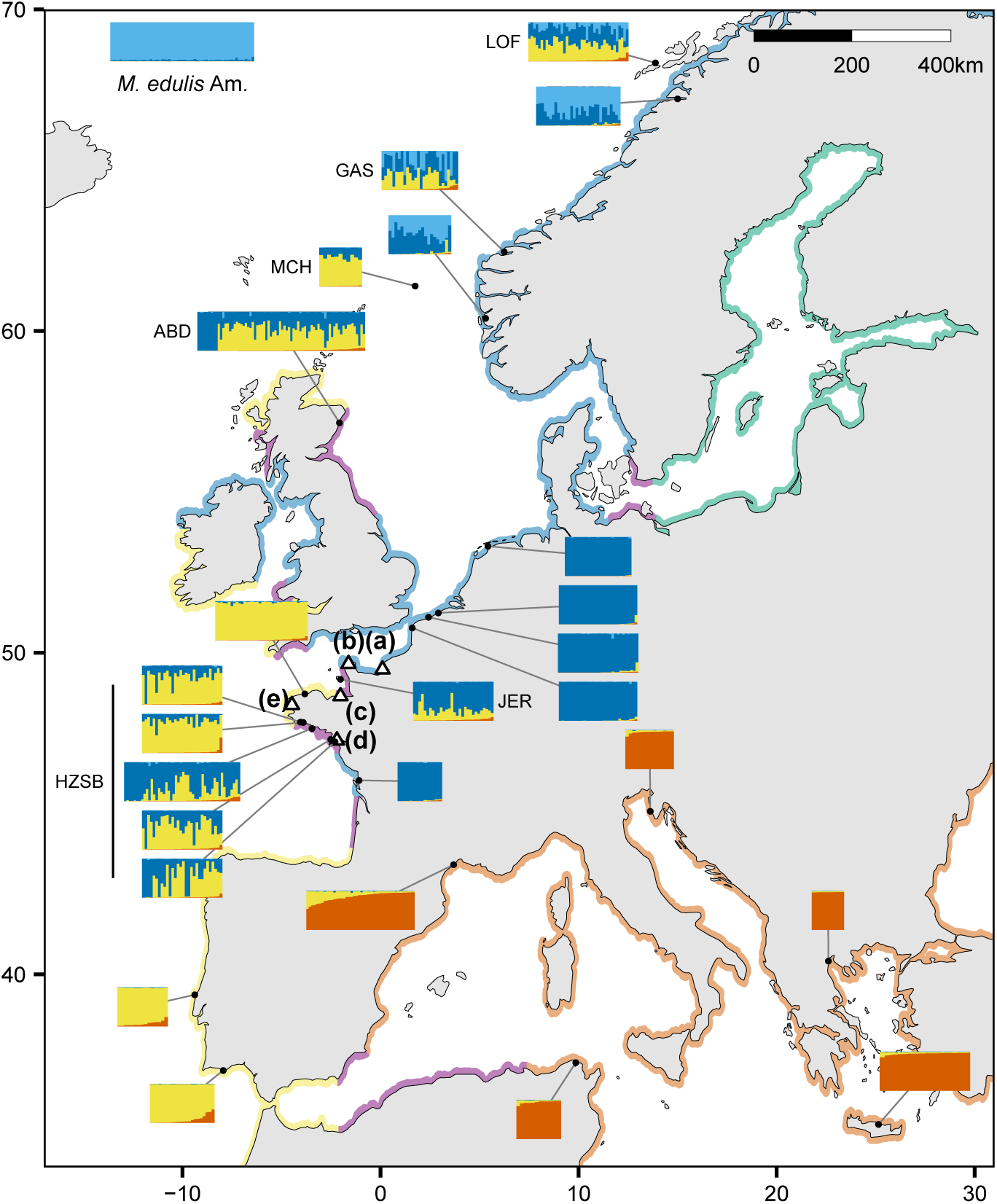
Location and ancestry composition of sites for reference and admixed populations. Barplots represent ancestries of individuals from the focal site, estimated by Structure with *K* = 4. In all barplots, individuals have been sorted from left to right by their level of Mediterranean *M. galloprovincialis* ancestry. Coloured coastlines indicate the approximate distribution of parental genetic background, with colour code as used in Figure 1. Hybrid zones are coloured in purple. Points (a)-(e) correspond to the ports of Le Havre, Cherbourg, Saint-Malo, Saint-Nazaire and Brest respectively, which are detailed in fig. 3.

To improve this predefined set of reference samples, an initial genetic clustering was performed with the software Admixture (Alexander, Novembre, and Lange, 2009, full nuclear dataset, 3 clusters, 30 replicates, fig S4) and the results were combined with the CLUMPAK software (Kopelman, Mayzel, Jakobsson, Rosenberg, & Mayrose, 2015). All individuals with less than 85% ancestry from their putative cluster were removed from the reference set (this threshold was chosen to account for local introgression in some populations). This step ensures there are no migrants, either from introduction or from sympatric species, and no hybrids in the reference panel.

Once the reference dataset was established, Hardy-Weinberg equilibrium (HWE) was estimated in each L3 level for all markers. edu_eu_south was separated in two groups, corresponding to the bay of Biscay (int, as in Fraïsse et al., 2016) and the English Channel (ext), for this analysis only, as they do not mate randomly but do not show significant genetic differentiation (Table S6). We used the hw.test function of the R package pegas (Paradis, 2010) with 10^4^ Monte Carlo permutations and a Benjamini-Yekutieli false discovery rate correction. Markers 604 and 190 were identified as significantly departing from HWE in at least one reference group (Figure S3).

### 2.3 Genetic map

Estimates of linkage between markers allow us to account for admixture linkage disequilibrium in ancestry estimation (see Structure analyses below), and to estimate time since admixture.

We used F2 crosses to produce a genetic map for a subset of markers analysed by Simon et al. (2018). This dataset comprises 97 markers genotyped for 110 reference *M. edulis* individuals, 24 reference Med. *M. galloprovincialis* individuals, 6 F1 parents (1 female, 5 males) and 132 F2 offspring. Markers that were not heterozygotic in all F1 parents, or with an allele frequency difference between species lower than 0.2 were removed to avoid spurious distortions and orientation. We also removed two markers with >10% missing data. This left a final dataset of 40 informative markers, and 114 F2 offspring. Alleles were oriented according to their frequencies in reference samples. We then used the R package qtl to produce a genetic map (Broman, Wu, Sen, & Churchill, 2003). Four additional markers were dropped by the internal checks in the package, for not passing the Mendelian segregation test in F2s (with Holm-Bonferroni correction). The final genetic map comprises 36 markers scattered among 16 linkage groups (Table S5). Only the first 8 linkage groups contain more than one marker.

An ‘unlinked’ set of markers was created by keeping the marker with the least missing data in each linkage group or physical contig. Markers not included in the linkage map analysis were considered to be unlinked. See Table S5 for a list of unlinked markers.

### 2.4 Population differentiation and genetic clustering

We aimed to identify known lineages of the *M. edulis* species complex to assign individual ancestry estimations and filter individuals based on their genetic compositions for downstream analyses.

Population differentiation analysis was used to assess the power of our set of ancestry-informative markers, and to test differences between admixed populations. Genetic clustering was then used to assign individuals to known lineages or to assess levels of admixture in the studied populations.

A principal component analysis (PCA) was performed in R, using the adegenet package (Jombart, 2008). The genotype data were centred and scaled, with the replacement of missing data by the mean allele frequencies. Any individuals identified as *M. trossulus* were removed from this analysis.

Hierarchical population differentiation tests were carried out with the R package hierfstat (Goudet, 2005). We used 10^4^ permutations for all tests. The Weir and Cockerham *F_ST_* estimator is reported when presenting population differentiation results. When calculating population differentiation between reference groups, markers with more than 30% missing data in *M. trossulus* populations were removed because of badly typed markers in this species (Table S3).

Ancestry estimation was performed with the Bayesian model implemented in the program Structure (Falush, Stephens, & Pritchard, 2003), which includes additional models of interest compared to the aforementioned Admixture software. Each result is composed of 25 replicates for each assessed number of genetic clusters, *K*, run for 8 *·* 10^4^ steps after a 2 *·* 10^4^ steps burn-in. The standard deviation for the *α* prior was set to 0.05 for better mixing of the chains. All analyses use uncorrelated allele frequencies (FREQSCORR = 0) and a separate and inferred *α* for each population (POPALPHAS = 1, INFERALPHA = 1, Wang, 2017). Replicates were merged with the program CLUMPAK (default parameters and MCL threshold set at 0.7) and the major clustering output of the most parsimonious *K* was used.

For Structure analyses, markers that departed from Hardy-Weinberg equilibrium in focal reference populations were removed to avoid departure from the algorithm model. The program was either run using the admixture model with linkage, using the F2 genetic map described above, or using a no-admixture model with the unlinked dataset (Table S5), as both models cannot be used simultaneously.

A first Structure analysis on the full dataset was used to remove all individuals with *M. trossulus* ancestry to focus on a ‘reduced dataset’ of *M. edulis* and *M. galloprovincialis*. Because, *M. trossulus* is present in sympatry in Norway and can hybridise with its congeners, a threshold of 10% ancestry was used to identify parental and most recent hybrid individuals (Table S3). From this reduced dataset, two analyses – with and without the admixture model – were performed (*K* in 3 to 6). Additionally, to allow a better classification of individuals at bay scales, Structure analyses were performed on a ‘local dataset’ with the ports and surrounding populations, with and without admixture, and without including the reference populations (*K* in 2 to 5). Finally, specific Structure runs with the linkage model were used to estimate the age of the admixture (cf. Supplementary information, section 1). Briefly, admixture linkage disequilibrium allows the estimation of the number of breakpoints per Morgan since the admixture event, *r*, which can be interpreted as an estimate of the number of generations since a single admixture event (Falush et al., 2003).

Mussels from the admixed populations with Atl. *M. galloprovincialis* (introduced and natural) were classified using the reduced dataset without admixture, using the yellow and grey clusters corresponding to pure Atl. *M. galloprovincialis* and admixed *M. galloprovincialis* respectively (*K* = 5, Figure S19). To obtain a finer classification in port areas, mussels were assigned to *M. edulis*, Atl. *M. galloprovincialis* or dock mussel clusters using the local Structure analysis without admixture (*K* = 3, Figure S20). See Table S3 for details on the selection thresholds for each group and Figure S21 for independent plots of selected individuals.

The software Newhybrids (Anderson & Thompson, 2002) was used to evaluate the probability that individuals were first or second generation hybrids between the dock mussels and native lineages (Figures S26-S27).

### 2.5 Comparison of ancestry levels

To investigate the similarities and differences in the ancestry compositions of samples from different admixture events and localities (Table 1), we formally tested for variation in ancestry levels.

Independent comparisons were used for admixtures implicating Med. and Atl. *M. galloprovincialis*. For each population of interest, admixed individuals (identified as described in the previous section) were selected and native individuals were removed. The Structure ancestry estimates with admixture, identifying the four clusters edu_eu_south, gallo_atl, gallo_med and edu_am, were used (*K* = 4, Figure S21). This selection allowed a homogeneous comparison of ancestry levels between all admixed populations (Figure S23).

A non-parametric Kruskal-Wallis one-way ANOVA was used to test the statistical difference of the four ancestry values (*Q*) between populations of each admixture type. Additionally, a non-parametric post-hoc pairwise comparisons test was carried out, using the Dwass-Steel-Crichtlow-Fligner test (Critchlow & Fligner, 1991; Hollander, Wolfe, & Chicken, 2015). We applied Benjamini-Yekutieli corrections for multiple testing.

To test the hypothesis of increased introgression of Med. *M. galloprovincialis* ancestry coming from dock mussels into Atl. *M. galloprovincialis* in the Bay of Brest, native Atl. *M. galloprovincialis* groups from Brittany were identified and their ancestries were compared: (i) mussels distant from the Bay of Brest, Northern Brittany population (gallo_atl_brit); (ii) individuals outside the Bay of Brest (the limit being the entrance straight), taken as reference local individuals; and (iii) individuals inside the Bay of Brest classified as Atl. *M. galloprovincialis* with the local Structure without admixture result (Figure S20).

### 2.6 Least cost distance analyses and Geographic cline fitting

To visualise transitions at the port entrance at the locus level, we fitted clines of allele frequencies along a spatial axis. The objective is to assess the concordance of transitions among markers and with the observed global ancestry.

As a proxy for connectivity between sampling sites, least cost path distance matrices were produced for each port and took into account obstacles such as land and human made barriers (e.g., breakwaters and seawalls). A raster of costs was built for each port from polygon shapefiles (‘Trait de côte Histolitt Métropole et Corse V2’, produced by SHOM and IGN) modified to include small port structures that could stop larval dispersal or to exclude inaccessible parts. Locks inside ports were considered as opened for the purposes of distance calculation between isolated points. We used the program QGIS to handle polygons and raster creation. Land was coded as missing data and water was set to have a conductance of one. The R package gdistance was used to compute transition matrices based on those cost rasters and to compute least cost distances between points for each dataset (van Etten, 2017).

Geographic clines per SNP were fitted for each port (excluding Saint-Malo which only had one port sample) with the R package hzar (Derryberry, Derryberry, Maley, & Brumfield, 2014). The port of Le Havre was divided into two independent transects: North and South corresponding to the historic basins and the ‘Port 2000’ recent installations respectively. The least cost distance from the most inward site in each port (indicated by a triangle in Figure 3) was taken as a proxy for geographic distance and to project geographic relationships on a single axis. For the Bay of Brest, the starting site was taken as the right-most population in Figure 1g, up the Élorn estuary. The three points in the bottom-right corner of Figure 3e containing Med. *M. galloprovincialis* ancestry were excluded from the fit, to account for discrepancies between least cost path distances and the presence of the dock mussels. Pure *M. edulis* individuals were removed for the analysis in the bay of Brest and Atl. *M. galloprovincialis* individuals for the ports of Le Havre, Saint-Nazaire and Cherbourg. Clines were fitted using a free scaling for minimum and maximum frequency values and independence of the two tails parameters. We used a burn-in of 10^4^ and a chain length of 10^5^ for the MCMC parameter fit. Only differentiated loci are presented in Figure 4 (panels a-d: allele frequency difference (AFD) > 0.5, panel e: AFD > 0.3; see Figures S28-S32 for details).

**Figure 3:**
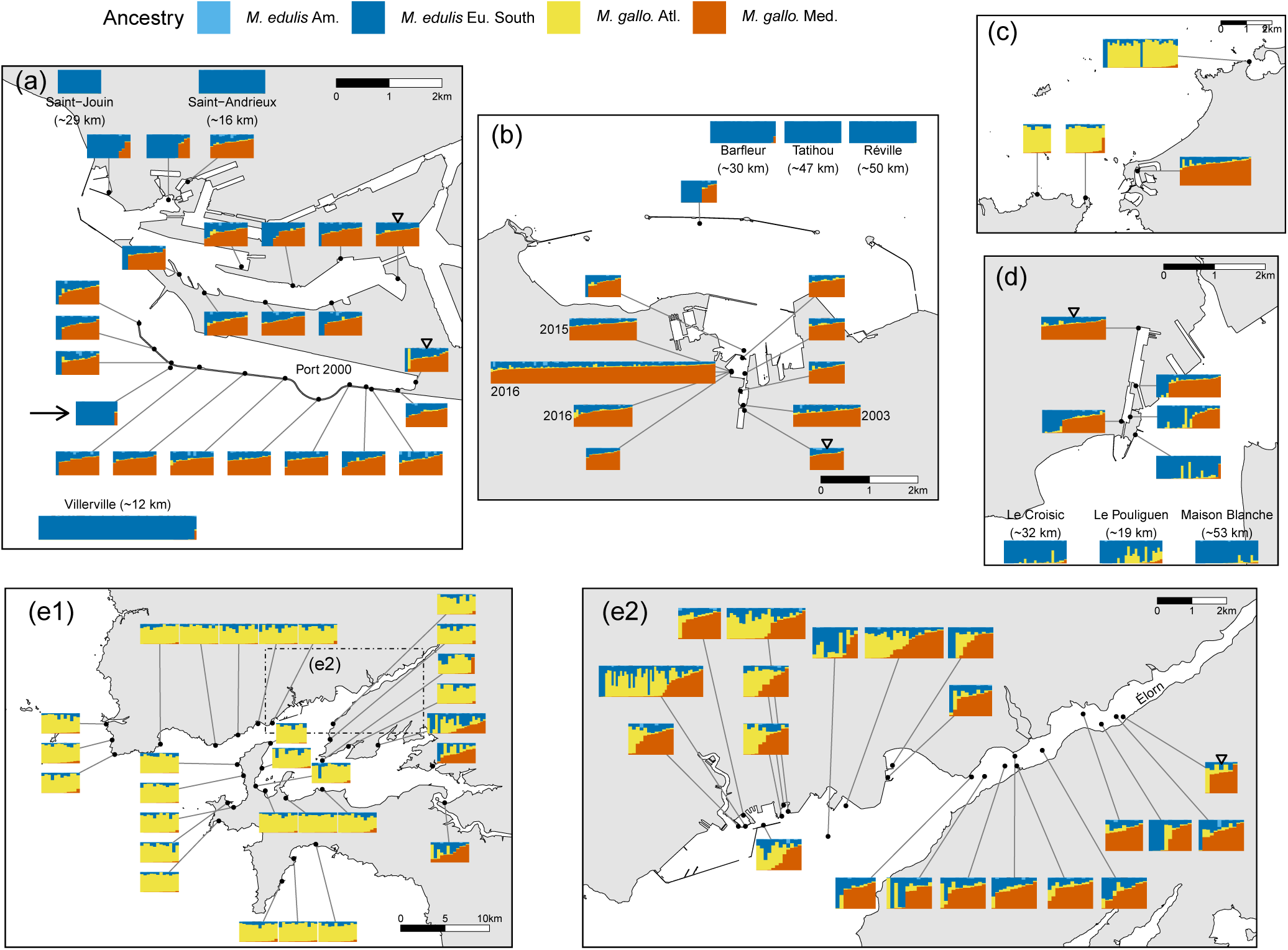
Ancestry composition of sites for each port. As in Figure 2, barplots represent the ancestry estimation for individuals at the indicated locations and are ordered from left to right by their Med. *M. galloprovincialis* ancestry. Barplots at the map edges correspond to distant populations with the least cost path distance from the port indicated in parentheses. The inner-most populations used to fit geographic clines are indicated by the reversed triangles. (a) Le Havre; note that the two distinct main basins (North and South-Port 2000) found in this port were separated for geographic cline analyses; the arrow indicates a site located on the estuary side of the dyke, characterised by a majority of *M. edulis* individuals. (b) Cherbourg; dates indicate collection year; all other samples were collected in 2017. (c) Saint-Malo. (d) Saint-Nazaire. (e1) Bay of Brest. (e2) Detailed map of the port of Brest and the Élorn estuary, which corresponds to the inset rectangle in panel (e1).

**Figure 4:**
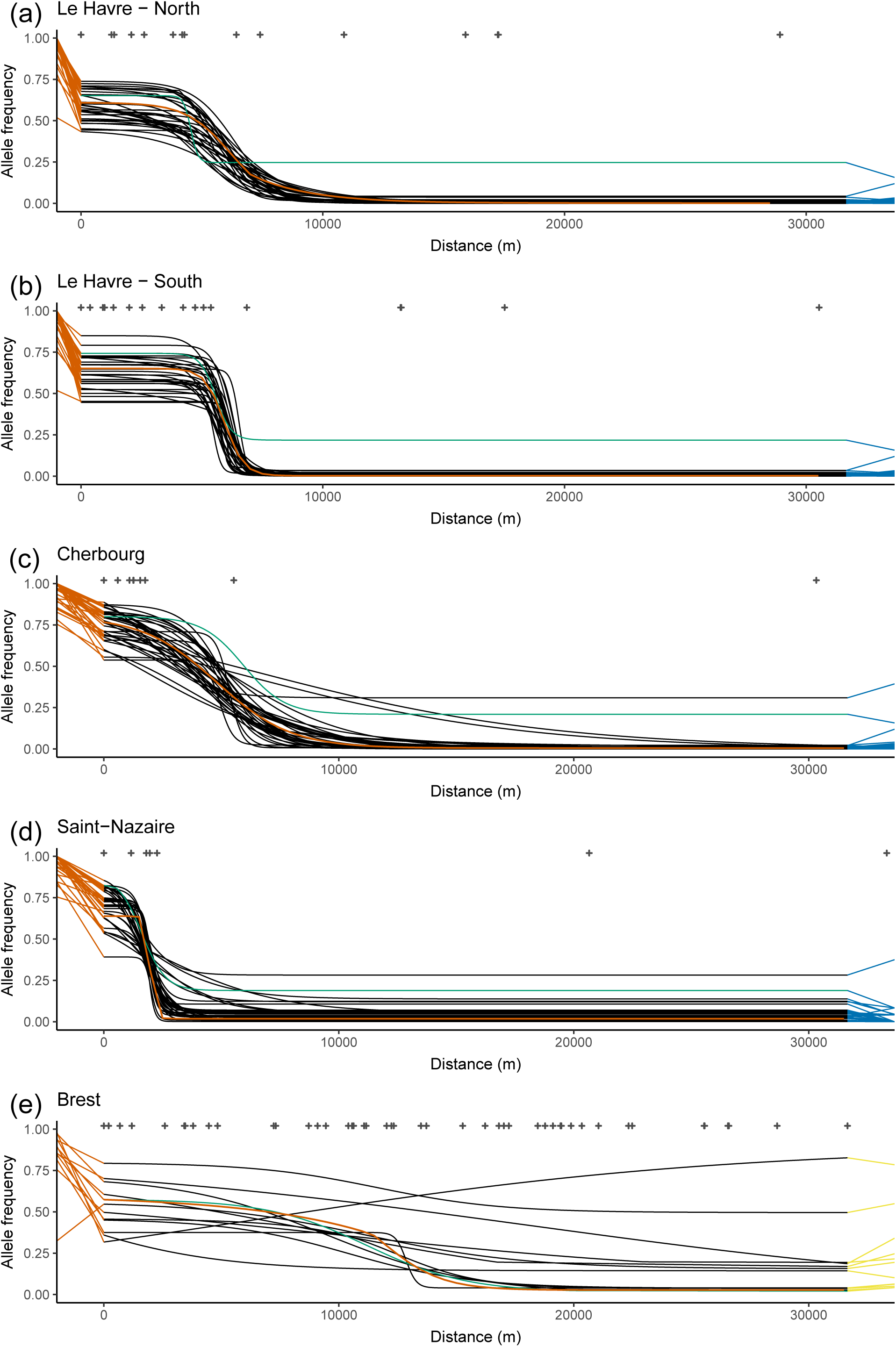
Geographic clines computed with the package hzar in each study ports (except St-Malo, see text). The x-axis is the distance from the most inward point (reversed triangles in figure 3) determined by a least-cost path analysis. Top crosses indicate the distance of each site considered. For representation purposes, some distant points are not displayed, but were used in the cline fit. Only alleles with a frequency difference of 0.5 between left-most port population and sea-side reference are presented (except for panel (e) where the threshold is 0.3), each with a distinct black line. For each marker, left and right segments join the frequency fitted at the end of the cline to the frequency observed in reference populations, with Med. *M. galloprovincialis* in orange and South-Eu. *M. edulis* in blue (or Atl. *M. galloprovincialis* in yellow). For (a)-(d), references are Mediterranean *M. galloprovincialis* on the left and *M. edulis* on the right. For (e), the right hand side reference is the local Atlantic *M. galloprovincialis*. The orange cline is the mean cline computed from the Mediterranean *M. galloprovincialis* Q-value from Structure, in mean proportion of ancestry. The cline of the female mitochondrial marker (601) is shown in green. (a) Le Havre, North transect (historic basin). (b) Le Havre, South transect (Port 2000). (c) Cherbourg. (d) Saint-Nazaire. (e) Bay of Brest.

### 2.7 Distortions from expected frequencies and correlations

Our data include multiple admixture events. To ask if outcomes were similar across events, we compared the deviations of marker allele frequencies from their expected values in each situation.

We denote the expected frequency of an allele in an admixed focal population as *f_exp_*. This expected value is calculated from the observed allele frequencies in pure-lineage reference populations, and from the mean ancestry values across all markers for the focal population, as estimated from Structure.

Admixed population frequencies are calculated only with admixed individuals in each population (see section 2.5 for details and Figure S21 for selected individuals). We used the results of ancestry estimation from Structure with *K* = 4 clusters (edu_eu_south, gallo_atl, gallo_med, and edu_am) and summed ancestries from South-Eu. and American *M. edulis*, giving the composite ancestry estimation *Q_edu_* for each individual:

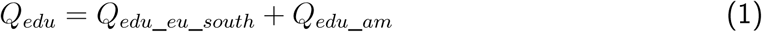

In particular, with three reference populations, the expected allele frequency is:

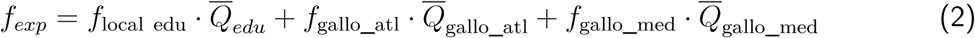

Here, *f* values denote the allele frequencies in the reference population indicated by the subscript, and the *Q*-values denote the mean ancestry from the focal admixed population. gallo_med and gallo_atl correspond to the L2 level encompassing lower population classifications (fig 1c and Table S4) as the precise origin of the parental populations are not known below this level.

For lab crosses, the parental Med. *M. galloprovincialis* L3 level is known and corresponds to gallo_med_west. Therefore its frequency was used in place of *f*_gallo_med_. For dock mussels the ‘local edu’ lineage is taken to be the South-Eu. *M. edulis* one (edu_eu_south). For LOF and GAS admixed populations, we used the North-Eu. *M. edulis* lineage (edu_eu_north) to estimate parental allele frequencies (*f*_local edu_) while using the usual *Q̄*_edu_ estimation.

The deviation of the observe frequency *f_obs_* from the expected frequency *f_exp_* is defined as:

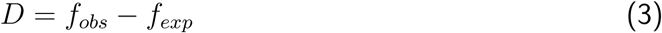

This computation allows us to estimate a distortion by locus from the average genomic expectation given the population ancestry and parental allele frequencies. The correlation of distortions by locus are then computed between admixed populations, corresponding to different admixture events (e.g. between one dock mussel and one Norway admixed population). For each correlation, we used Pearson’s *r* to estimate the strength of the correlation and tested the significance with a permutation test (5 *·* 10^4^ permutations). The classic t-test was not used due to the distortions not following normality.

When multiple correlations pertained to the same null hypothesis (e.g. that distortions in lab backcrosses do not correlate with distortion in ports), and datasets contained possible non-independence (e.g., from migration of hybrids between ports), we used a modified Fisher’s method to combine *p* values, developed by Poole, Gibbs, Shmulevich, Bernard, and Knijnenburg (2016) and implemented in the R package EmpiricalBrownsMethod.

## 3 Results

### 3.1 Differentiation between lineages and characterisation of admixed populations

We collected or reanalysed samples from several locations, with known or suspected admixture between different species or lineages of *Mytilus* mussels (Figure 1c, Table 1).

We first verified that our dataset could distinguish between species and focal lineages. Hierarchical genetic differentiation tests based on putative reference groups (Figure 1c, Table S4) showed significant *F_ST_* distances until the grouping level L3. *F_ST_* ranges between 0.72 and 0.81 at the species level (L1), between 0.38 and 0.48 for L2 levels within species and between 0.0024 and 0.31 for L3 levels within L2 (see Table S6 for details; note that our SNP panel is enriched for ancestry-informative SNPs and so these values should not be interpreted as genome-wide averages).

Initial PCA and Structure analyses identified the presence of all three *Mytilus* species. However, *M. trossulus* was present in only a few populations (i.e. Norway, North Sea), consistent with previous knowledge of its range (Figure S5). Because *M. trossulus* is not centrally relevant to the present work, individuals with more than 10% *M. trossulus* ancestry were removed from subsequent analyses.

After removing *M. trossulus* individuals, both the PCA (Figure 1a-b) and the Structure Bayesian clustering (*K* = 4, Figures S6-S15) show a clear differentiation between the parental lineages (edu_am, edu_eu_south, gallo_atl and gallo_med). Both methods also allow us to identify and further characterise three characteristic patterns of admixture in our data, which we called ‘naturally admixed’, ‘Norway admixed’ and ‘dock mussels’. We describe each of these in detail below.

Each admixed pattern was further investigated by comparing ancestry estimations of populations to characterise the variation between locations (Structure *Q*-values, *K* = 4, Figure S23).

### 3.2 Natural hybridisation

Several samples are the result of natural admixture between Atl. *M. galloprovincialis* and South-Eu. *M. edulis* and are called ‘naturally admixed’ (Figure 1c, Table 1). This category includes geographically distant samples from Scotland (ABD), the English Channel island of Jersey (JER), the Murchison oil platform in the Norwegian Sea (MCH) and the natural hybrid zone in South Brittany (HZSB, Figure 2). As far as we know, these groups are free from human-mediated introductions.

Naturally admixed populations cover much of the range of admixture proportions observed between the two parental species (Figure S23). These four populations exhibit significant differences in their Atl. *M. galloprovincialis* ancestry, with the exception of the MCH/HZSB comparison (Table S10). JER is the most *M. edulis*-like population while MCH and ABD are the most *M. galloprovincialis*-like, with HZSB being the most variable one. Interestingly, JER exhibit a homogeneous excess of South-Eu. *M. edulis* ancestry, contrasting with the Atl. *M. galloprovincialis* ancestry excess of the three other natural populations (Figures 2 and S23). Atl. *M. galloprovincialis* ancestry excess is usually observed in contact zones reflecting the asymmetric introgression with South-Eu. *M. edulis* (Fraïsse et al., 2016).

### 3.3 Admixed populations in Norway

We named a second admixture pattern ‘Norway admixed’, because it includes two Norwegian populations (LOF, GAS). These admixed mussels involve Atl. *M. galloprovincialis* and North-Eu. *M. edulis* (Figure 1b), and are defined as non-indigenous (Mathiesen et al., 2016). LOF and GAS do not differ significantly at any of the four different ancestry estimates (Table S10).

These admixed mussels are on average composed of 40% Eu. *M. edulis* (*SD* = 15.82, *N* = 63), 16% American *M. edulis* (*SD* = 15.35), 41% Atl. *M. galloprovincialis* (*SD* = 13.91), and 3% Med. *M. galloprovincialis* (*SD* = 3.83) (Figures S21 and S23). The presence of individuals with some Atl. *M. galloprovincialis* ancestry was also confirmed in Svalbard (Figure S14; Mathiesen et al., 2016). On average, admixed mussels in Svalbard have lower proportions of Atl. *M. galloprovincialis* ancestry. These individuals were not used in downstream analyses, due to their small number.

Norway admixed populations were also compared to naturally admixed populations given they both involve the Atl. *M. galloprovincialis* lineage. Nearly all pairwise comparisons of the Atl. *M. galloprovincialis* ancestry are significantly different, with the exception of the GAS/JER comparison (Table S10). GAS and LOF appear to be more similar to JER, with an excess of *M. edulis* ancestry, than they are to the other three naturally admixed populations.

### 3.4 Dock mussels

#### 3.4.1 An admixture between geographically distant lineages

We identified a group that we labelled ‘dock mussels’, found in five French ports, and more rarely in their vicinity. They exhibit a characteristic admixture between Med. *M. galloprovincialis* and South-Eu. *M. edulis*, and are defined as the intermediate cluster between these two lineages (Figure 1, Table. 1). The selection of individuals defined as dock mussels is based on a Structure analysis without admixture (Figure. S20). Dock mussels are closer to Med. *M. galloprovincialis* than to *M. edulis* in the PCA, reflecting the estimated ancestries, and are not differentiated by other axes of the PCA (Figure 1a). Additionally, they show a large variance in all directions, presumably including inter-specific hybrids with *M. edulis* and inter-lineage hybrids with Atl. *M. galloprovincialis*. It is noteworthy that apart from the dock mussels, and the lab crosses between Med. *M. galloprovincialis* and South-Eu. *M. edulis*, no other population clusters in this region of the PCA (i.e. intermediate placement between Med. *M. galloprovincialis* and South-Eu. *M. edulis*). This implies that no natural hybridisation is observed between these two lineages in our dataset. This is in accordance with the distribution of the *Mytilus* lineages (Figure 2).

We analysed three other large ports to search for dock mussels, but none showed the presence of this class of mussels: La Rochelle (France, Figure S16), Bilbao (Spain, Figure S17) and New York city (USA, Figure S18).

In the five colonised ports, individuals of native parental genetic backgrounds are found in addition to dock mussels (Figures 1a-b and 3). These native mussels are (i) Pure South-Eu. *M. edulis* around Cherbourg, Le Havre and Saint-Nazaire, and (ii) Pure Atl. *M. galloprovincialis* from Brittany around Brest, Saint-Malo and Saint-Nazaire. We also observed intermediate individuals between Atl. *M. galloprovincialis* and *M. edulis* corresponding to admixed individuals or hybrids in the Bay of Brest area, Saint-Nazaire and Saint-Malo. All of these locations are in or close to natural hybrid zones between those two species, while the aquaculture of *M. edulis* in the Bay of Brest, imported from the Bay of Biscay, is an additional source of *M. edulis* in this area, especially since dispersing larvae from aquaculture sites are common (for details see Figure S11).

In term of estimated ancestries (Structure *Q*-values), dock mussels are on average composed of 25% Eu. *M. edulis* (*SD* = 11.17, *N* = 879), 69% Med. *M. galloprovincialis* (*SD* = 11.85), 4% Atl. *M. galloprovincialis* (*SD* = 6.08) and 2% American *M. edulis* (*SD* = 3.04) (Figure S21). Allele frequencies of dock mussels for markers differentiated between *M. edulis* and Med. *M. galloprovincialis* are also consistent with the observed levels of admixture, and are strongly concordant between markers (Figure S22). All port populations are highly similar, both spatially and temporally, in their variance of allele frequencies regardless of their overall level of introgression (Figure S22).

When comparing ports, Cherbourg, Saint-Nazaire and Saint-Malo are the least introgressed populations (Figure S23, Table S11). Le Havre appear to be the most introgressed by South-Eu. *M. edulis*. Brest also have reduced levels of Med. *M. galloprovincialis* ancestry, equivalent to what is found in Le Havre, but due to an excess of Atl. *M. galloprovincialis* ancestry. Cherbourg, Saint-Malo and Saint-Nazaire do not differ significantly in South-Eu. *M. edulis*, Atl. and Med. *M. galloprovincialis* ancestries, despite the fact they are in different native species contexts.

For the port of Cherbourg, we were able to analyse several temporal samples between 2003 and 2017 (Figure 3b). These exhibit a small differentiation between the 2003 sample and later years (2015 and 2016; *F_ST_* = 0.0066 and 0.0097, Table S8) and this seems to be driven by a small increase in Med. *M. galloprovincialis* ancestry in 2015 and 2016 (significant only between 2003 and 2016, Table S12). The only other historical sample in our collection was a site in the Bay of Brest that showed the absence of dock mussels in 1997 (Pointe de L’Armorique, PtArm97, fig S11). However, this area also exhibited only one dock mussel genotype 20 years later (Brest-24).

#### 3.4.2 Dating the admixture of dock mussels

To estimate the age of the admixture event which resulted in the dock mussels, we inferred levels of linkage disequilibria (Figure S24). Disequilibria were present, but at low levels indicating that there had been several generations of recombination since admixture. We computed a linkage map from the lab produced F2, and found that it was consistent with the disequilibria present in the dock mussels. Using this map, and the linkage option in the Structure package, we estimated the admixture time to be between 4 and 14 generations, depending on the port (Table S14 and supplementary methods).

As survival and lifetime are highly variable and environment dependent in mussels, it is difficult to translate these estimates into clock time. However, given that mussels reach maturity at ∼1 year and have a high early life mortality rate, 1-2 years seems a reasonable estimate of the generation time, dating the admixtures at between 4 and 28 years ago. We note that our oldest sample from Cherbourg in 2003, provides one of the oldest estimates, and so could not be used to calibrate a ‘recombination clock’.

#### 3.4.3 Dock mussels are spatially restricted to ports

The individual ancestries were plotted spatially to assess their distribution in and around the five studied French ports (Figures 3).

The ports of interest are localised in regions characterised by different native species (Figure 2). The native species around Le Havre and Cherbourg is South-Eu. *M. edulis* while in the Bay of Brest, the native mussels are Atl. *M. galloprovincialis* (Figure 3). Saint-Malo and Saint-Nazaire lie on the limits of hybrid zones between *M. edulis* and *M. galloprovincialis*. However, surroundings of Saint-Malo are mostly inhabited by Atl. *M. galloprovincialis* (Figure 3c), and Saint-Nazaire is located in a zone mostly composed of *M. edulis* with the presence of Atl. *M. galloprovincialis* in sympatry (Figure 2 and 3d). Around the latter, local *M. galloprovincialis* are more introgressed by *M. edulis* than those found in Brittany as they lie at the far end of the South Brittany hybrid zone (Bierne et al., 2003).

Four of the five studied ports (all except Brest) have locked basins where the dock mussels were found. Importantly, dock mussels are nearly all localised inside port infrastructures, and we observed a sharp shift at the port entrance (Figure 3). For the ports of Saint-Nazaire, Saint-Malo, Cherbourg and Le Havre only four individuals with Med. *M. galloprovincialis* ancestry were detected in coastal wild populations (out of 341 individuals presented in Figure 3). Those individuals were observed at distances between a few hundred meters to 30 km from the entrance of the ports.

In the opposite direction (from the natural coast to the port), we mainly find native migrants close to the port entrance inside Le Havre, Cherbourg and Saint-Nazaire (Figure 3). Le Havre and Saint-Nazaire are the ports containing the largest number of *M. edulis* migrants, yet Le Havre is the only one where F1 hybrids between dock mussels and *M. edulis* have been observed (identified with Newhybrids, Figure S26).

The Bay of Brest is of particular interest for two reasons (Figure 3e1-e2). First, the local background is the Atl. *M. galloprovincialis* lineage, contrasting with the other ports where the native background is *M. edulis* (with the exception of Saint-Malo), and exhibiting higher sympatry inside port infrastructure than anywhere else. Second, mussels with a typical dock mussel admixed genetic background have been detected outside port infrastructures, which motivated an extensive sampling. Contrary to the other ports, dock mussels extensively colonised the local environment, mainly inside and close to estuarine areas.

Dock mussels are, however, restricted to the inside of the bay with no detectable influence on external *M. galloprovincialis* populations. We compared several groups of Atl. *M. galloprovincialis* from Brittany (away, close and inside the Bay of Brest) to assess the potential introgression from dock mussels to the local populations. While levels of *M. edulis* ancestry increased and levels of Atl. *M. galloprovincialis* decreased significantly from distant populations to inside the Bay of Brest, the levels of Med. *M. galloprovincialis* ancestry did not differ significantly (Table S13). Nonetheless, we note that the tail of the distribution of Med. *M. galloprovincialis* ancestry in the Bay of Brest is skewed towards higher values (Figure S23). This tail is due to the presence of hybrids between dock mussels and the local native Atl. *M. galloprovincialis* (Figure S27).

#### 3.4.4 Geographic clines show sharp and concordant transitions at the port entrance

Allele frequencies shift sharply at the entrance of ports (Figure 4a-d) and clines are highly concordant both between markers and with the mean ancestry cline (red line). Compared to the reference Med. *M. galloprovincialis* frequencies, dock mussels show a global decrease of allele frequency due to a genome wide introgression from the local species.

Clines have narrow widths across all ports. Average widths are 3.99 km (*SD* = 1.80) and 1.30 km (*SD* = 0.52) for the North and South transects of Le Havre respectively (Figure 4a-d); 7.37 km (*SD* = 5.38) in Cherbourg (Figure 4c); 2.16 km (*SD* = 2.15) in Saint-Nazaire (Figure 4c), and 18.51 km (*SD* = 14.03) in the Bay of Brest (Figure 4e).

The difference between the North and South transects in Le Havre is best explained by the presence of more *M. edulis* or hybrid individuals at the entry of the North basin (Figure 3a). The interpretation in the Bay of Brest is more difficult due to two factors. First, the spread of dock mussels and sympatry with local ones in several populations make allele frequencies more variable between close populations (Figure 3e-f). Second, we had a reduced number of differentiated markers between Atl. and Med. *M. galloprovincialis* in our dataset with lower level of differentiation.

### 3.5 Repeatability of allele frequency deviations between admixture events

If admixture events are non-independent (e.g., due to migration between ports), or if admixture events are independent, but lead to repeatable patterns of natural selection, then we would expect to see the same alleles over- or under-represented in different locations.

We cannot compare allele frequencies directly, because different locations are characterised by different overall levels of ancestry. Therefore, for each marker, in each location, we calculated its deviation from expected values. These expected frequencies were calculated from the allele’s frequencies in the reference parental populations, combined with the overall levels of ancestry in the sampled location (this is Barton’s concordance analysis, eq. 1-3).

Examination of these allele frequency deviations showed some suggestive similarities between admixture events. For example, the mitochondrial marker (601) is differentiated between the Med. and the Atl. *M. galloprovincialis* lineages (Figure S38). This locus exhibits large distortions (*D*) towards the Med. *M. galloprovincialis* lineage in Le Havre, Cherbourg and Saint-Nazaire (0.11, 0.16, 0.13 respectively), while displaying smaller distortions in Brest and Saint-Malo (0.03 in both cases).

More formally, the repeatability of admixture events can be assessed by correlating the complete set of allele-frequency deviations between events. Four types of comparisons corresponding to differences in implicated lineages are presented in Figure 5.

**Figure 5:**
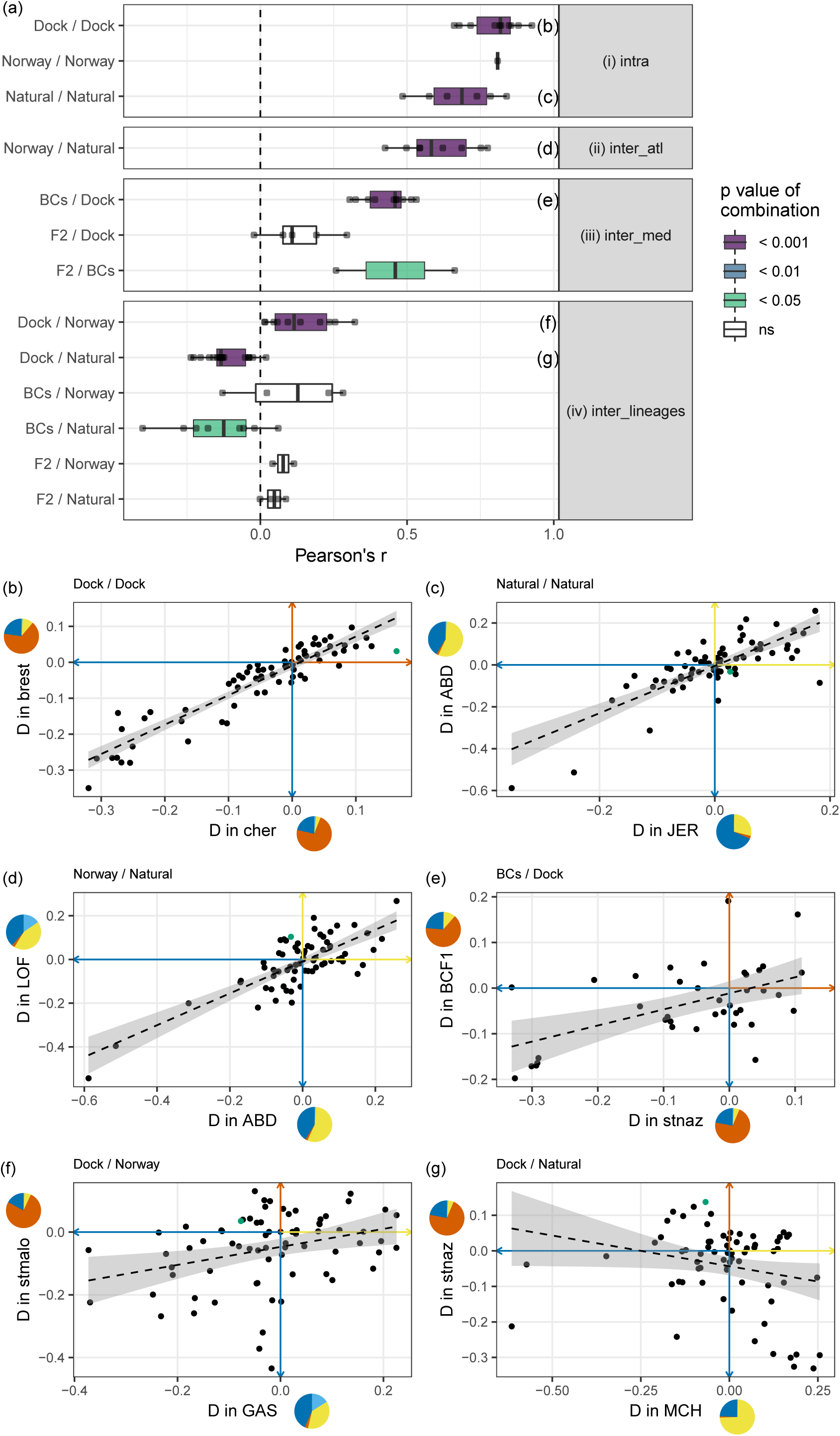
(a) Pearson’s *r* correlation coefficients of distortions (*D*) between groups of admixture types. The admixture types are: dock mussels (Dock), Norway admixed (Norway), naturally admixed (Natural) or crosses (BCs and F2). Each grey dot is a correlation between two sites (e.g. havre *vs.* cher is one of the point shown in the Dock/Dock row or BCF1 *vs.* MCH in the BCs/Norway row). The significance level correspond to the combination of *p* values among comparisons (see methods). Four types of comparisons were tested: (i) intra - comparisons among the same types of admixture events; (ii) inter_atl - comparisons of the admixture events between Atl. *M. galloprovincialis* and *M. edulis*; (iii) inter_med - comparisons of the admixture events involving South-Eu. *M. edulis* and Med. *M. galloprovincialis*; (iv) inter_lineages - comparisons of admixture events between different backgrounds. Panels (b)-(g) at the bottom show examples of correlations between distortions computed in two locations, for the highest significance levels per type comparisons (purple colour in panel [a]). All correlations presented are significant and linear models with 95% confidence intervals are plotted. The colour of the axis shows the direction of the distortion in term of lineage, using the colour code shown in Figure 1. Pies show the mean ancestry composition of the population considered. Distortion corresponding to the mitochondrial marker (601) is highlighted in green in panels (b)-(g)

We examined all pairwise comparisons involving the same parental backgrounds in similar conditions (Figure 5a-[i]): the five dock mussels populations from French ports (‘Dock / Dock’), the two Norwegian introductions (‘Norway / Norway’), and the natural hybrid zones (‘Natural / Natural’). In each case, the allele frequency deviations are significantly and positively correlated between events, with large to medium effect sizes (Figures 5 and S33-S34). The same was also true when we compared the Norwegian introductions to the natural hybrid zones involving the same *M. galloprovincialis* genetic background (‘Norway / Natural’, Figure 5a-[ii]). Remarkably, strong correlations were also observed when we compared dock mussels to lab crosses involving the same lineages (Figure 5a-[iii]). The correlations were strongest for lab backcrosses (BCs), and much weaker and non-significant for the F2. This is consistent with the genetic makeup of the dock mussels, which have hybrid indexes closer to BC genotypes than to F2s (fig S23 and S25), albeit more recombined.

Globally, the level and consistency of correlations increases with the similarity between admixture events (from groups [iv] to [i] in Figure 5). Panels (i)-(iii) indicate that admixture events of different kinds can lead to strongly repeatable results. But this is only true when the same genetic backgrounds are involved. To show this, Figure 5a-(iv) shows results from pairs of admixture events involving different backgrounds (e.g. Dock vs. Norway admixture). In this case, effect sizes are small to medium, and sometimes negative.

### 3.6 Additional putative anthropogenic introductions

While the overall genetic composition of many of our sampled populations was as expected, we also obtained some isolated but unexpected results which we report in the following section. First, the port of New York showed higher levels of South-Eu. *M. edulis* ancestry, up to 30%, compared to other populations from Long Island Sound (Figure S18). Therefore we cannot exclude the possibility that there has been an introduction of Eu. *M. edulis* in or close to the port of New York.

Second, outside of ports, multiple long distance migrants from different origins were identified. The reanalysis of the Coolen (2017) samples did not show any pure *M. galloprovincialis* individuals (Figure S13). However, one population contained six individuals composed of 10 to 30% Med. *M. galloprovincialis* ancestry (Q13A, Figure S13). This population is located offshore, at around 25 km from the entry of the port of Rotterdam, which is the largest commercial port of Europe. Given the greater proportion of migrants at this distance, as compared to results from other ports, the presence of dock mussels in Rotterdam is highly probable and will require further investigation.

Similarly, one population in the bay of Biscay, on the Atl. coast of Oléron Island (ROC_VER), contained an individual with pure Med. *M. galloprovincialis* ancestry and a few individuals with some levels of Med. *M. galloprovincialis* ancestry in an Atl. *M. galloprovincialis* background. Those latter individuals might plausibly be migrants Atl. *M. galloprovincialis* from the Basque Country. Indeed, unlike populations from Brittany, Iberian Atl. *M. galloprovincialis* populations south of the last hybrid zone with *M. edulis*, have low to medium levels of Med. *M. galloprovincialis* ancestry due to their contact with this lineage in the South (see Bilbao port samples, Figure S17 and classification as Atl. *M. galloprovincialis* in Figure S19).

Other unexpected ancestries were observed in other locations. For example, we found at least one Atl. *M. galloprovincialis* in the port of Le Havre (LeHa_P11, Figure S8). We also report here the presence of an F1 hybrid between *M. edulis* and Atl. *M. galloprovincialis* in the port of Sète (France, Mediterranean coast) despite the fact that neither of these lineages are found in this area. We also analysed two samples from a ferry hull collected in 2011 and 2013. The ferry crosses the English Channel between a *M. galloprovincialis* region in Brittany (Roscoff) and a hybrid zone in the UK (Plymouth) where *M. edulis* and *M. galloprovincialis* are found in sympatry (Hilbish, Carson, Plante, Weaver, & Gilg, 2002, and personal communication). Both samples showed a mixture of *M. edulis* and Atl. *M. galloprovincialis* individuals (Figure S15, Fer11 and Fer13), highlighting once again the role of ship traffic in the displacement of species and their role as meeting points where hybridisation can occur.

We also detected a signature of Atl. *M. galloprovincialis* in the northern English Channel, and southern North Sea, indicating the presence or recurrent migration of Atl. *M. galloprovincialis* in those regions (Dieppe, Ostende, Ault, Dunkerque ‘Dun’, Figure S5).

Finally, one population from Korea (KOR, Figure S15) is completely composed of pure Med. *M. galloprovincialis*, corresponding to the known introduction in Asia (Han et al., 2016; McDonald et al., 1991). Another study showed that the introduction on the Pacific coast of the USA was similarly composed by pure Med. *M. galloprovincialis* (Simon et al., 2019). Those observations preclude the idea that previously observed Med. *M. galloprovincialis* introductions are related to dock mussels.

## 4 Discussion

We have uncovered a singular type of mussels in five ports in Western France. These dock mussel populations display a recent admixture pattern between non-native Med. *M. galloprovincialis* and South-Eu. *M. edulis*. While secondary admixture also occurred with genetic lineages encountered locally, dock mussels exhibit a high level of similarity between ports. In addition, our spatial sampling in ports allowed us to document the striking confinement and association of these genotypes to the interior of the ports, resulting in narrow shifts at port entrances. Some variation to this observation was, however, observed between ports, potentially due to their different layouts and conditions. Based on these results, we assume that dock mussels have been introduced.

By including and reanalysing *M. galloprovincialis* populations in Norway, experimental crosses, and newly identified admixed populations from several sites, we were able to compare admixture patterns between equivalent situations as well as between different genetic backgrounds and thus investigate the extent of parallelism in such secondary admixture processes.

### 4.1 The introduction of dock mussels and the timing of admixture

Dock mussels constitute homogeneous populations composed of around 70% Med. *M. galloprovincialis* ancestry, which may sometimes be called a ‘hybrid swarm’ due to a uni-modal distribution of hybrid indices and a complete mixing of ancestries along the genome (Allendorf, Leary, Spruell, & Wenburg, 2001; Beninde, Feldmeier, Veith, & Hochkirch, 2018; Jiggins & Mallet, 2000). We additionally show that there is ongoing secondary admixture between the dock mussel cluster and native genetic backgrounds, exemplified by the detection of F1 hybrids in Le Havre (Figure S26). While no F1 hybrids have been identified in the Bay of Brest by Newhybrids (Figure S27) – which most probably results from reduced power of identification between the two *M. galloprovincialis* lineages – the distribution of ancestries observed leaves little doubt that hybridisation is ongoing between dock mussels and Atl. *M. galloprovincialis* (Figures 3 and S23). Given the possibilities of local admixture, the relative global homogeneity of dock mussels could be explained either by the recentness of the introduction, by the existence of extrinsic or intrinsic barriers to introgressions, or by both.

The evidence of limited natural dispersal outside ports, presented in this study, provides a strong case for a saltatory colonisation of ports through human-mediated ‘jump dispersal’. In our view, the most parsimonious hypothesis of colonisation involves an initial admixture between pure Med. *M. galloprovincialis* and South-Eu. *M. edulis* in a yet unknown location, followed by secondary events of anthropogenically mediated dispersal. Both the genetic homogeneity of dock mussels and the absence of pure parental Med. *M. galloprovincialis* in all sampled ports provide arguments for this hypothesis. For instance in the Bay of Brest or in Saint-Malo, the presence of dock mussels with similar genetic compositions to the other ports (Figure S23), where the local native species is however different (i.e., predominantly Atl. *M. galloprovincialis* rather than *M. edulis*), suggests that the admixture with *M. edulis* happened before the introduction of dock mussels in these ports.

Ship traffic is thus likely to be the main source of these introductions to ports. The five studied infrastructures are large commercial and military ports that may have facilitated the primary introduction of mussels (C. L. Hewitt, Gollasch, & Minchin, 2009; Sylvester et al., 2011). Given the presence of marinas in the vicinity of the large studied ports and their colonisation by dock mussels, they constitute a possible way of secondary expansion at a regional scale. Indeed, marinas and associated activities, e.g. leisure boating, have been shown to contribute to regional NIS expansion (Clarke Murray, Pakhomov, & Therriault, 2011) and create chaotic genetic structure in both native and non-native species inhabiting these artificial habitats (Guzinski, Ballenghien, Daguin-Thiébaut, Lévêque, & Viard, 2018; Hudson, Viard, Roby, & Rius, 2016). For now, in the Bay of Brest, only the marinas close to the large port contained dock mussels. The other marinas outside of the bay (e.g. Camaret and Morgat, Figure S11 Brest-11 and 13 respectively) – potentially exchanging a lot of traffic with Brest marinas – did not, and this supports the absence of a secondary introduction. Colonisation seems therefore so far limited to large port infrastructure, and nearby marinas, with dispersal due to large vessel traffic. This situation might nonetheless change over time, and genetic monitoring should be pursued.

We have estimated an admixture time for dock mussels of 4 to 28 years ago. In addition to the inherent difficulty of this dating and the limitation of our dataset, we note that this estimate assumes neutrality, and no gene flow since admixture. We have evidence, at least in Le Havre, of a constant input of new chromosome tracts from the native *M. edulis*. In addition, we can suspect a continuing propagule pressure of Med. *M. galloprovincialis* from the maritime traffic. It is also likely that selection acts to maintain parental gene combinations against recombination (Bierne et al., 2006; Simon et al., 2018). Both effects, gene flow and selection, tend to bias the date estimates towards more recent times (Corbett-Detig & Nielsen, 2017). A precise estimation of the admixture event will require a recombination map in mussels and the distribution of ancestry track lengths along the genome of admixed individuals.

Interestingly, in 1978, Prof. David Skibinski analysed hybrids from natural populations in the Swansea region (UK) with allozymes (Skibinski, Beardmore, & Ahmad, 1978) and noticed that the ‘King’s dock’ populations (Swansea port) were unusual (Figure S39). Those populations showed linkage and Hardy-Weinberg equilibria, and intermediate allele frequencies between *M. edulis* and *M. galloprovincialis*. A closer look at the allele frequency shows that, at one particular allozyme subsequently shown to differentiate Atl. from Med. *M. galloprovincialis* (*Ap*, Quesada, Zapata, & Alvarez, 1995), King’s dock populations had allele frequencies that were closer to those of Med. mussels than to local Atl. *M. galloprovincialis*. This evidence suggests that introduced dock mussels were already present, and already admixed with *M. edulis* at the same level in the Swansea port, 40 years ago. This provides further indication that our estimate of admixture time is potentially underestimated. The term ‘dock mussels’ was chosen in reference to this work. We do not know if dock mussels persisted in the Swansea port and this matter needs further investigation.

Both of the above considerations suggest that the admixture event leading to dock mussels is a few decades old. The mussel introductions therefore appear relatively recent, especially compared to the several centuries over which human maritime traffic could have been a vector of fouling NIS (J. T. Carlton & Hodder, 1995). However, as stated by Hulme (2009), ‘the highest rates of introductions in Europe occurred in the last 25 years’ (p. 11) due to an increase in the rate of global exchange. It is therefore possible that dock mussels were spread to multiple ports in this time-frame, especially if a large propagule size is a prerequisite for successful introduction under strong demographic and/or genetic Allee effect (Barton & Turelli, 2011).

Dock mussels are not isolated cases of anthropogenic hybridisation in the *M. edulis* species complex. Recently, Zbawicka et al. (2018) reported the presence of an admixed population between introduced Med. *M. galloprovincialis* and native *M. platensis* close to the city of Puerto Madryn in the middle of the Atlantic coast of Argentina. Their randomly ascertained SNPs did not allow a precise analysis of individual admixture proportions but the average admixture appeared well-balanced. In this issue, Popovic et al. (2019) reported two independent introductions of *M. galloprovincialis* in Australia, one by the Atl. *M. galloprovincialis* in Batemans Bay and the other by the Med. *M. galloprovincialis* in Sydney Harbour, both accompanied by admixture with the native genetic background (*M. planulatus*). In New-Zealand, Gardner, Zbawicka, Westfall, and Wenne (2016) found evidence suggesting possible admixture between introduced *M. galloprovincialis* and the native *Mytilus* species. Such observations are additional indications of the frequent occurrence of the admixture process where *M. galloprovincialis* has been introduced in an area already inhabited by a native lineage of *Mytilus*.

Conversely, there was little to no introgression during the introduction of Med. *M. galloprovincialis* in California (Saarman & Pogson, 2015) and Asia (Brannock, Wethey, & Hilbish, 2009, and Korean sample in this study) where the native species is *M. trossulus*. Those last two cases may be the result of increased intrinsic and extrinsic reproductive isolation with *M. trossulus* that is much more divergent. Alternatively, the introduction and initial spread may have happened in a place devoid of native *M. trossulus* and with a more balanced demographic context than for dock mussels. Finally, events of admixture are not restricted to *M. galloprovincialis*. For instance, evidence of admixture has been found in the Kerguelen Islands (Fraïsse, Haguenauer, et al., 2018; Zbawicka, Gardner, & Wenne, 2019).

### 4.2 Confinement of the introduced mussels, local introgression and potential impacts

In all studied ports, the introduced dock mussels form sharp human-induced hybrid zones at the port entrance. By contrast, natural clines in mussels are usually on the order of tens to hundreds of kilometres (Lassen & Turano, 1978; Strelkov, Katolikova, & Väinolä, 2017; Väinolä & Hvilsom, 1991). Saarman and Pogson (2015) also found differences in the sharpness of genomic clines between the anthropogenically driven contact in California and old natural secondary contacts. If the natural clines are due to post-zygotic selection in a tension zone model (Barton & Hewitt, 1985; Bierne, David, Boudry, & Bonhomme, 2002), then the narrow clines in ports imply additional processes. Those processes could include habitat choice during the larval settlement stage at a small spatial scale (Bierne et al., 2003; Comesaña & Sanjuan, 1997; Katolikova, Khaitov, Väinölä, Gantsevich, & Strelkov, 2016) or early stage larval or post-settlement ecological selection to the port environment. For instance, selection in mussels could act through attachment strength (Willis & Skibinski, 1992), pollution tolerance (Loria, Cristescu, and Gonzalez, 2019, for a review; and McKenzie, Brooks, and Johnston, 2011, for an example in a bryozoan), or competition for space linked to different growth rates (Branch & Steffani, 2004; Saarman & Pogson, 2015). Additionally, genetic differentiation in mussels has been shown to be associated with sewage treatment plants (Larsson, 2017; Larsson et al., 2016).

Although our sampling around ports was not exhaustive, dock mussels do appear to be restricted to the port interiors, with only a few introduced mussels detected in distant populations. While the presence of introduced migrants up to 30 km from ports may appear concerning, most distant individuals are hybrids between dock mussels and the local background (Figures 3 and S26-S27). Therefore, we can hypothesise that the propagule pressure from ports will be swamped by large native populations for most of the ports. Conversely, native mussels are relatively rare inside the ports (except for Brest). Were they more numerous, hybridisation might favour an increase in introgression by the possibility of backcrossing to the native mussels. The concern of genetic pollution seems increased in the Bay of Brest where the potential for dispersion and hybridisation appears greater. Additionally, populations of introduced mussels were found in basins closed by locks (Saint-Malo, Le Havre, Cherbourg, Saint-Nazaire). In such contexts, both the exit and entry of mussel larva from any species may be limited and those populations may act as reservoirs of introduced backgrounds.

The introduction cases in ports and Norway agree well with the expectation of asymmetric introgression from the established taxon into the propagating one (Barton, 1979a; Currat, Ruedi, Petit, & Excoffier, 2008; Moran, 1981). Introgression levels can reach much higher levels in a moving hybrid zone than in stable ones (Currat et al., 2008). Genetic pollution by NIS is unlikely to be substantial during invasion, while the reverse is true although less concerning (Currat et al., 2008). However, when the invasion wave is halted and trapped at a natural barrier, density trough, or ecotone, introgression can start to proceed in both directions. Introgression of native mussel populations by dock mussel alleles could therefore become a concern. Nonetheless, the evolutionary future of Med. *M. galloprovincialis* alleles in native populations are hard to predict. They could for example be counter-selected like in the westslope cutthroat trout (*Oncorhynchus clarkii lewisi*), where introgression impacts the fitness of native populations and selection against introduced alleles in wild populations seems to be acting (Kovach et al., 2016; Muhlfeld et al., 2009). While this is an interesting outcome, some parts of the native genome may still be impacted. Indeed, in the brown trout (*Salmo trutta*), a haplotype-based method showed that residual introduced tracts are present in native populations and go undetected by classical ancestry estimation methods (Leitwein, Gagnaire, Desmarais, Berrebi, & Guinand, 2018).

The Bay of Brest is an interesting case study both in terms of implicated species – this is a crossroad between three lineages – and of introduction. In this area, unlike the other ports, introduced mussels were found beyond the major human-made structures. Yet, even in distant sites from ports, mussels were predominantly found on artificial structures (buoys, pillars, piers, etc.). However, this observation may be more related to space competition with oysters on natural sites than to habitat selection, as finding mussels of any type on natural rocky shores in the bay was difficult.

The spread of dock mussels in the Bay of Brest might be due to several interacting factors. First, the port – and notably the commercial area – has a more open layout compared to the other four ports (some of which, such as Saint-Nazaire, have locks at their entry). Second, compared to other ports, habitats suitable for dock mussels might have been available. Third, the closer genetic distance with Atl. *M. galloprovincialis* when compared to *M. edulis* might facilitate hybridisation by avoiding stronger reproductive incompatibilities (both pre- and post-zygotic). Therefore, the prediction of the invasion by dock mussels will require a thorough understanding of the reproductive incompatibilities between non-indigenous and native mussels (Blum, Walters, Burkhead, Freeman, & Porter, 2010; Hall, Hastings, & Ayres, 2006).

When interacting species have accumulated too many incompatibilities for hybridisation to lead to viable and fertile offspring, inter-specific mating represents lost reproductive effort (Allendorf et al., 2001). For less reproductively isolated species, hybridisation has been considered by Mesgaran et al. (2016) as a way to escape demographic Allee effects during colonisation. As small introduced mussel populations may suffer from a strong Allee effect, hybridisation has potentially provided the initial demographic boost to the first introduction of Med. *M. galloprovincialis*. Conversely, hybrid breakdown would have impeded both the introduction of the hybrid background, which would then have required a tremendous propagule pressure from maritime traffic. The same applies to the subsequent spread of dock mussels, even if fitter (Barton & Turelli, 2011), and this could explain their confinement inside ports. Stochasticity (drift and variation in population density and dispersal) could free the introduced background after a lag time (Piálek & Barton, 1997). Although the delay is expected to be long, confined dock mussel populations could represent hidden bombshells able to escape and spread globally in the future.

The introduced dock mussels display an important component of *M. galloprovincialis* ancestry. Based on the worldwide spread and displacement of local species (Branch et al., 2008; Gardner et al., 2016; Saarman & Pogson, 2015), *M. galloprovincialis* is expected to have a competitive advantage in diverse conditions. It is thus tempting to predict that dock mussels should spread. However, the specific ecological characteristics of these dock mussels as well as the native mussels that first colonized the study ports are unknown, which strongly limits any attempts to predict the impact and the fate of the introduced populations. Their local impact will require further investigation. Nonetheless, we are left with the fact that in ports and in natural environments in the Bay of Brest, dock mussels have probably displaced the native lineages. Michalek, Ventura, and Sanders (2016) report impacts of hybridisation on *Mytilus* aquaculture in Scotland and Larraín et al. (2018) raise concerns of economic impacts in Chile. In Scotland, a recent demographic increase of *M. trossulus* has produced large losses to *M. edulis* aquaculture due to their colonisation of culture ropes and their shell being more fragile (Beaumont, Hawkins, Doig, Davies, & Snow, 2008; Dias et al., 2009). In Brittany and Normandy, most of the cultured mussels are imported spat from the bay of Biscay because *M. edulis* is easier to cultivate, with a shorter reproduction period, and has a higher commercial value for consumers than Atl. *M. galloprovincialis*. Therefore, spat collection of introduced mussels and involuntary culture of the wrong genetic background should impact the quality of cultured mussels and the growing cycle used in mussel farms, but only in case of a massive invasion.

While there may be few direct impacts of dock mussels on native and cultivated mussel populations, indirect effects via parasite hitch-hiking during introduction and their transmission to native species has been documented both in terrestrial and marine systems (Prenter, MacNeil, Dick, & Dunn, 2004; Torchin, Lafferty, & Kuris, 2002). We should therefore be concerned about the potential parasites NIS may have brought into natural and cultivated populations (‘spillover’ effect). Additionally, the ‘spillback’ effect, due to the NIS being a competent host for native parasites and constituting a new reservoir for local diseases, should not be neglected (Kelly, Paterson, Townsend, Poulin, & Tompkins, 2009). We can note that, at this time, we did not detect the *M. trossulus* transmissible cancer in dock mussels (Metzger et al., 2016; Riquet et al., 2017). On an evolutionary perspective, the introduction of Atl. and Med. *M. galloprovincialis* into *M. edulis* ranges and the following gene flow may confer some parasitism adaptations to the native species. For example, it has been demonstrated that *M. galloprovincialis* is more resistant to Pea crab parasitism than *M. edulis* living in the same region (Seed, 1969).

If management is to be considered, multiple steps need to be taken. First, genetic detection methods such as the one used in this work need to be routinely used to assess the extent of the introduction in all large North-European ports. Second, the introduction is to be followed in time and space around the points of introduction, notably to determine the speed of the expansion front, if any, and thus ascert if dock mussels are becoming invasive. Third, to understand the introduction process in the different ports, there needs to be an integration of genetics and ecology (Lawson Handley et al., 2011). However, we have a large gap in our ecological knowledge of the port environments and what influences mussel populations. A thorough study of the ecology of mussels in ports will be needed to untangle the roles of ecological variation in the distribution of dock mussels. Both habitat choice and postsettlement selection are likely to play a role. The final objective would be to produce a fine scale environmental niche model. Fourth, a vector risk assessment will be necessary to predict the possible human induced secondary displacements (e.g., Herborg, O’Hara, & Therriault, 2009). Finally, at a local scale, larval dispersal through oceanographic constraints will play a major role in the potential spread of dock mussels and dispersal models for NIS in ports will be needed (see David, Matthee, Loveday, & Simon, 2016, for an example at a large scale). While some studies of water flows, tide or wave physical constraints in ports of the English Channel exist (Guillou & Chapalain, 2011, 2012; Jouanneau, Sentchev, & Dumas, 2013), none include a biological module. A study of wave entrance in the southern basin of Le Havre would suggest the likely dispersal of *M. edulis* larvae within this basin (Guillou & Chapalain, 2012), while the whole basin proved populated by dock mussels, providing further evidence for habitat choice or early stage selection. Overall, a large effort will be needed to produce consistent models of larval dispersal at the scale of ports of interest. At a medium scale, in the Bay of Brest, the model of Bessin (2017) could help investigate the relative weights of dispersion, habitat selection and ecological constraints on the distribution of genetic backgrounds. At any rate, managing dock mussels will require the combination of vector risk assessment, network theory, and environmental niche and oceanographic models to build a complete risk assessment model (Frost et al., 2019; Herborg et al., 2009; Hulme, 2009).

In addition to allowing the study of introduction and evolutionary biology, the *Mytilus* model could be of interest for the recent field of urban ecology and evolution, investigating the impact of urbanisation on evolutionary trajectories and the feedbacks with the environment (Rivkin et al., 2018; Thompson, Rieseberg, & Schluter, 2018). The marine environment is not left untouched by urbanisation and human infrastructures have large impacts on coastal communities and their abiotic conditions (Critchley & Bishop, 2019; Mayer-Pinto et al., 2018). This is the ‘Ocean Sprawl’, in the words of Duarte et al. (2012), which has broad effects encompassing connectivity modifications and environmental and toxicological changes (for a review see Firth et al., 2016).

### 4.3 Parallelism of distortions

The parallelism in allele frequency distortions that we observed between admixture events, suggests that patterns produced during such events can be highly repeatable. This is probably due to a combination of processes. As discussed above, port introductions are expected to partly share a pre-introduction history of admixture. The two introduced Atl. *M. galloprovincialis* populations we studied in Norway are also likely to share the same history of admixture. However, the composition in *M. edulis* ancestry of these populations is in accordance with an independent admixture event with the local *M. edulis* background. Naturally admixed Atl. *M. galloprovincialis* combine an old history of introgression during glacial oscillation periods (Fraïsse, Roux, et al., 2018; Roux et al., 2014) with ongoing local introgression from the native *M. edulis* populations in direct contact within the mosaic hybrid zone observed today (Fraïsse et al., 2016; Simon et al., 2019).

Shared colonisation history cannot be the whole story, however, because we also found repeatable patterns between admixture events that must be considered independent. This includes not only the comparisons of natural admixture to the introduced Atl. *M. galloprovincialis* (involving two different backgrounds of *M. edulis*: South- and North-Eu.), but also the comparison of port samples to experimental backcrosses.

Our comparison of Atl. *M. galloprovincialis* admixtures includes populations with a wide variety of contributions from the parental lineages. These range from a high *M. edulis* contribution in JER to a high Atl. *M. galloprovincialis* contribution in MCH. The high positive correlations of distortions observed between all Atl. *M. galloprovincialis* admixture, despite variable contributions of the two parental backgrounds, is particularly interesting. The calculation of *D* corrects for ancient introgression of parental backgrounds, and we are unlikely to have missed a hidden parental population given our broad geographic survey (this work and Simon et al., 2019) and the large-scale genetic panmixia usually observed in mussels outside hybrid zones (e.g. East vs. West Mediterranean Sea). Genomic regions do tend to deviate consistently toward an excess of *M. galloprovincialis* ancestry or an excess of *M. edulis* ancestry. This suggests selective processes and a shared architecture of the barrier to gene flow. A first possible explanation is that some loci are closer to barrier loci than others (Ravinet et al., 2017). Barrier loci can be local adaptation genes or genetic incompatibilities. Schumer et al. (2018) found that in several events of admixture between swordtail fish species contributing differently to the resulting population, local ancestry were nonetheless positively correlated. They showed that parallel correlations, despite opposite parental contributions, can be the result of selection in the same direction to resolve pairwise epistatic incompatibilities. In addition, an interesting interpretation of the parallelism observed in mussels would be that our loci belong to genomic regions with different rates of recombination. *M. edulis* and *M. galloprovincialis* are close to the 2% net synonymous divergence limit (1.89%), where there is a high probability of strong reproductive isolation, either due to physical constraints or sufficient accumulation of incompatibilities (Roux et al., 2016). They are therefore expected to be incompatible at a high number of differentiated sites (Simon et al., 2018). With such a highly polygenic determinism of post-zygotic selection one expect a correlation between recombination rates and introgression (Barton & Bengtsson, 1986), which has recently been observed in multiple study systems (*Mimulus*, Aeschbacher, Selby, Willis, and Coop, 2017; sea bass, Duranton et al., 2018; oyster, Gagnaire et al., 2018; stickleback, Roesti, Moser, and Berner, 2013; swordtail fish, Schumer et al., 2018 or *Heliconius*, Martin, Davey, Salazar, and Jiggins, 2019).

While patterns of hybridisation are strongly repeatable when the same *M. galloprovincialis* lineages are involved, equally notable is the lack of repeatability with different lineages. A possible explanation is that different sets of incompatible loci may be implicated in the reproductive isolation between *M. edulis* and the two *M. galloprovincialis* lineages. However, the history of divergence between the two *M. galloprovincialis* lineages is much younger than the divergence with *M. edulis* and most of the fixed mutations are expected to be shared by the two lineages (Fraïsse et al., 2016). Additionally, Atl. *M. galloprovincialis* is in contact with *M. edulis* while Med. *M. galloprovincialis* is not. Atl. *M. galloprovincialis* has experienced a punctuated history of introgression possibly swamped by bi-stable incompatibilities with an asymmetric advantage to the *M. edulis* allele (Fraïsse et al., 2016; Gosset & Bierne, 2013; Simon et al., 2019). This differential introgression might have erased, or even reversed, the selective effects in the two *M. galloprovincialis* backgrounds. This hypothesis requires further theoretical and experimental investigation. Finally, given that karyotypic differences have been suggested between the two *M. galloprovincialis* lineages (Martínez-Lage, González-Tizón, & Méndez, 1996), they potentially exhibit different recombination landscapes impacting the outcome of distortions.

## 5 Conclusion

*Mytilus* mussels, with their introduction and hybridisation potential, are a particularly useful model for studying the parallelism of admixture events, and the range of outcomes of introductions with hybridisation. Our study shows that admixture between the same genetic backgrounds are highly repeatable. This repeatability can be explained both by a shared history of pre-introduction admixture and parallel genomic processes. One category of anthropogenic hybridisations, the ‘dock mussels’, exhibit homogeneous patterns of admixture among all studied populations, and appear to be restricted to environments of large commercial ports. Follow-up investigations will be needed to understand how selection, hybridisation, environmental conditions and dispersal are shaping the distribution and genomic architecture of these dock mussels and similar introductions.

## Data availability

Raw data and scripts are available as a Zenodo archive 10.5281/zenodo.3375381.

## Acknowledgements

Data used in this work were partly produced through the genotyping and sequencing facilities of ISEM and LabEx CeMEB, an ANR “Investissements d’avenir” program (ANR-10-LABX-04-01). This project benefited from the Montpellier Bioinformatics Biodiversity platform supported by the LabEx CeMEB. We are grateful to the Marine Observation Department (SMO) at the Roscoff Biological Station for sampling in the bay of Brest. We thank Laure Paradis for providing maps of the study area and her assistance with map editing. We thank Prof. D. Skibinski for communications about the Swansea King’s Dock mussels. We also thank people who contributed to the acquisition of samples or gave advice: Galice Hoarau, Alison Gallet, Filip Volckaert, Association Port Vivant (Le Havre), CRC Normandie, Julien Normand, Sophie Arnaud-Haond, Lea-Anne Henry, Thierry Comtet, Pieternella Luttikhuizen and Jørgen Berge (UiT The Arctic University of Norway). We acknowledge the help of staff and contractors at CNR International UK Ltd for helping to collect and preserve the samples from the Murchison oil station. We thank Richard Crooijmans and Bert Dibbits from Animal Breeding and Genomics Wageningen University & Research for supplying material. This work was supported by the ANR Project HySea (ANR-12-BSV7-0011). Maria Skazina was supported by the Russian Science Foundation project 19-74-20024.

