## Supplementary material for "Replicated anthropogenic hybridisations reveal parallel patterns of admixture in marine mussels"

[1]ISEM, Univ Montpellier, CNRS, EPHE, IRD, Montpellier, France.
[2]Section for Marine Living Resources, National Institute of Aquatic Resources, Technical University of Denmark, Silkeborg, Denmark.
[3]SARL TOXEM, Le Havre, France.
[4]Ifremer Unité Biogéochimie et Écotoxicologie, Centre Atlantique, Nantes.
[5]SAS Eurêka Mer, Lézardrieux, France.
[6]Wageningen Marine Research, P.O. Box 57, 1780 AB Den Helder, The Netherlands.
[7]Wageningen University, Aquatic Ecology and Water Quality Management Group, Droevendaalsesteeg 3a, 6708 PD Wageningen, The Netherlands.
[8]Ifremer, SG2M-LGPMM, Laboratoire de Génétique et Pathologie des Mollusques Marins, La Tremblade, France.
[9]St. Petersburg State University, Universitetskaya Emb. 7/9, St. Petersburg 199034, Russia.
[10]Laboratory of Monitoring and Conservation of Natural Arctic Ecosystems, Murmansk Arctic State University, Kapitana Egorova Str. 16, Murmansk 183038, Russia.
[11]Department of Biology & CESAM, University of Aveiro, Campus Universitário de Santiago, 3810-193 Aveiro, Portugal.
[12]CBET Research Group, Dept. of Zoology and Animal Cell Biology, Fac. Science and Technology and Research Centre for Experimental Marine Biology and Biotechnology (PiE-UPV/EHU), University of the Basque Country (UPV/EHU), Bilbao, Spain.
[13]Department of Genetics, University of Cambridge, Downing St. Cambridge, CB23EH, UK.
[14]Sorbonne Universités, UPMC Univ Paris 06, CNRS, UMR 7144, Department AD2M, Station Biologique, Roscoff, France.

### Supplementary information

#### Contents

**1 Admixture time** **4**

**2 Supplementary Figures** **4**

**3 Supplementary Tables** **30**

| | | |
|---|---|---|
| S5 | Genetic map | 40 |
| S6 | Fst values between reference groups. | 41 |
| S7 | Fst values and significance between ports. | 41 |
| S8 | Fst values and significance between Cherbourg years. | 41 |
| S9 | Fst values and significance between naturally and Norway admixed. | 41 |
| S10 | Statistical tests of ancestry comparisons in naturally and Norway admixed | 42 |
| S11 | Statistical tests of ancestry comparisons in dock mussels | 43 |
| S12 | Statistical tests of ancestry comparisons in Cherbourg | 43 |
| S13 | Statistical tests of ancestry comparisons in Brittany | 43 |
| S14 | Admixture time estimation | 44 |

### 1 Admixture time

To estimate simply admixture time, `Structure` runs were used as indicated in Falush, Stephens, and Pritchard (2003) using the linkage model. For each port (or year for Cherbourg) only admixed dock mussels individuals were considered with additional reference populations being edu_eu_south and gallo_med. The popflag column was set to 1 for reference individuals to better estimate the parental allele frequencies. Only markers on the genetic map were used to avoid bias through undetected linkage with markers that could not be included in the genetic map analysis. Options used: LINKAGE = 1, PFROMPOPFLAGONLY = 1, NOADMIX = 0, LOG10RSTART = -2, LOG10RMIN = -3, LOG10RMAX = 2, LOG10RPROPSD = 0.1. See the parameter files for more details. Results of average $r$ for 25 runs are presented in table S14, the number of generations is given by $r \times 100$ as genetic distance are given in cM.

### 2 Supplementary Figures

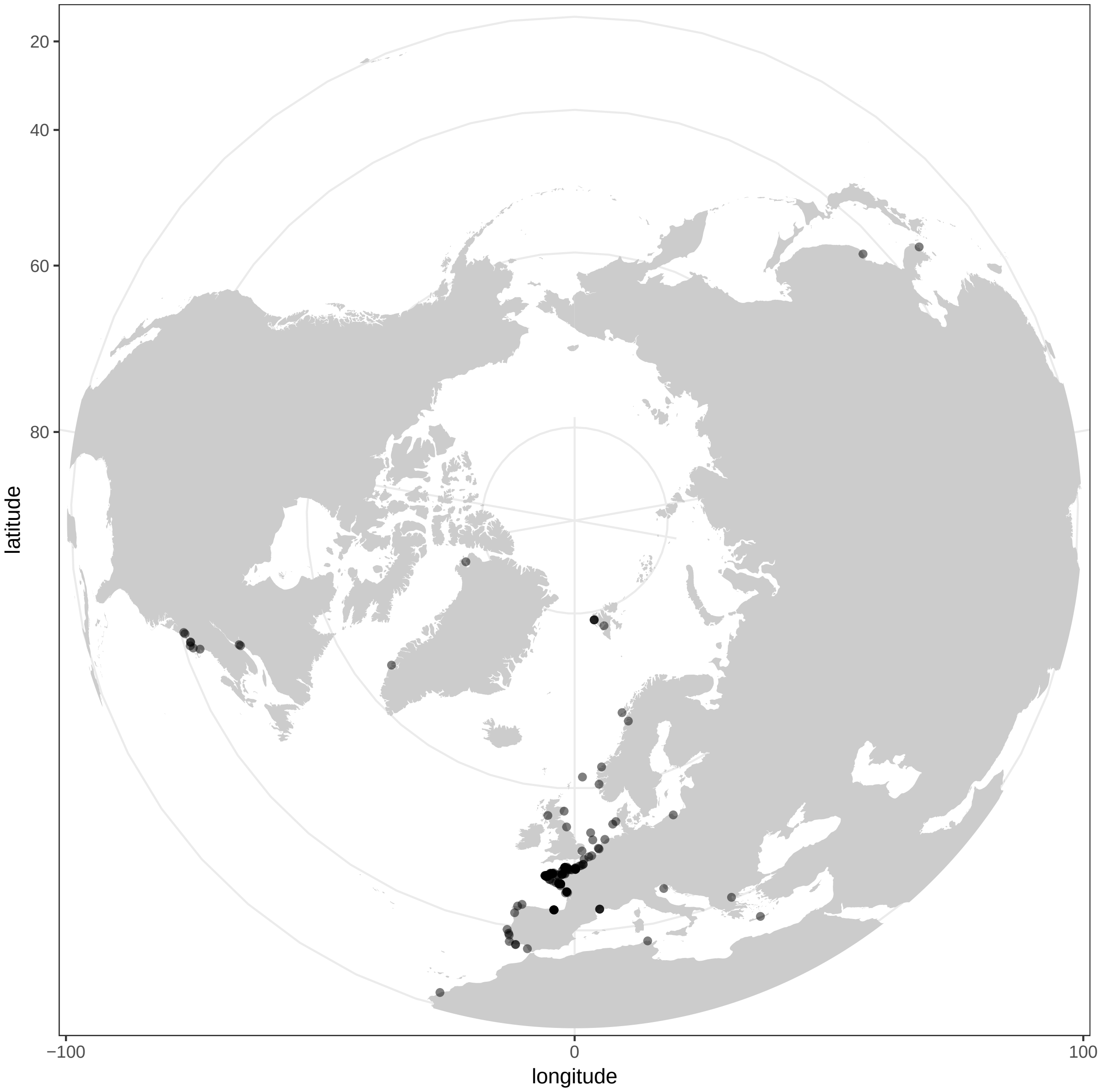

Figure S1: Location of all samples considered in the study.

**Steps data**

| | | | |
|---|---|---|---|
| Tip1 | | 96 DW tip comb | |
| Pick-Up | | Tip Plate | |
| Bind | | Sample | |
| | Beginning of step | Precollect | No |
| | | Release time, speed | 00:00:30, Bottom mix |
| | Mixing / heating: | Mixing time, speed | 00:04:30, Half mix |
| | | Heating during mixing | No |
| | End of step | Postmix | No |
| | | Collect count | 3 |
| | | Collect time [s] | 1 |
| Wash 1 | | MB3 | |
| | Beginning of step | Precollect | No |
| | | Release time, speed | 00:00:10, Bottom mix |
| | Mixing / heating: | Mixing time, speed | 00:01:00, Half mix |
| | | Heating during mixing | No |
| | End of step | Postmix | No |
| | | Collect count | 3 |
| | | Collect time [s] | 1 |
| Wash 2 | | MB4 | |
| | Beginning of step | Precollect | No |
| | | Release time, speed | 00:00:10, Bottom mix |
| | Mixing / heating: | Mixing time, speed | 00:01:00, Half mix |
| | | Heating during mixing | No |
| | End of step | Postmix | No |
| | | Collect count | 3 |
| | | Collect time [s] | 1 |
| Wash 3 | | MB5 | |
| | Beginning of step | Precollect | No |
| | | Release beads | No |
| | Mixing / heating: | Mixing time, speed | 00:00:30, Slow |
| | | Heating during mixing | No |
| | End of step | Postmix | No |
| | | Collect beads | No |
| Elution | | Elution | |
| | Beginning of step | Precollect | No |
| | | Release time, speed | 00:00:15, Bottom mix |
| | Mixing / heating: | Shake 1 time, speed | 00:01:00, Medium |
| | | Shake 2 time, speed | 00:00:10, Bottom mix |
| | | Loop count | 8 |
| | | Heating temperature [°C] | 72 |
| | | Preheat | Yes |
| | End of step | Postmix | No |
| | | Collect count | 3 |
| | | Collect time [s] | 30 |
| ReleaseBeads1 | | MB5 | |
| | | Release time, speed | 00:00:10, Half mix |
| Leave | | MB5 | |

Figure S2: Program for DNA extraction with the Kingfisher Flex robot.

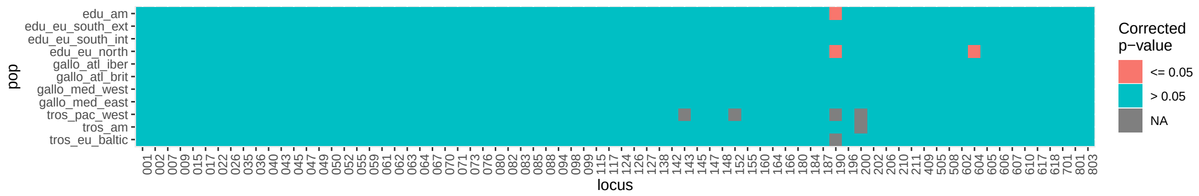

Figure S3: Hardy-Weinberg equilibrium test in each reference population. Benjamini-Yekutieli corrected $p$ values are used.

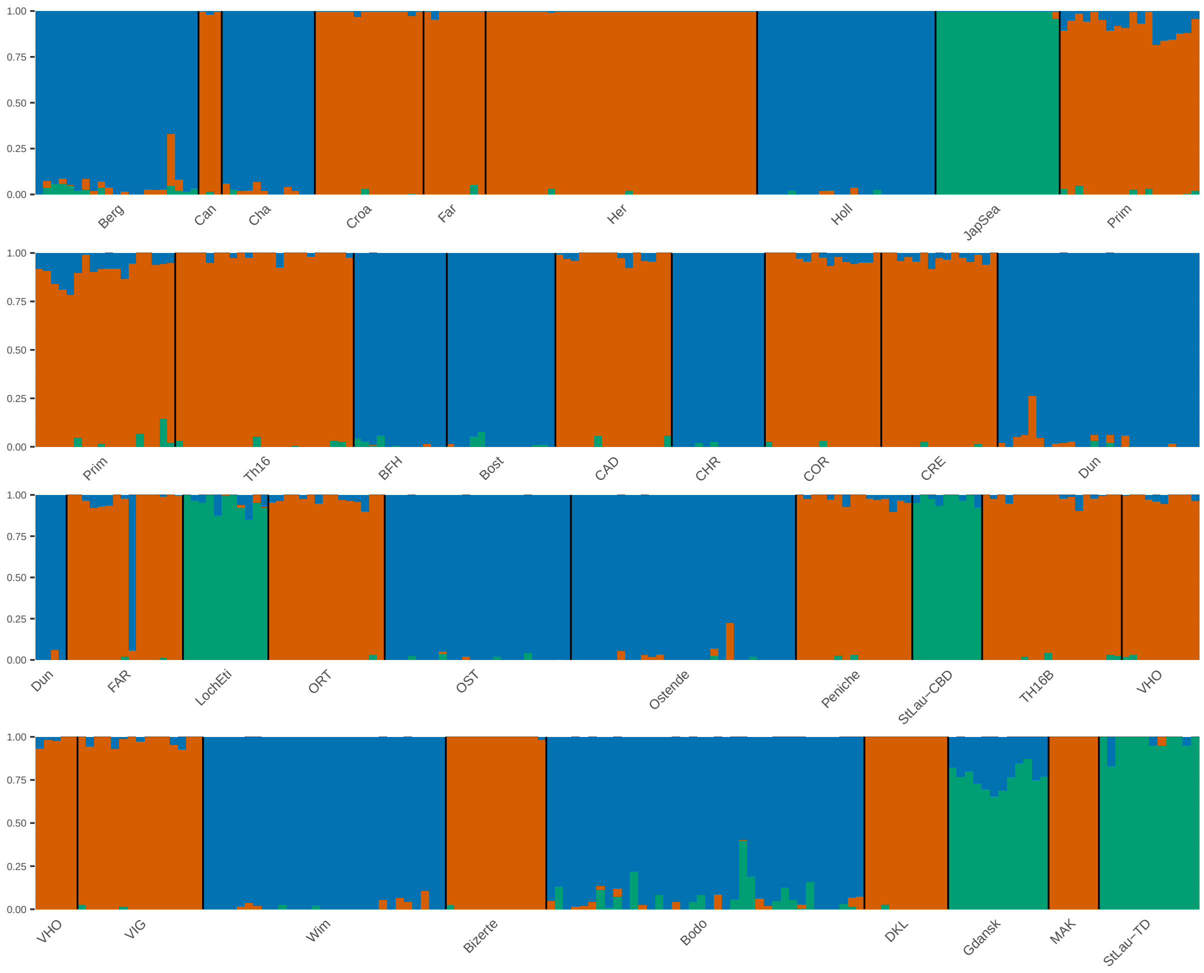

Figure S4: `Admixture` results for reference groups, $K = 3$, before filtration.

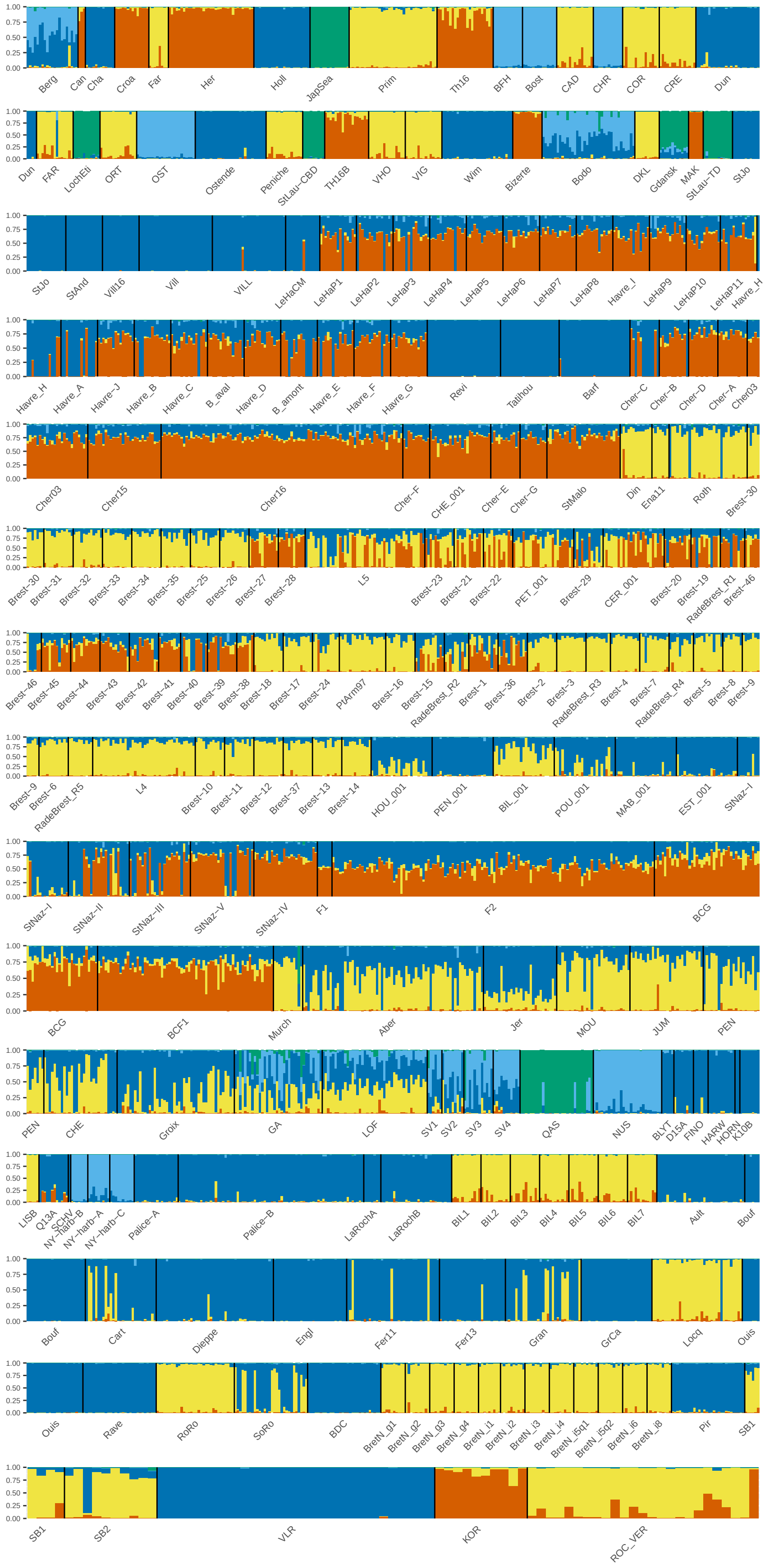

Figure S5: `Structure` results with all individuals, $K = 6$ major cluster.

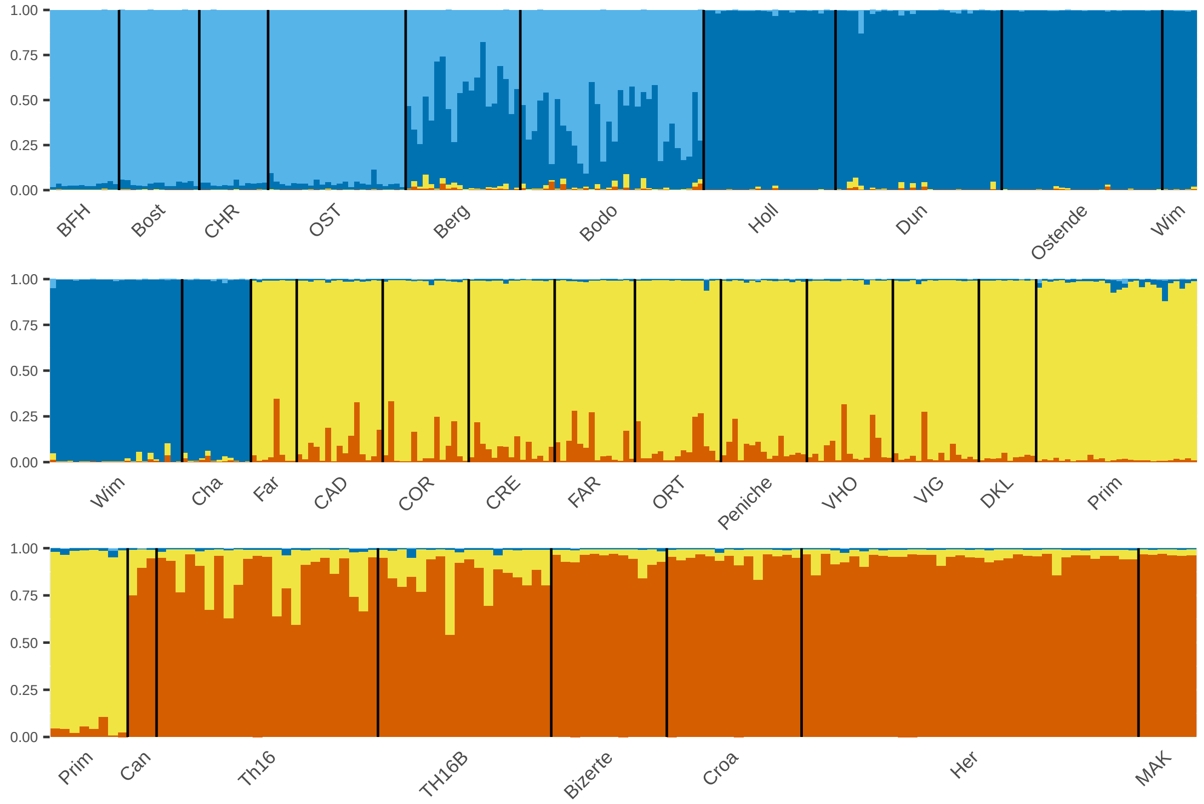

Figure S6: `Structure` results with all individuals excluding the ones with *M. trossulus* ancestry, $K = 4$ major cluster. Subset corresponding to reference populations only including individuals from the reference groups (table S4) after filtration.

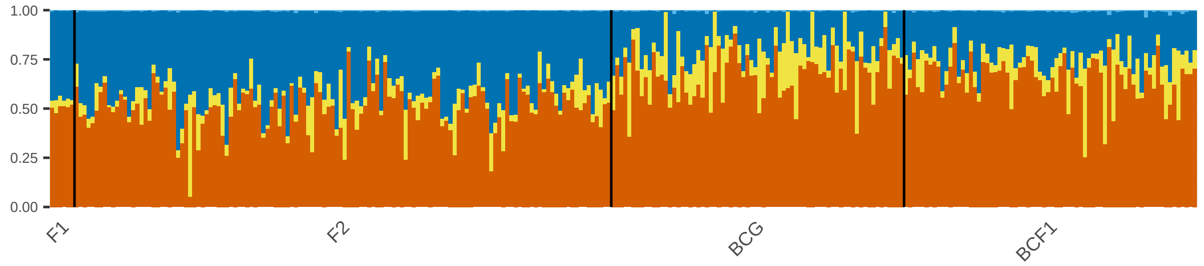

Figure S7: Subset of the lab crosses.

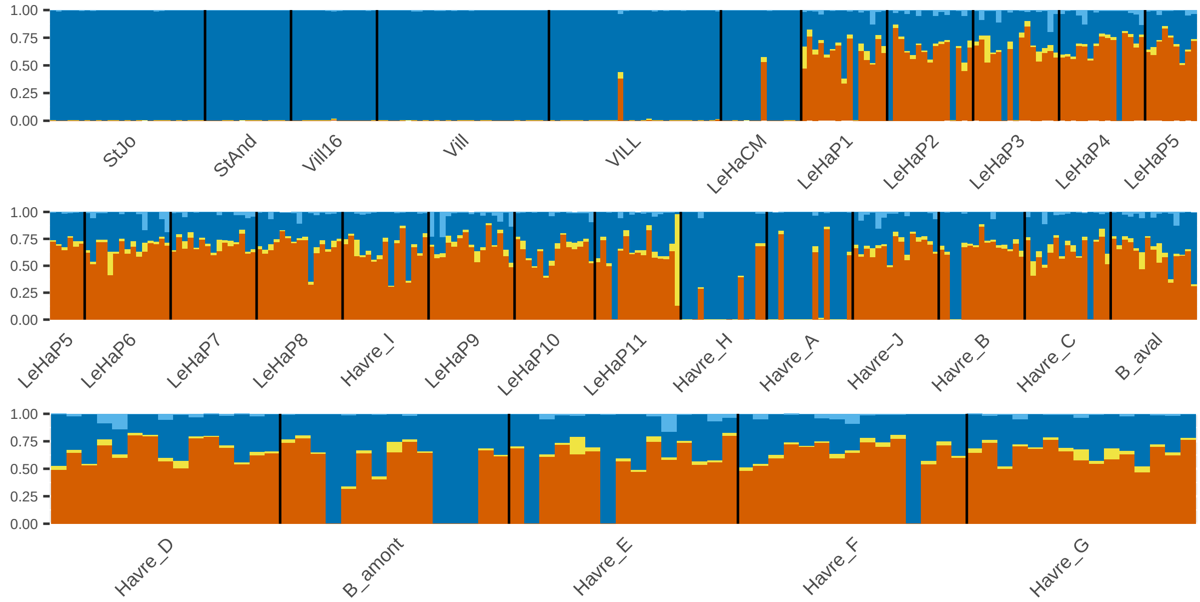

Figure S8: Subset of the port of Le Havre.

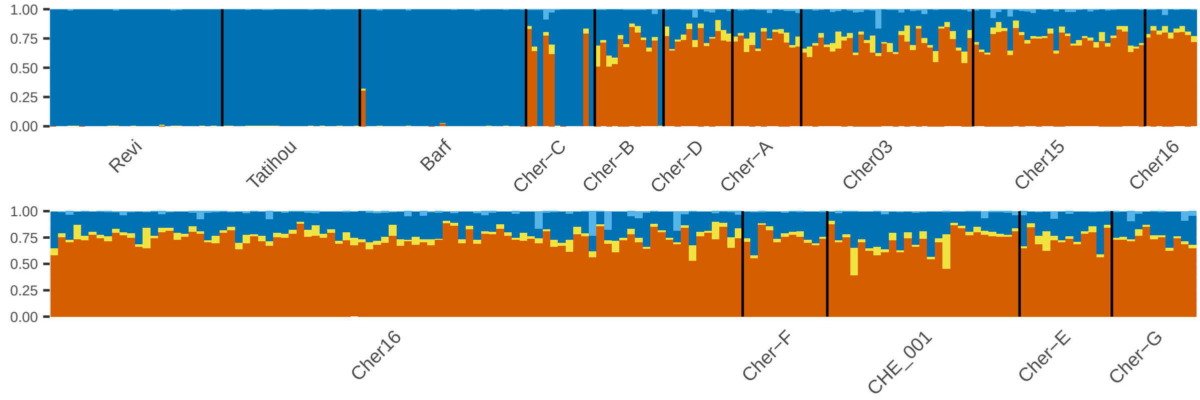

Figure S9: Subset of the port of Cherbourg.

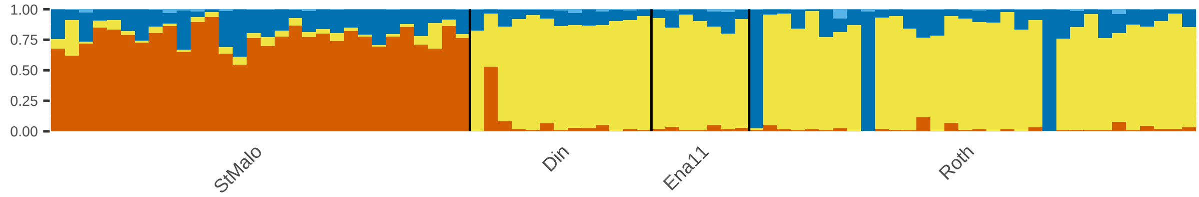

Figure S10: Subset of the port of Saint-Malo.

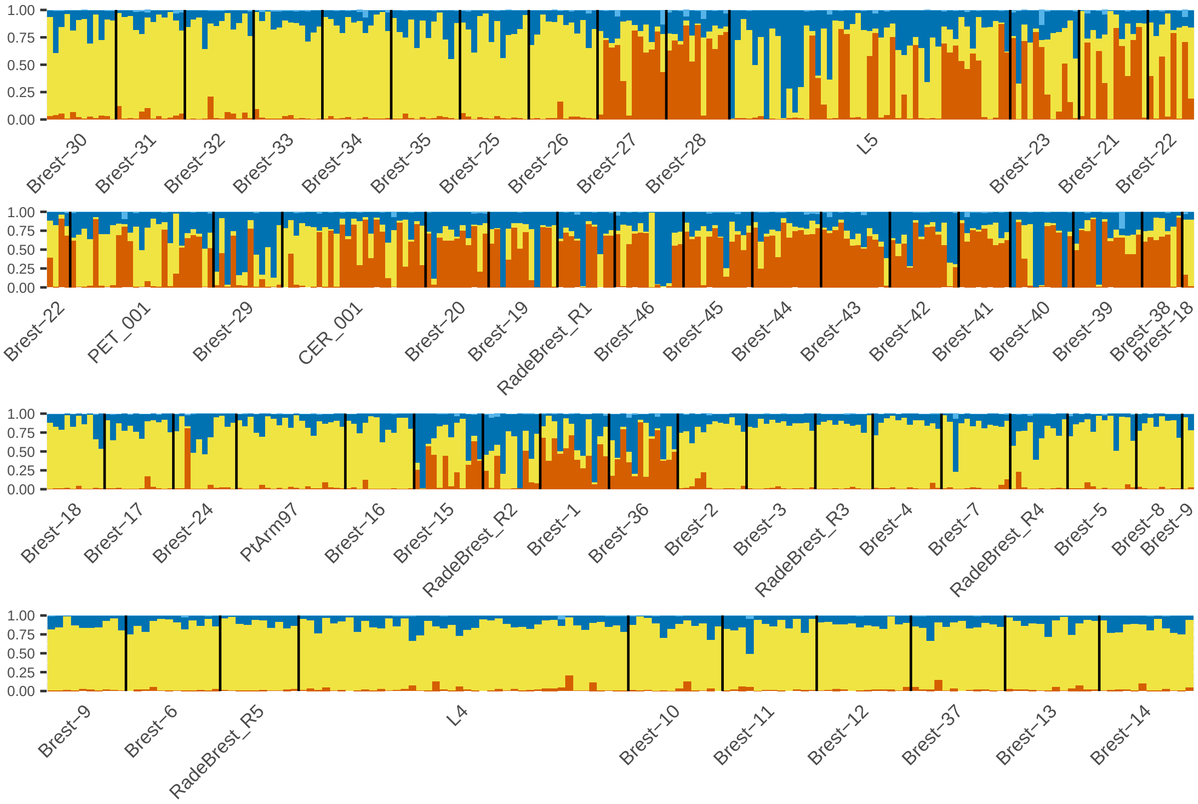

Figure S11: Subset of the bay of Brest area.

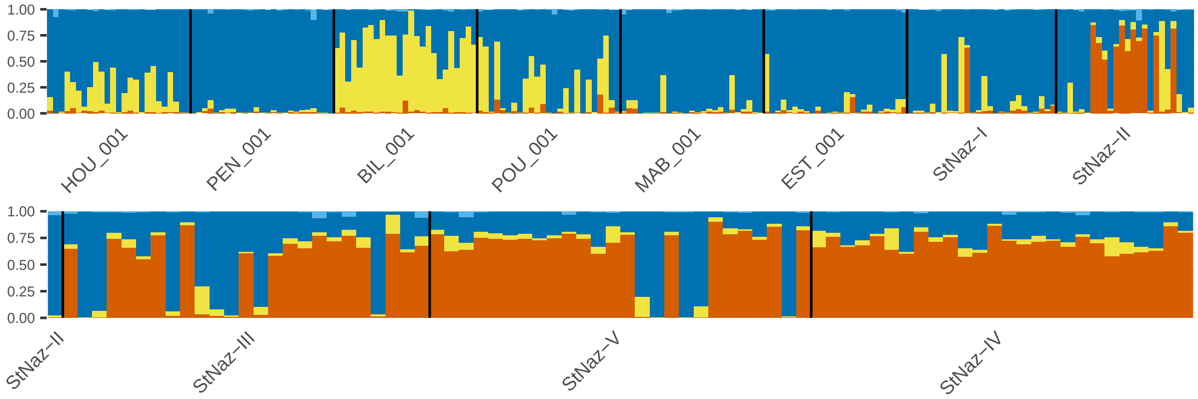

Figure S12: Subset of the port of Saint-Nazaire.

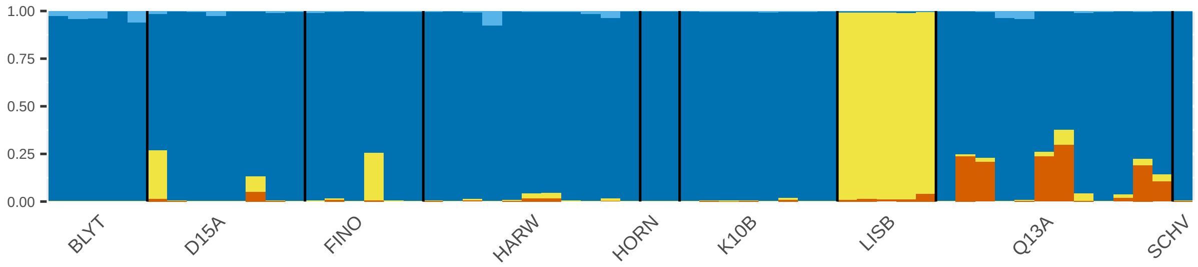

Figure S13: Subset of the populations studied in Coolen (2017).

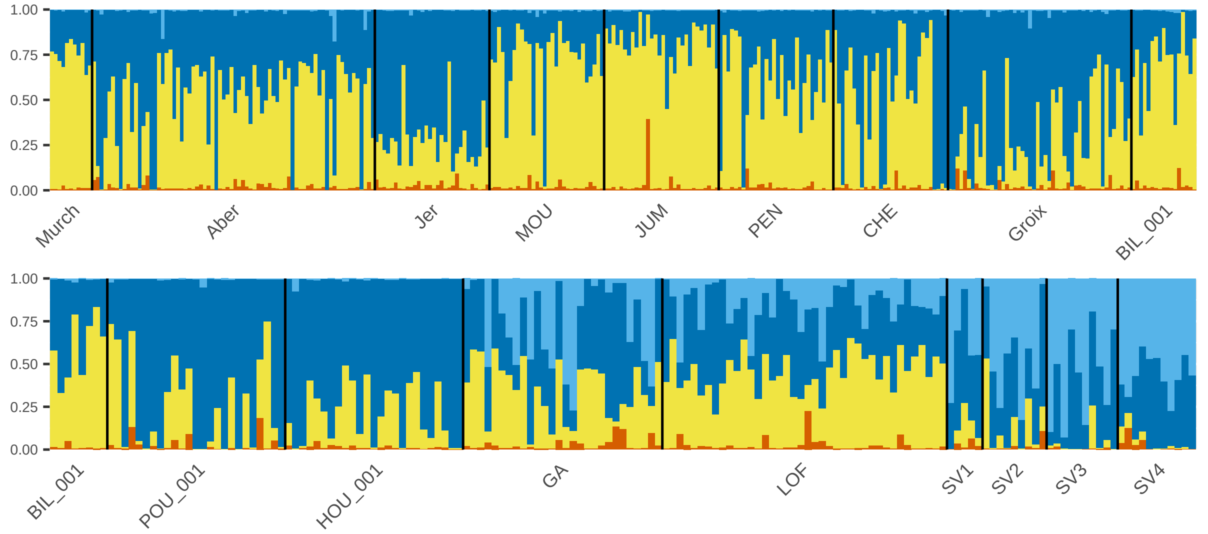

Figure S14: Subset of admixed populations with Atlantic *M. galloprovincialis* ancestries, including Natural admixed and Norway admixed. Populations LOF and SV1 to SV4 come from the Mathiesen et al. (2016) study.

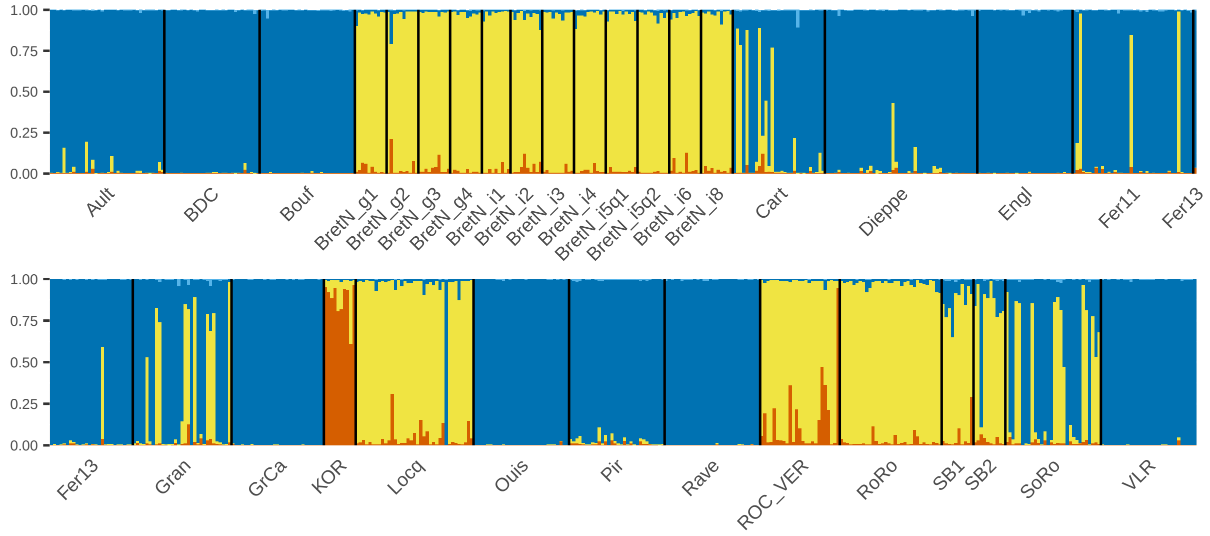

Figure S15: Subset of additional populations.

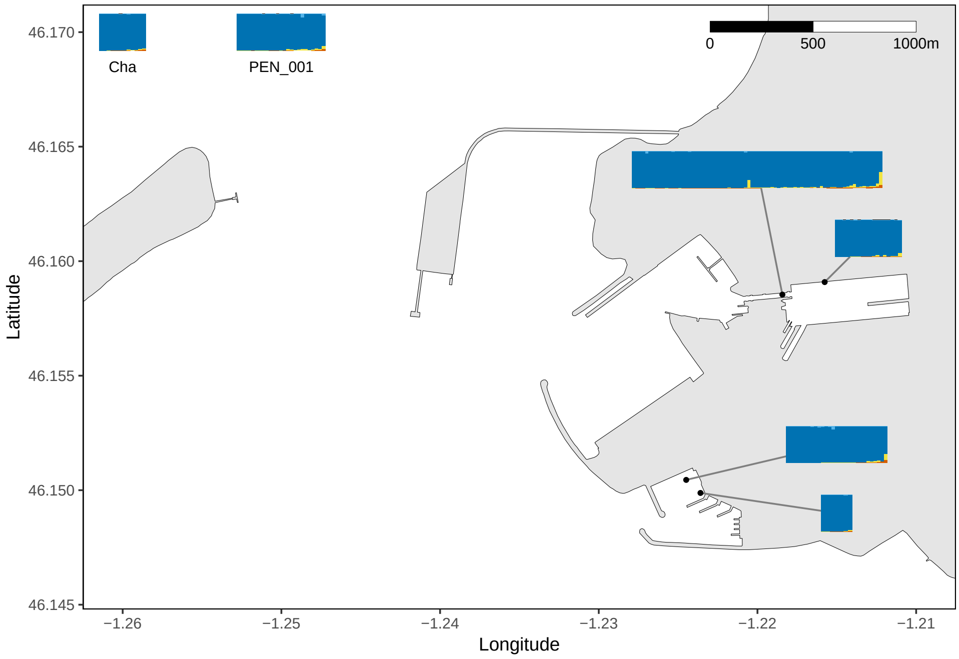

Figure S16: Map of the port of La Rochelle, bay of Biscay, France. Barplots represent individual ancestry proportions for each site. Barplots at the map edges correspond to distant populations. No Mediterranean *M. galloprovincialis* ancestry is detected in this port.

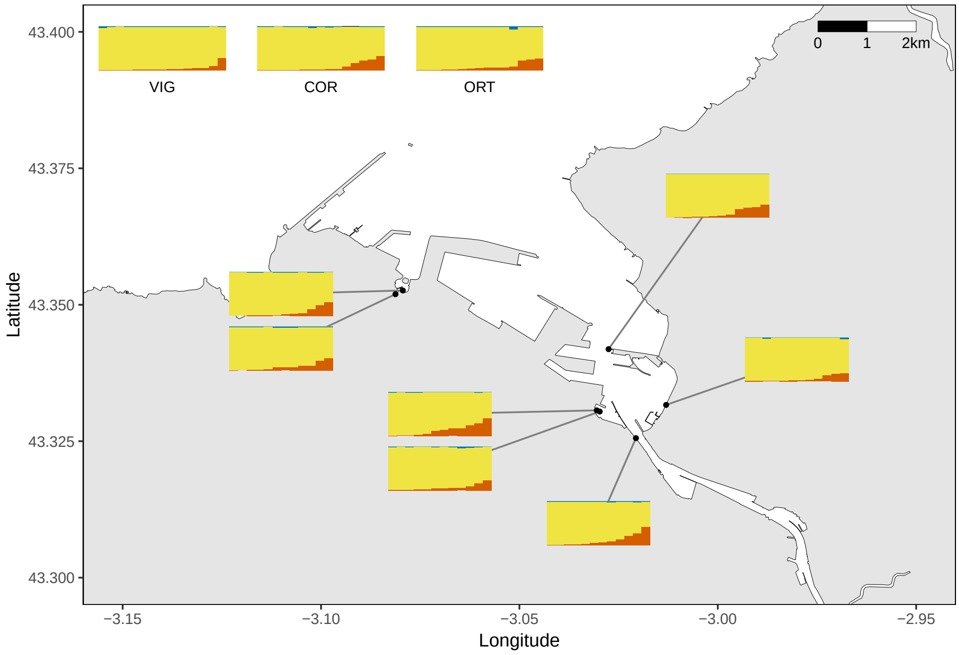

Figure S17: Map of the port of Bilbao, Basque Country, Spain. Sites inside the port do not exhibit different ancestriy compositions than more distant populations

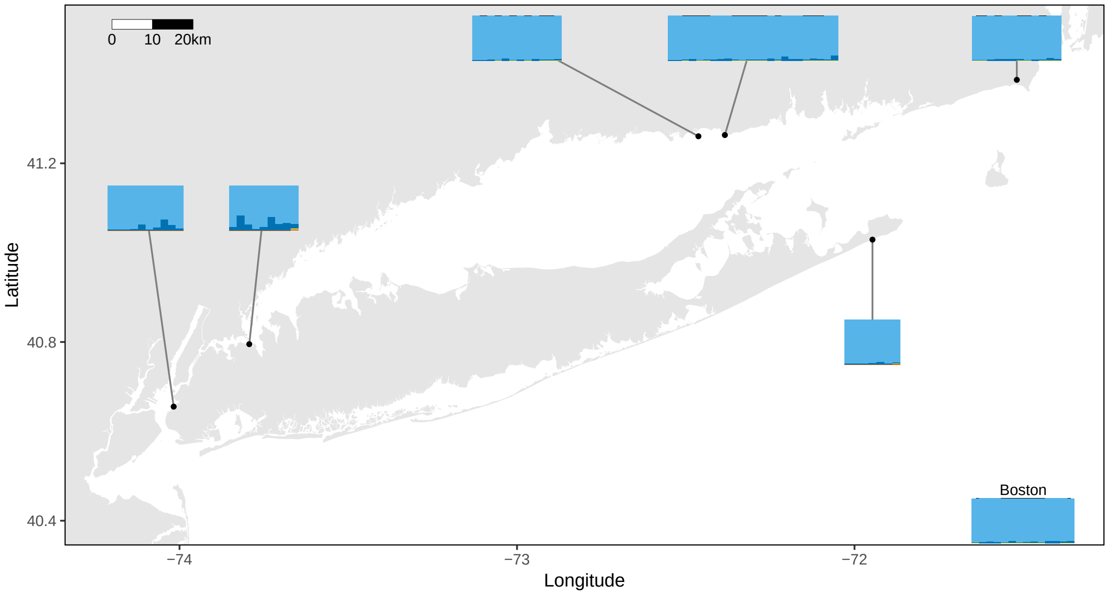

Figure S18: Map of the Long Island Sound, NY, USA. The two leftmost populations exhibit more European *M. edulis* ancestry than local American *M. edulis* populations.

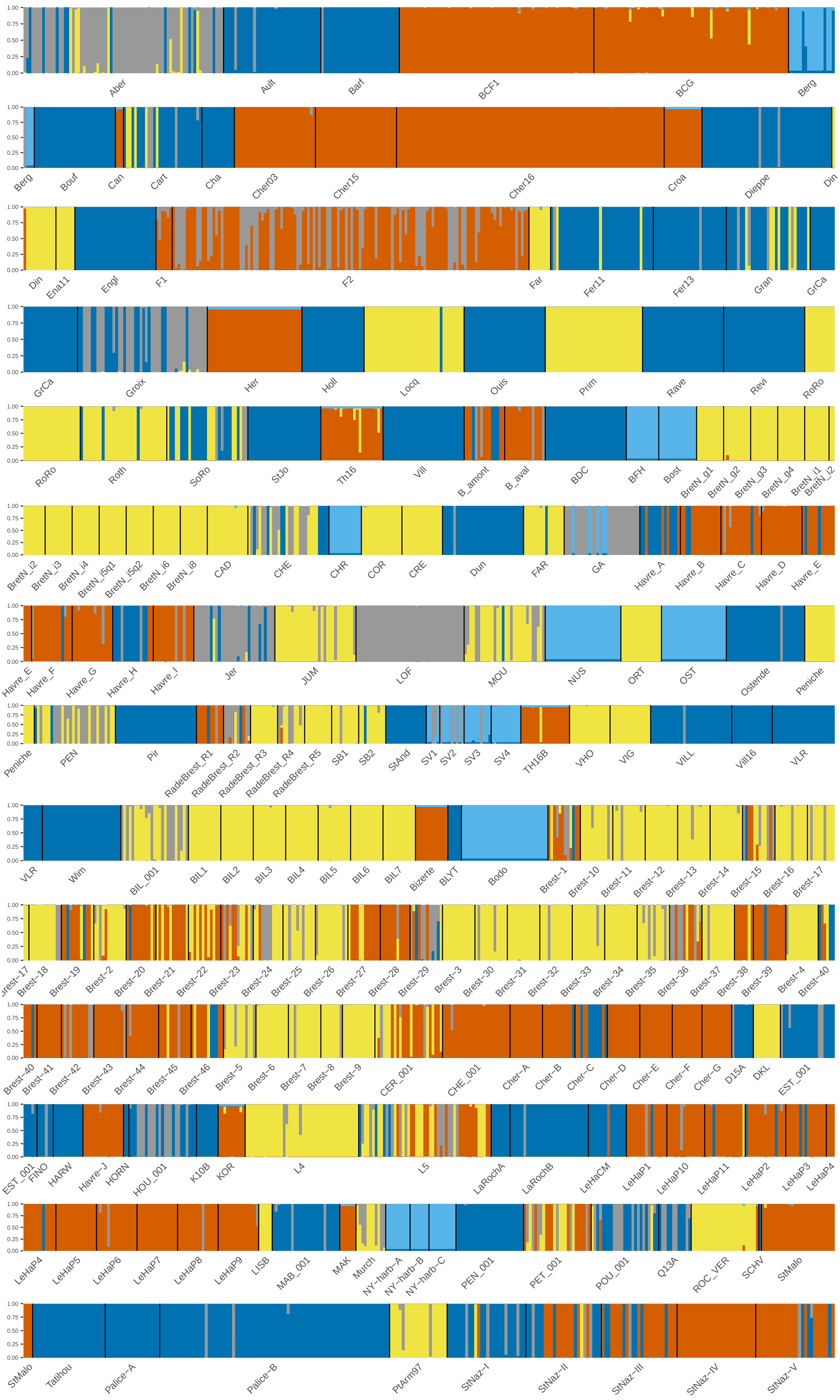

Figure S19: `Structure` without admixture model, $K = 5$

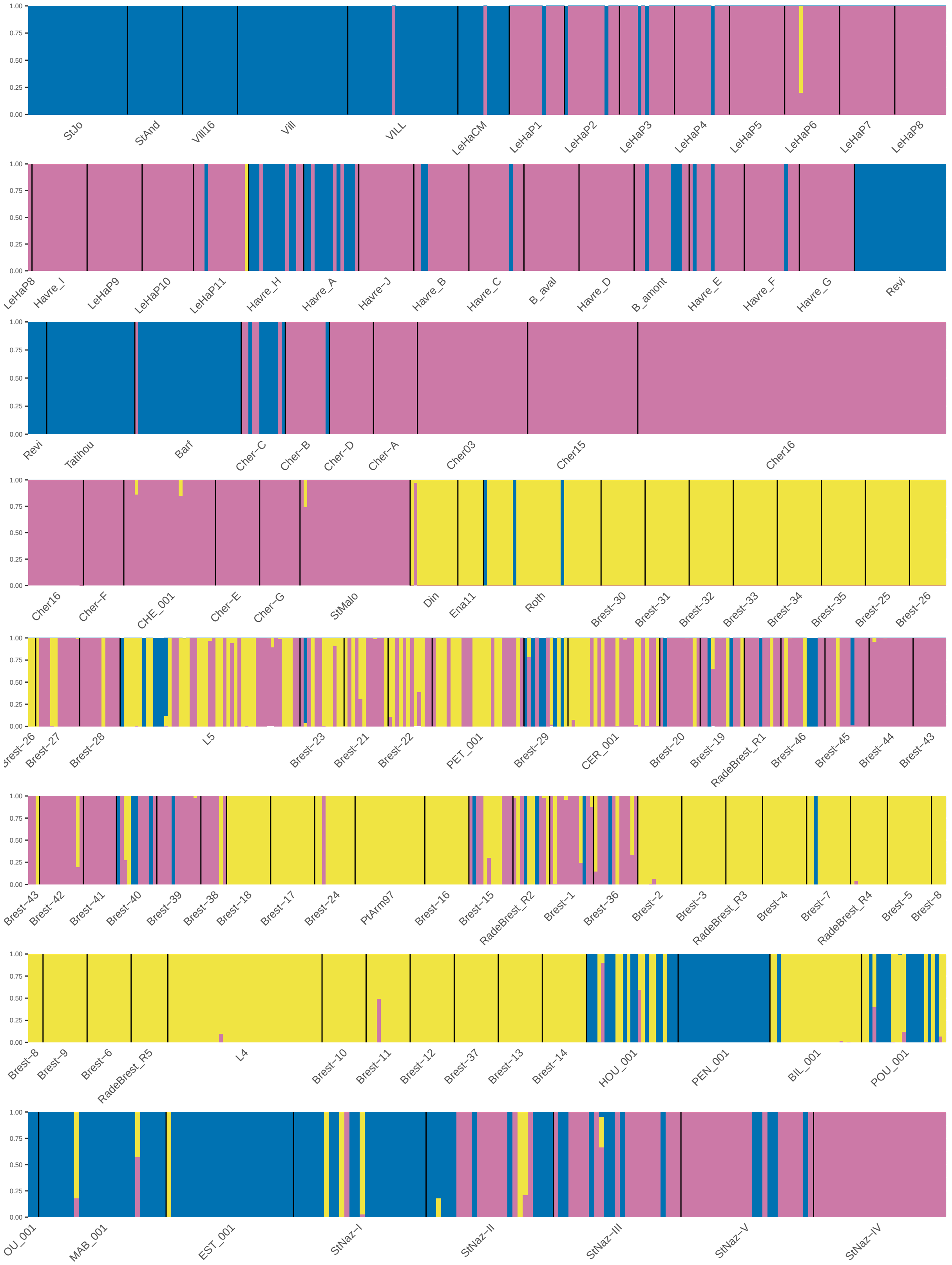

Figure S20: `Structure` without admixture model only with port and local populations, $K = 3$. This output is used to classify dock mussels for downstream analyses.

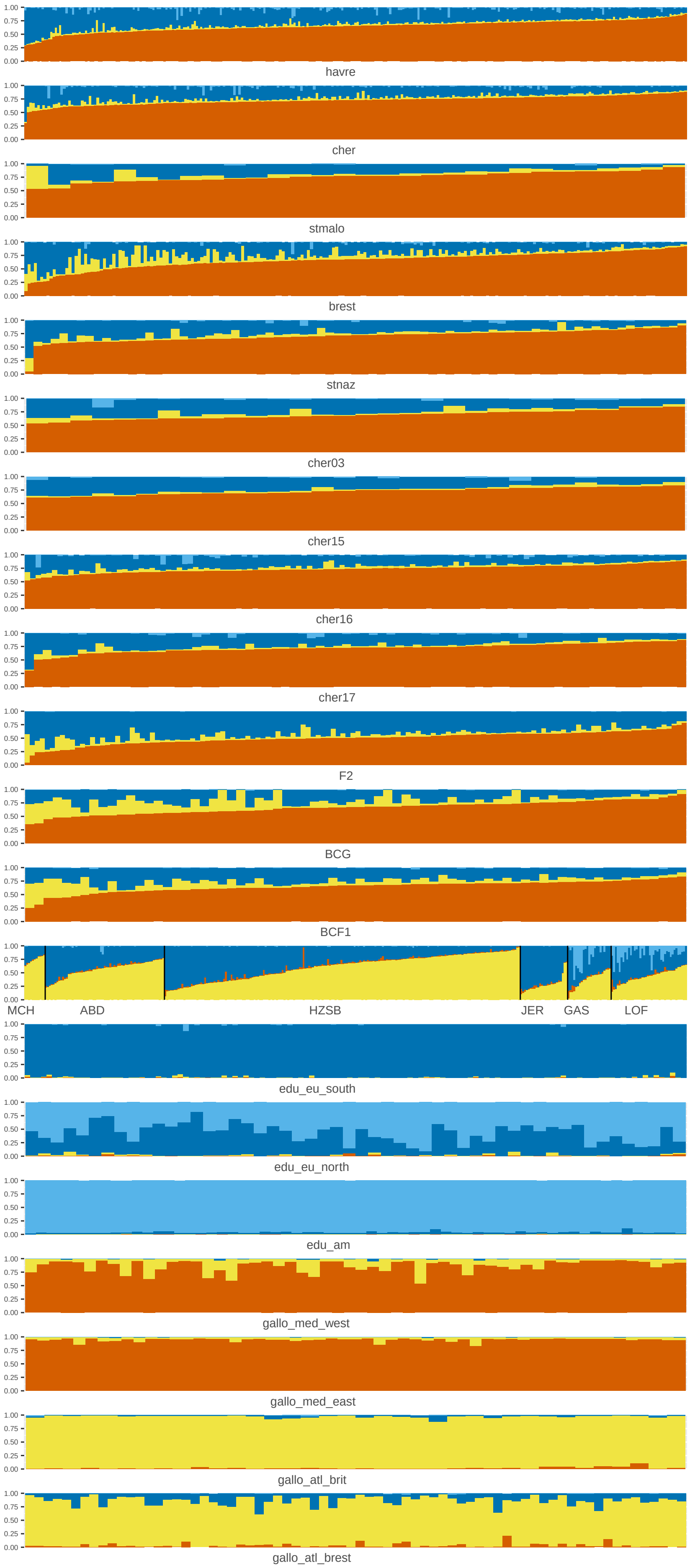

Figure S21: Structure plot of selected individuals for each group for downstream analyses

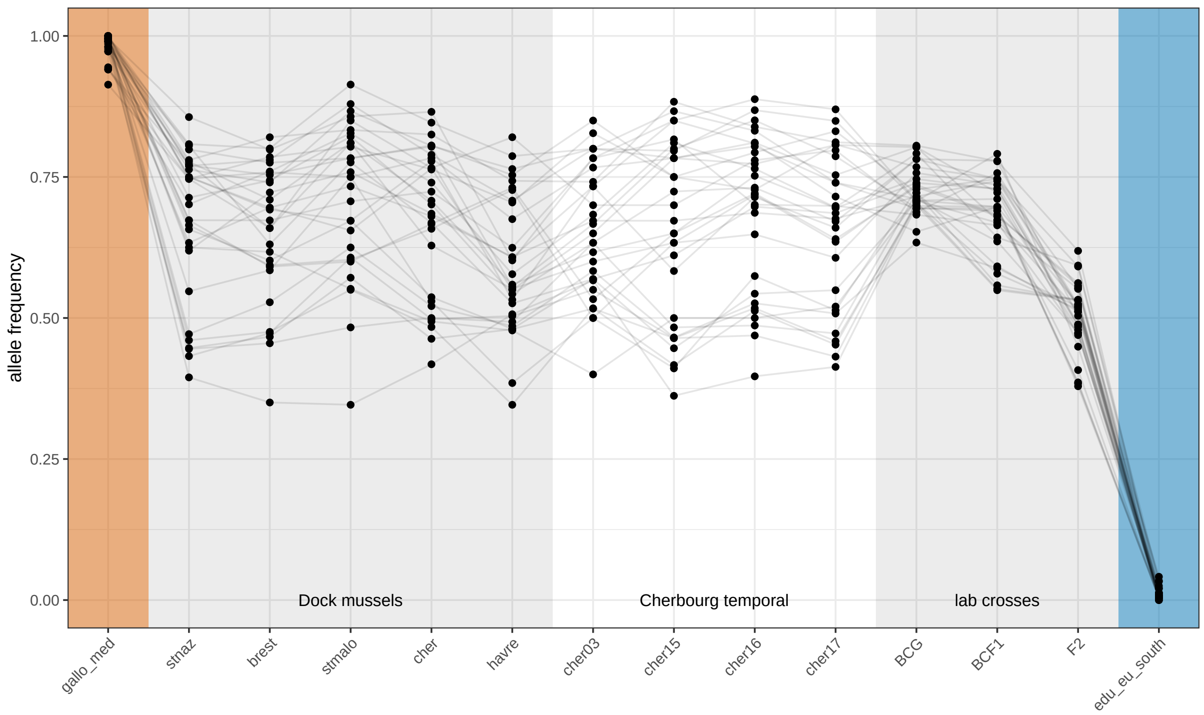

Figure S22: Allele frequencies of differentiated markers (AFD > 0.9) between Med. *M. galloprovincialis* and South-Eu. *M. edulis* in dock mussels, temporal samples of Cherbourg and laboratory crosses. Markers not heterozygous in the F1 cross were removed.

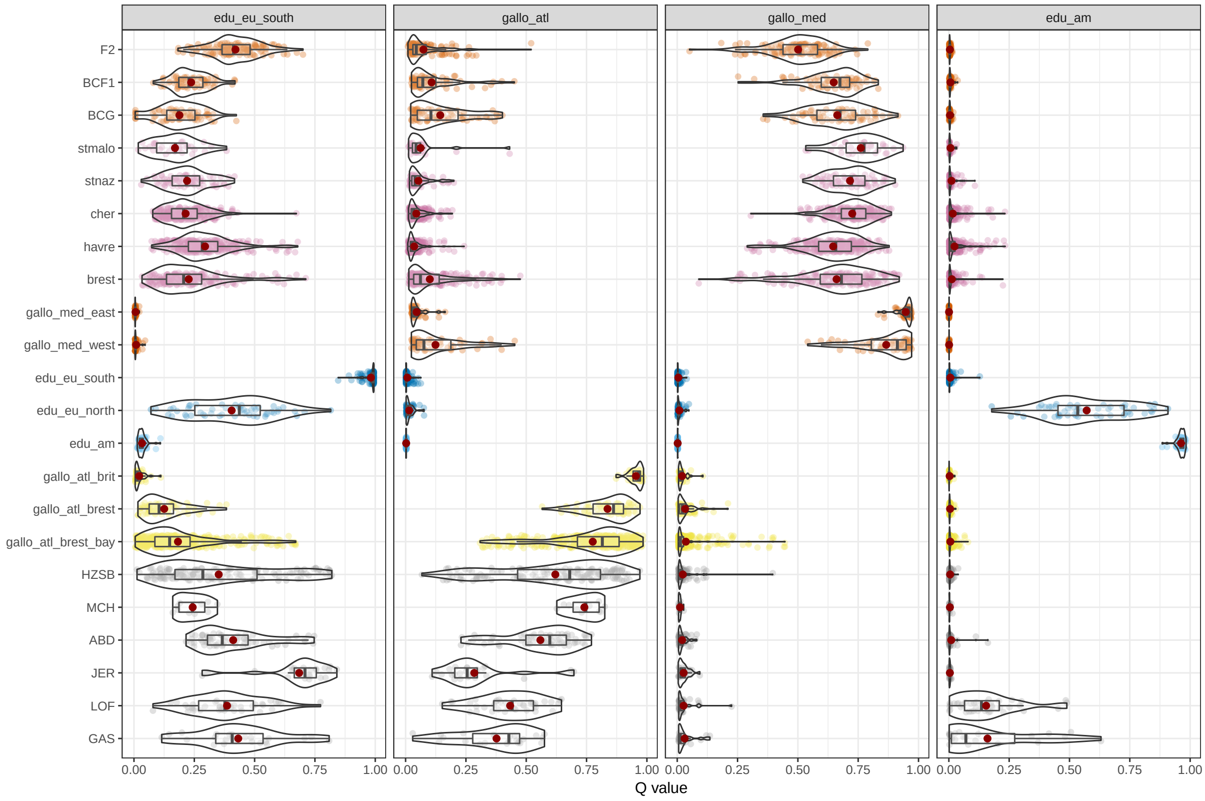

Figure S23: Ancestry levels for each `Structure` cluster in populations of admixed mussels and reference groups.

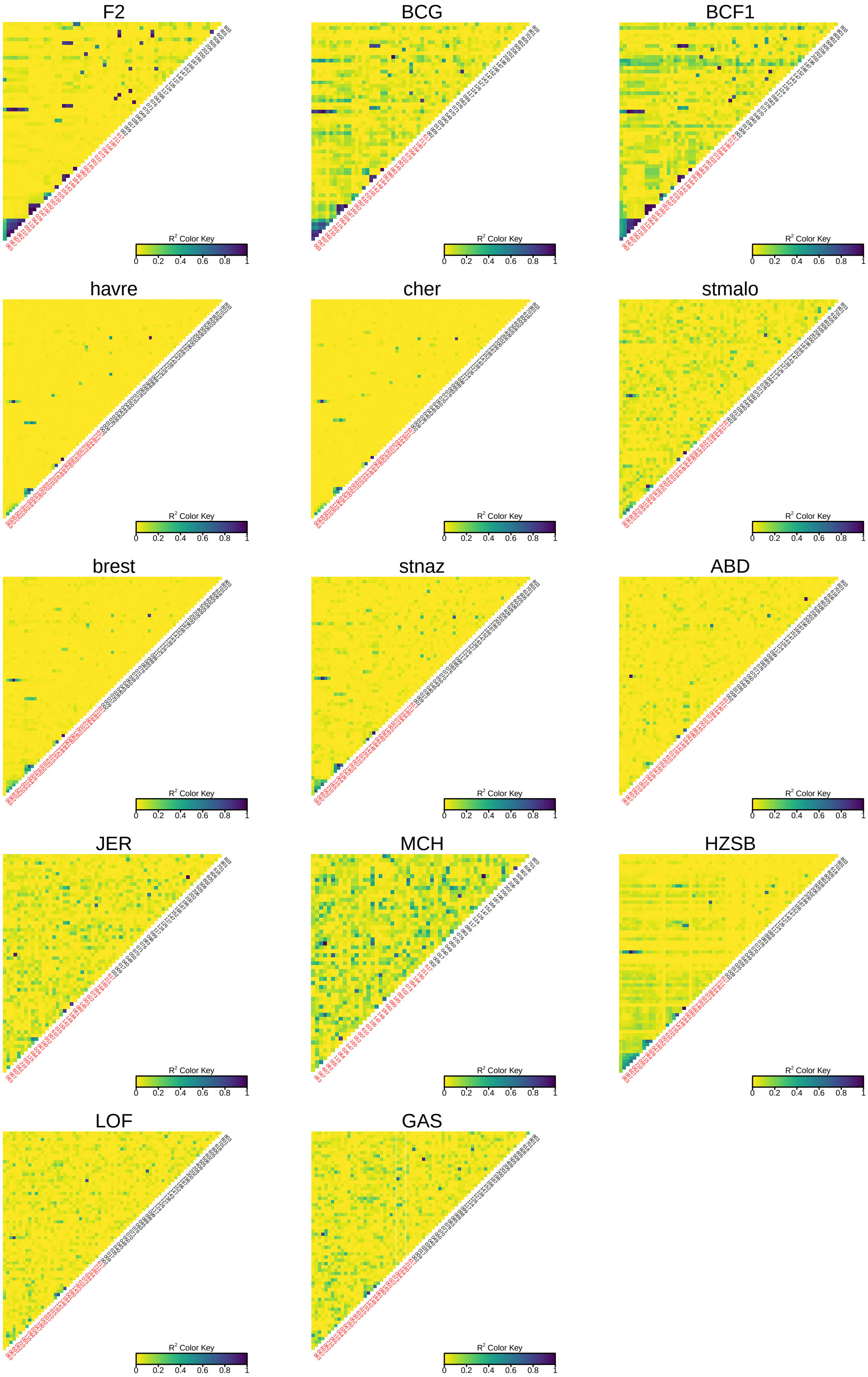

Figure S24: Linkage disequilibrium ($R^2$) in admixed populations. Markers are order following the genetic map (red labels) and then by name for loci that could be included in the genetic map construction.

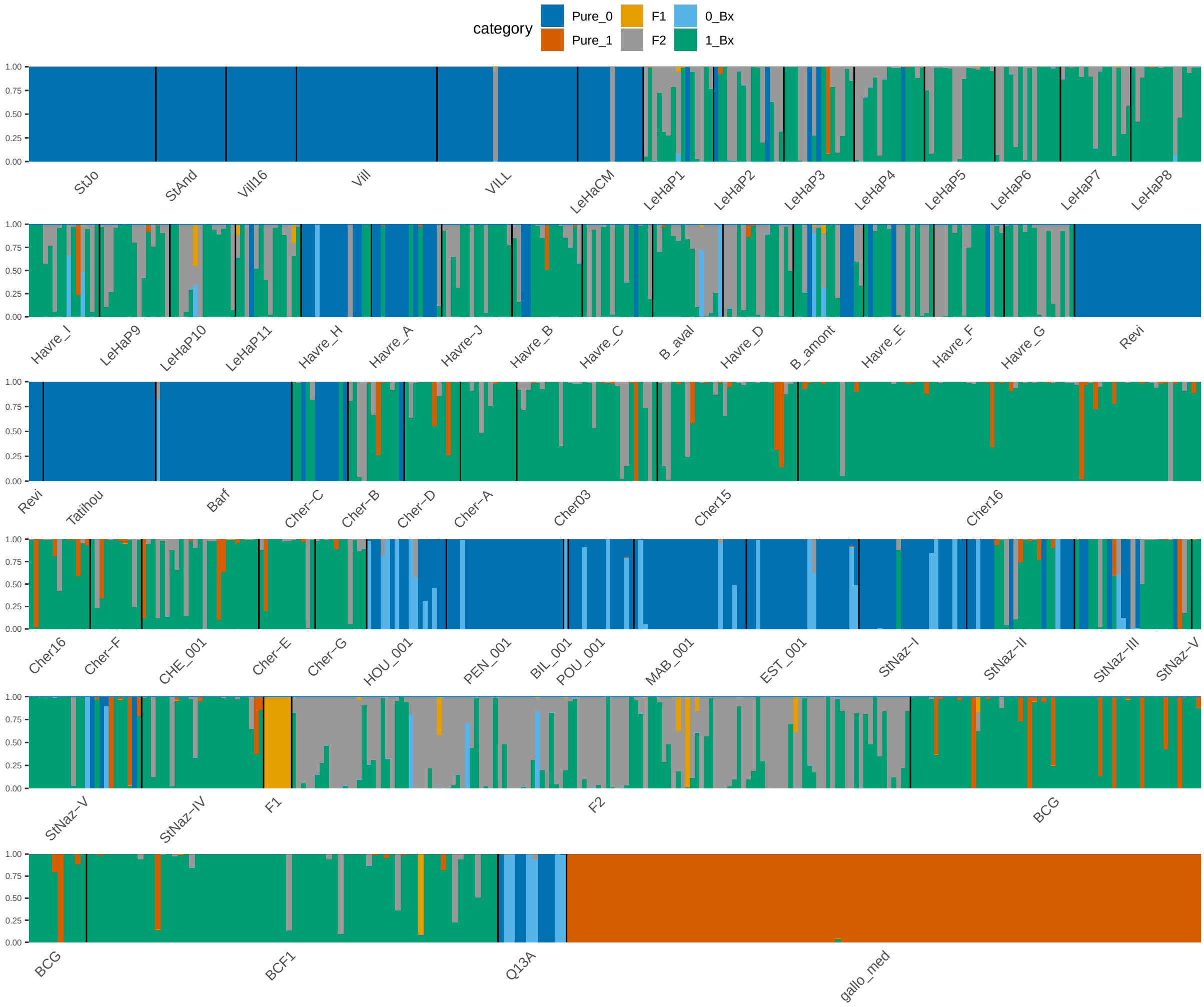

Figure S25: `Newhybrids` analysis including reference Mediterranean *M. galloprovincialis* (Pure_1) and local populations of all ports excluding Brest, containing South-Eu. *M. edulis* individuals (Pure_0). Lab crosses have been included and the results globally conform to the expected genotypes (F1, F2, BCG and BCF1). We hypothesise that pure Med. *M. galloprovincialis* assignements in ports are the result of marker sampling variance, as those assignments are also present in the lab BC samples.

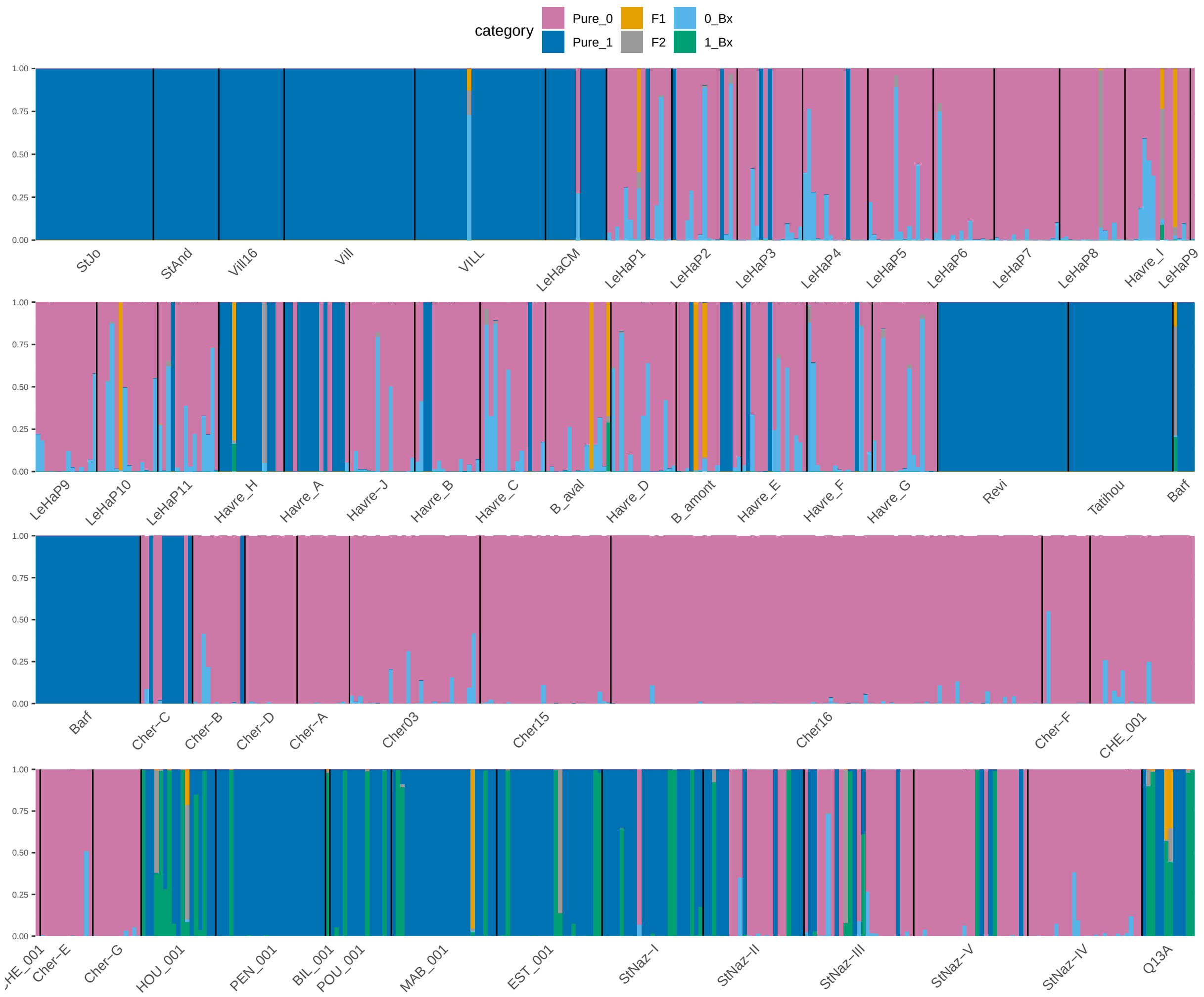

Figure S26: `Newhybrids` analysis comparing dock mussels (Pure_0) and the native Sout-Eu. *M. edulis* (Pure_1). In spite of the filtration to eliminate Atl. *M. galloprovincialis* genotypes, residual ancestry in individuals for populations around Saint-Nazaire (HOU_001 to StNaz-IV) produces spurious 1_Bx assignements.

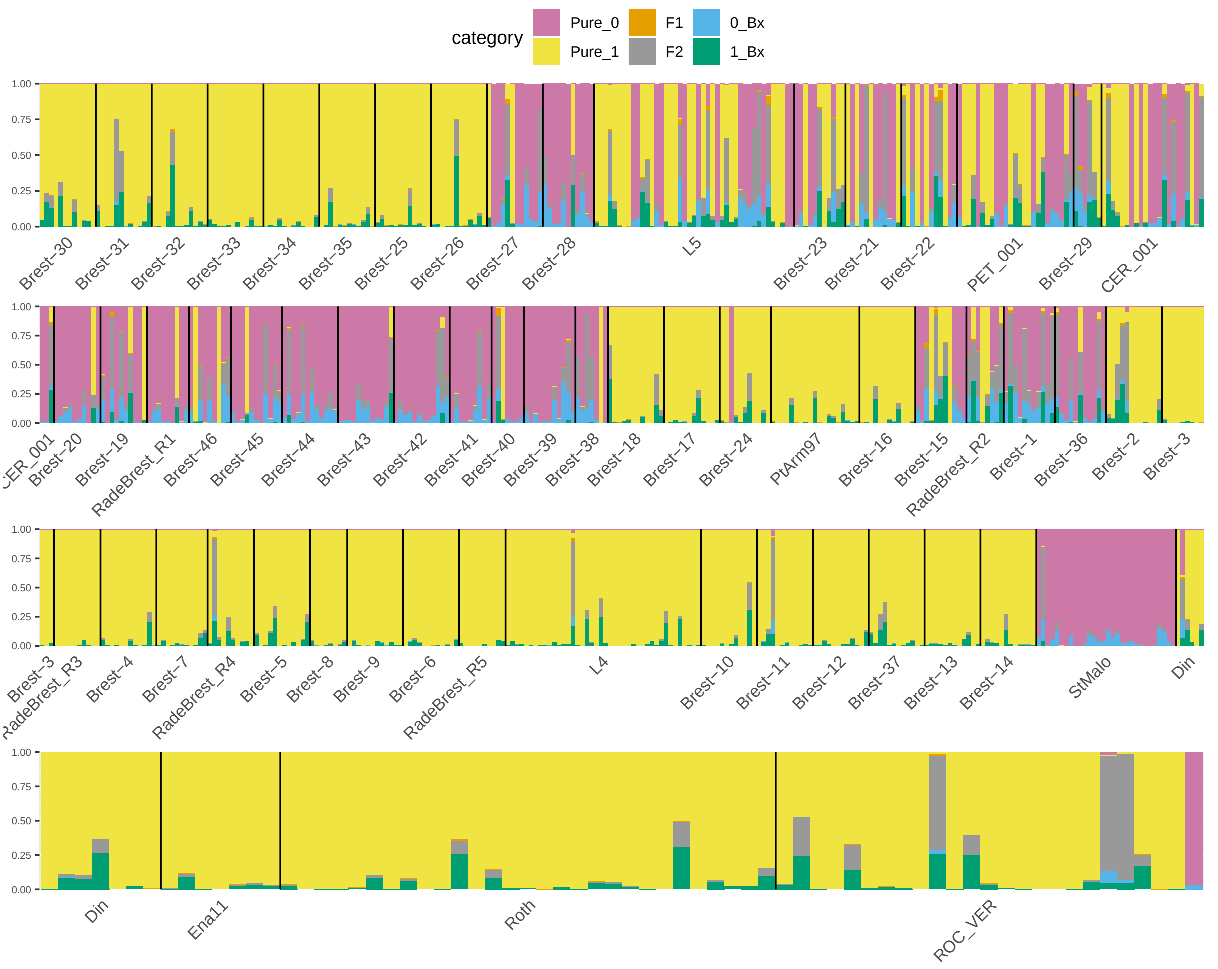

Figure S27: `Newhybrids` analysis comparing dock mussels (Pure_0) and native Atl. *M. galloprovincialis* (Pure_1).

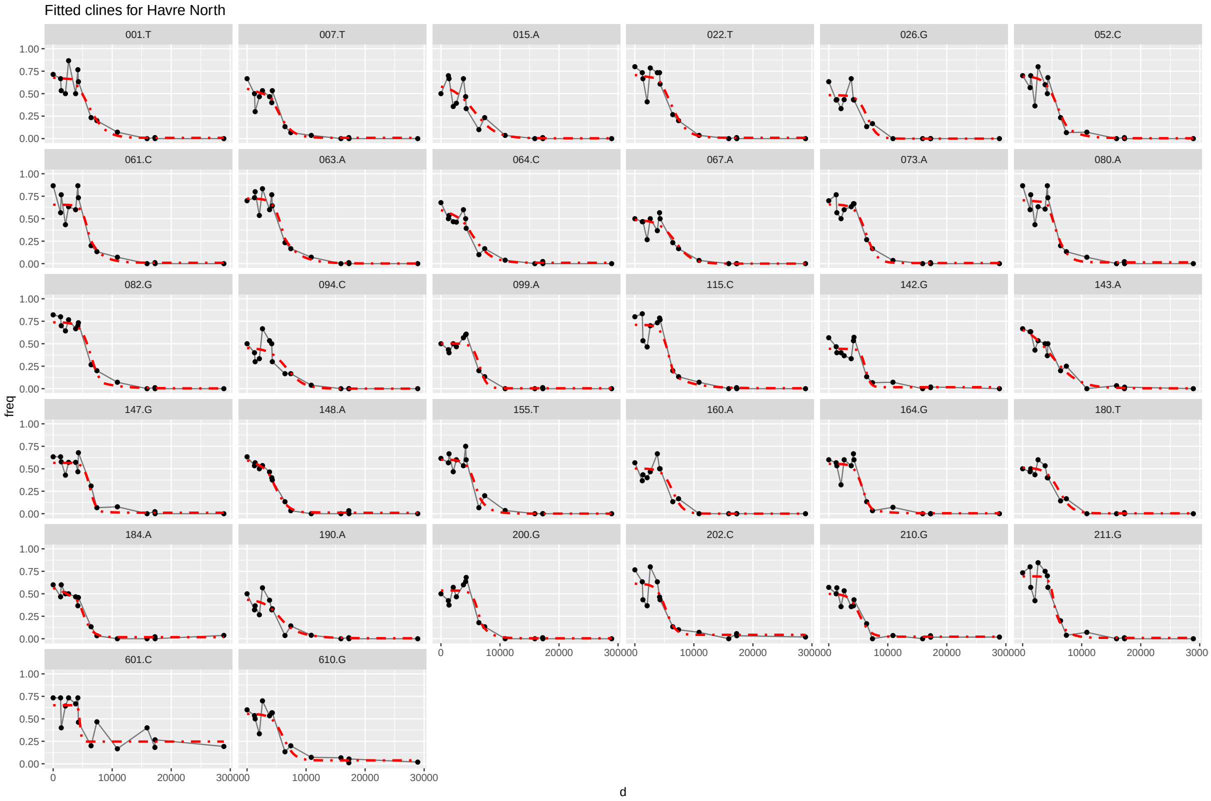

Figure S28: Details of geopgraphic clines per marker fitted in Le Havre North transect. Each point is the allele frequency of the Med. *M. galloprovincialis* allele at one site.

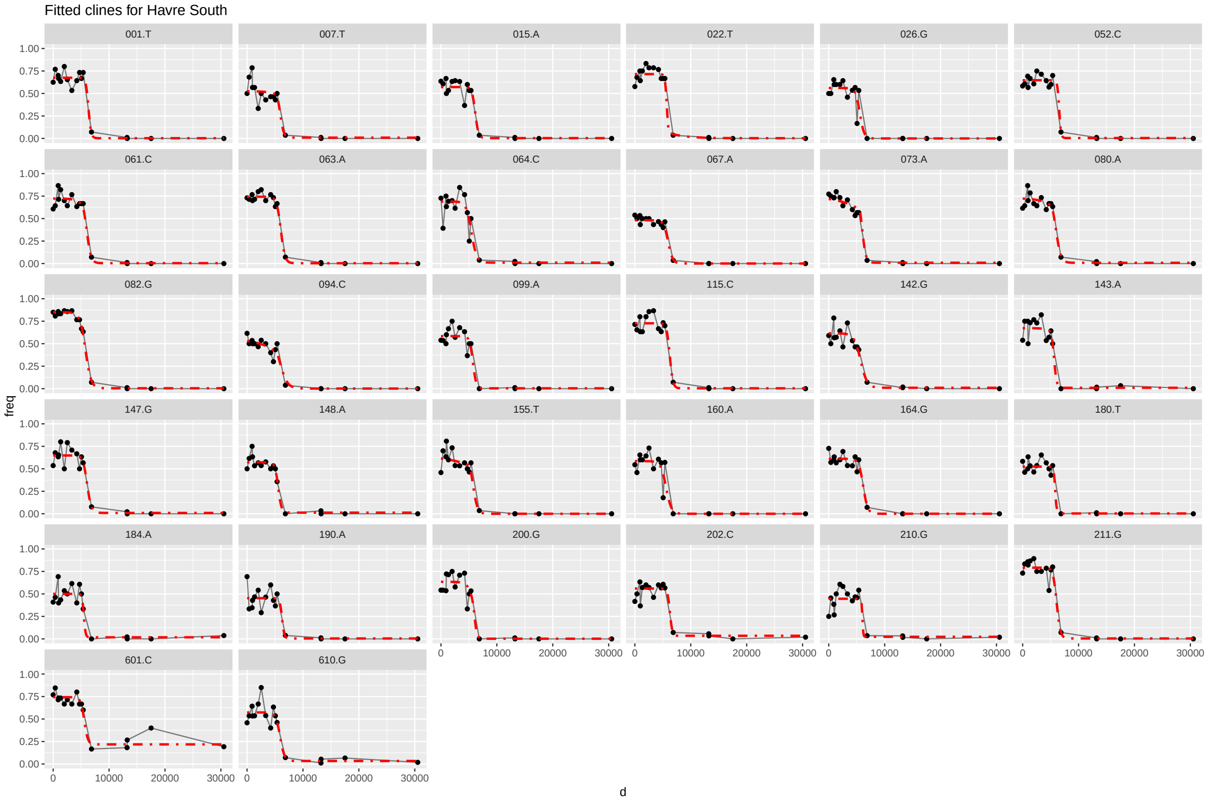

Figure S29: Details of geopgraphic clines per marker fitted in Le Havre South transect.

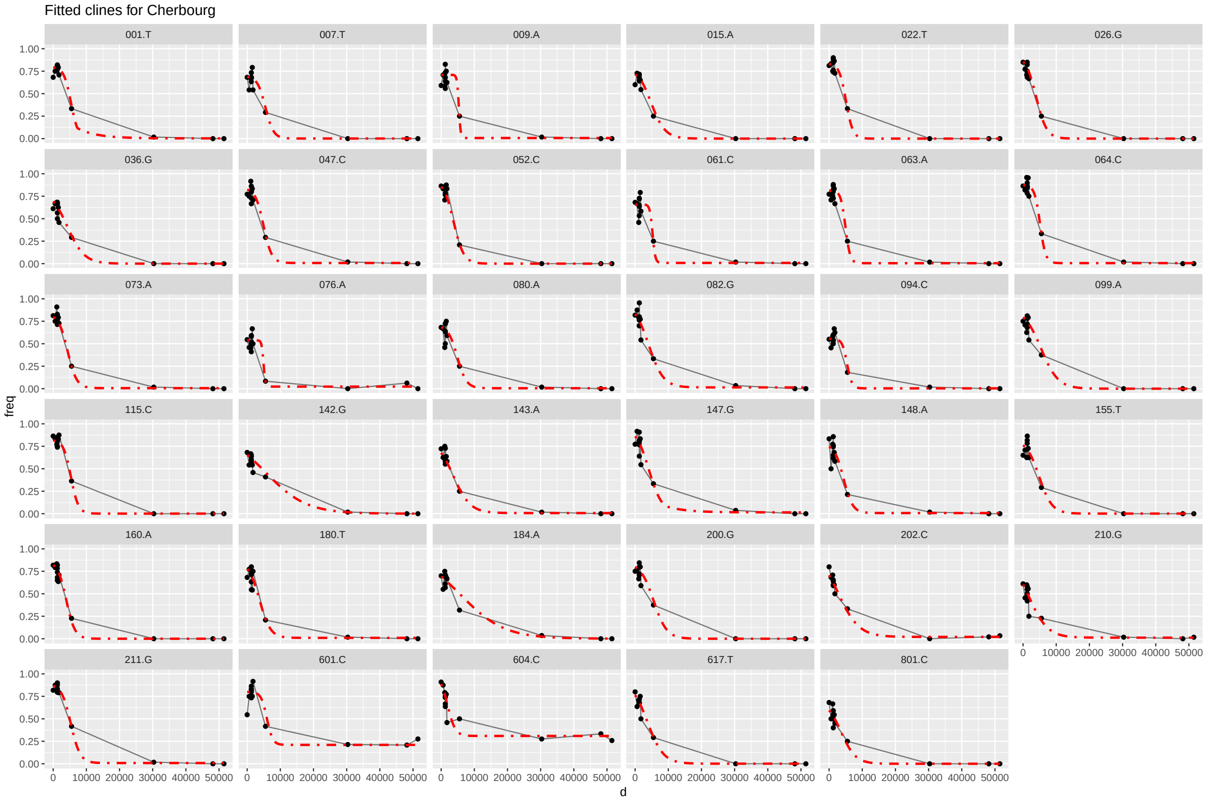

Figure S30: Details of geopgraphic clines per marker fitted in Cherbourg.

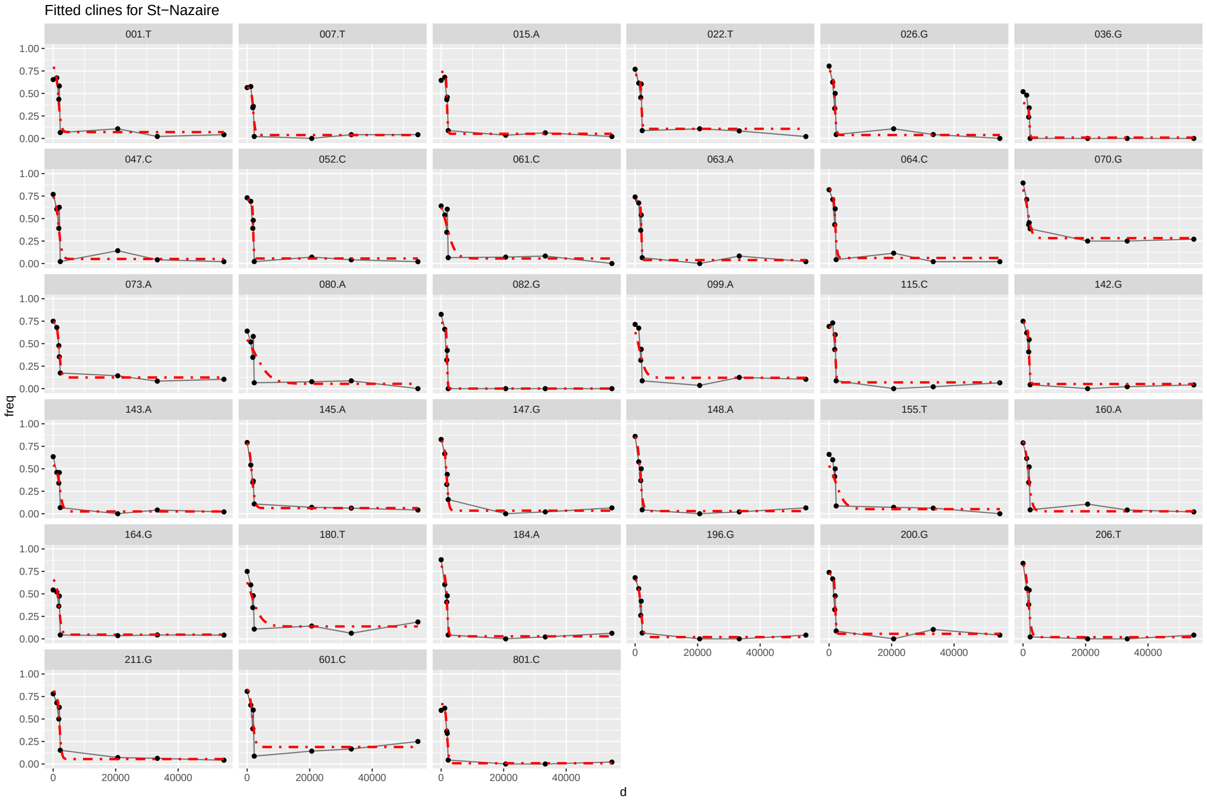

Figure S31: Details of geopgraphic clines per marker fitted in Saint-Nazaire.

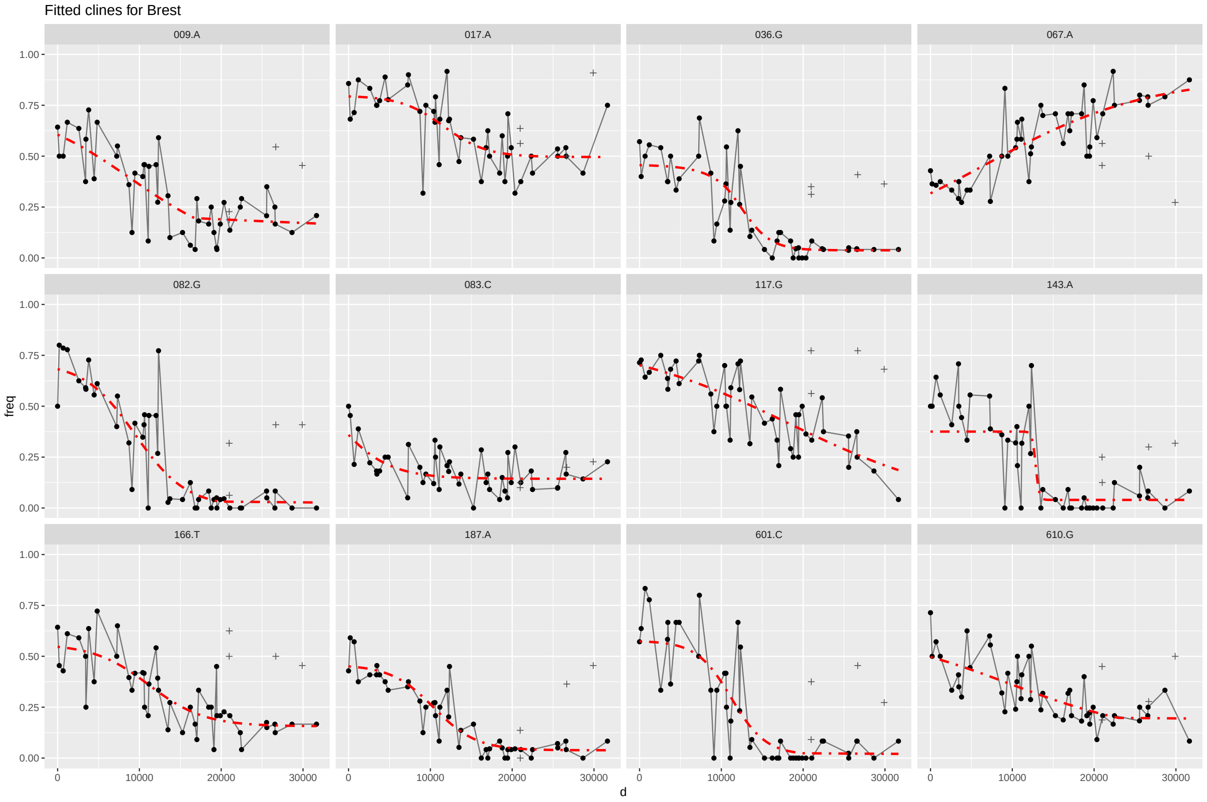

Figure S32: Details of geopgraphic clines per marker fitted in the bay of Brest. Crosses are sites not considered for the fit.

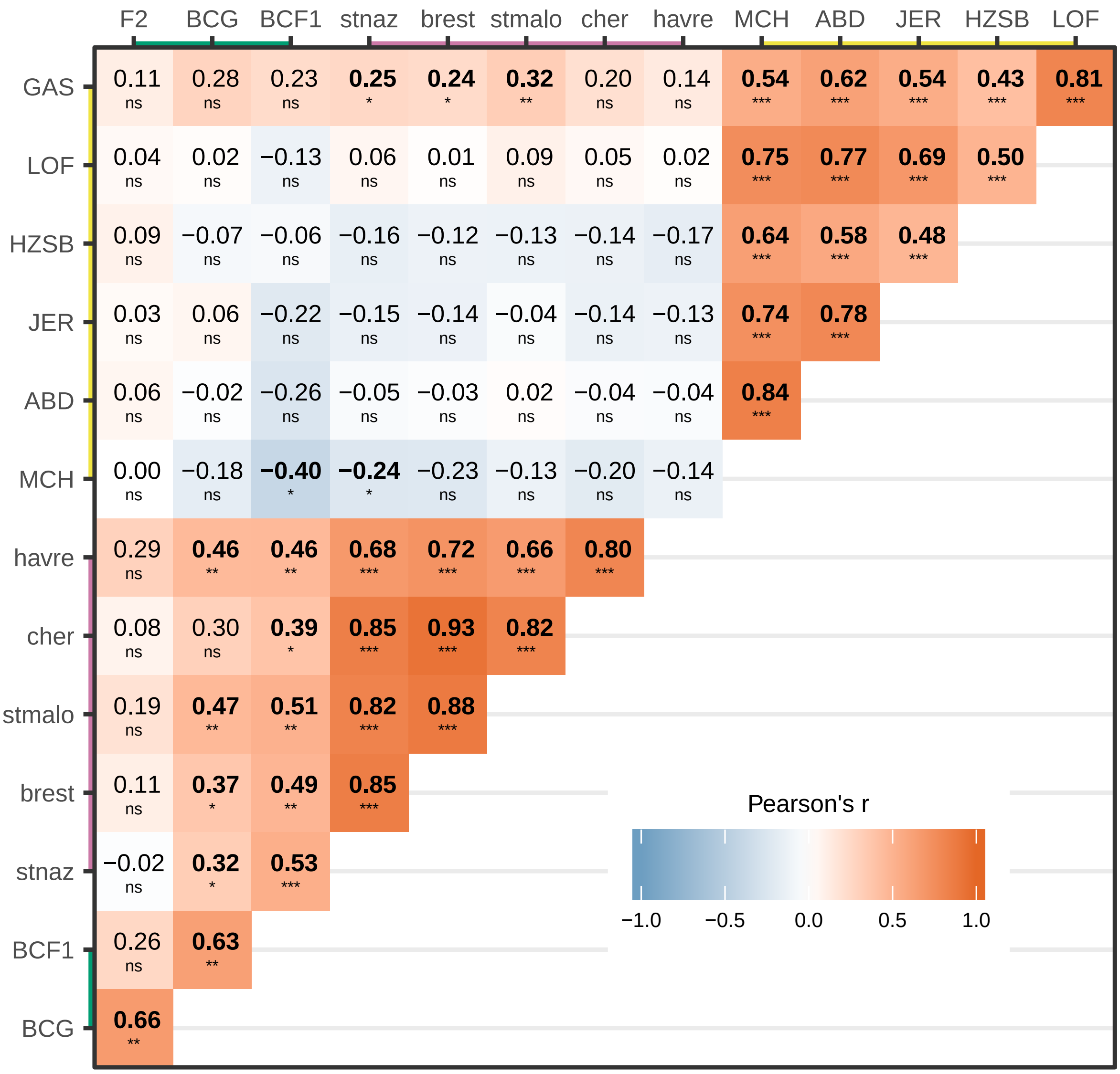

Figure S33: All correlations between populations.

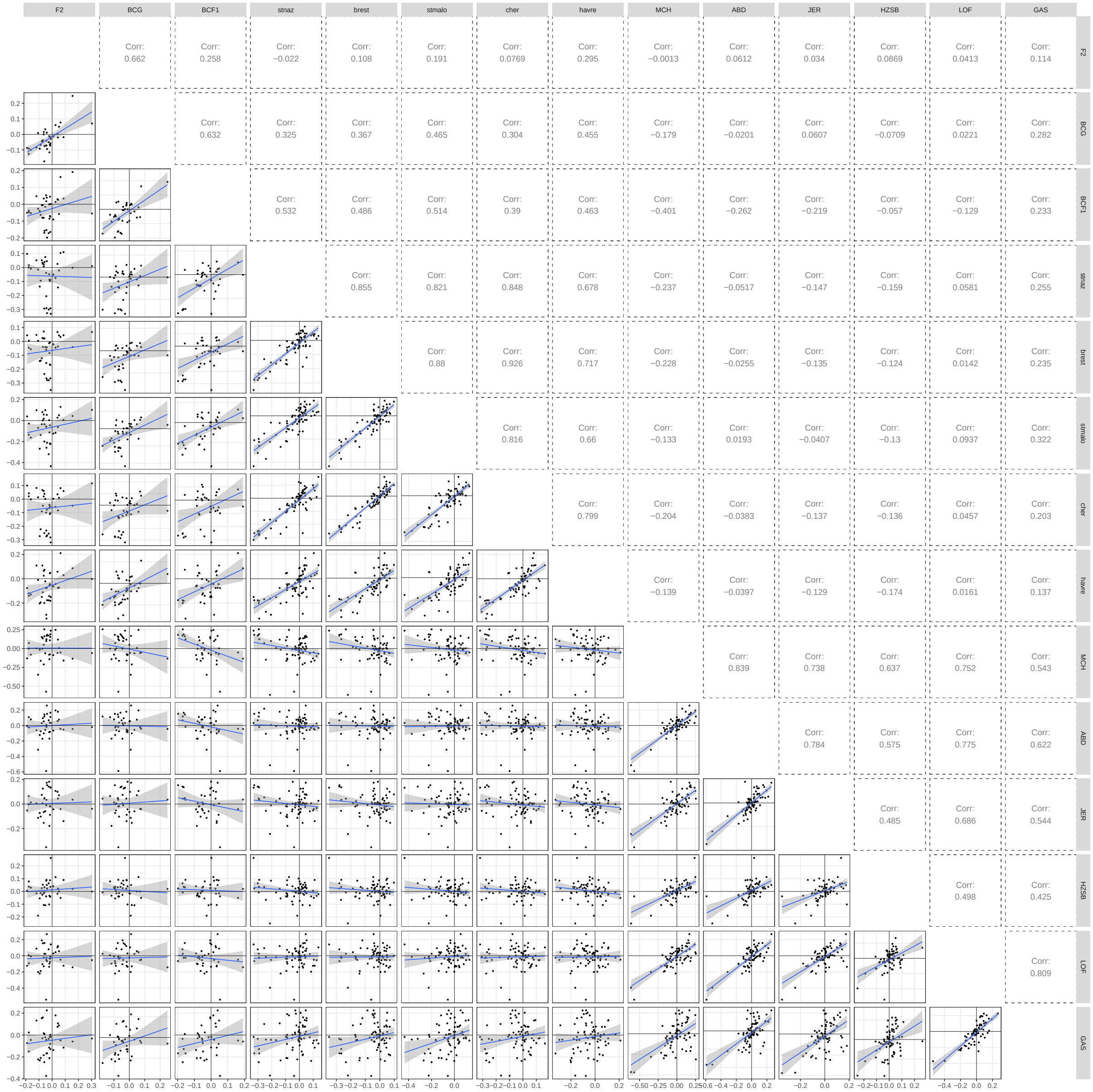

Figure S34: Scatter plots for all correlations between populations.

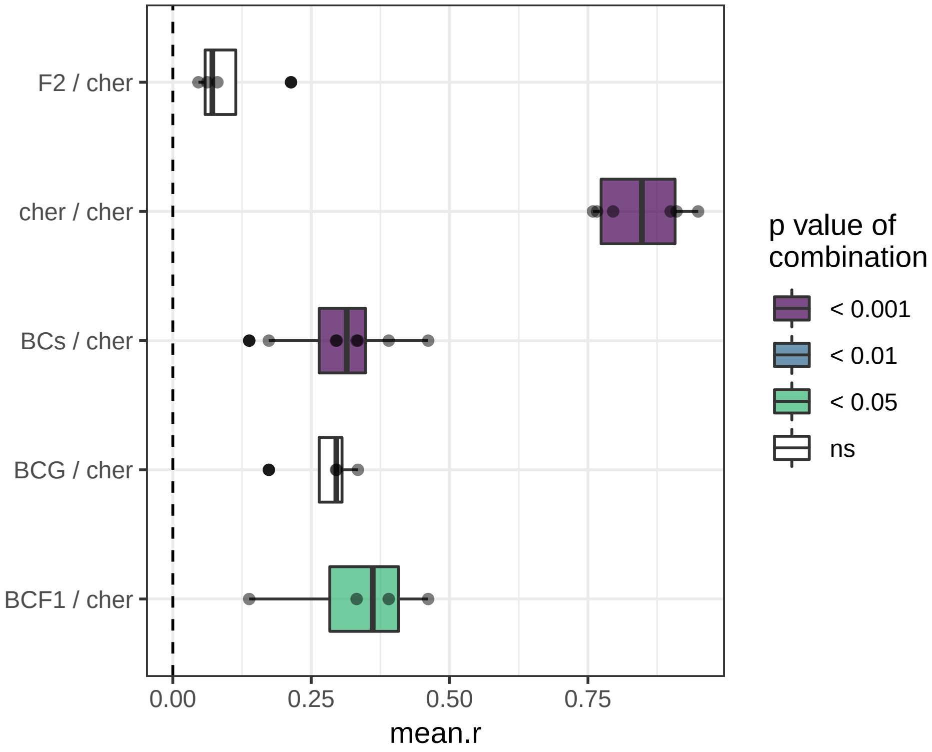

Figure S35: Correlation matrix for temporal variation in Cherbourg.

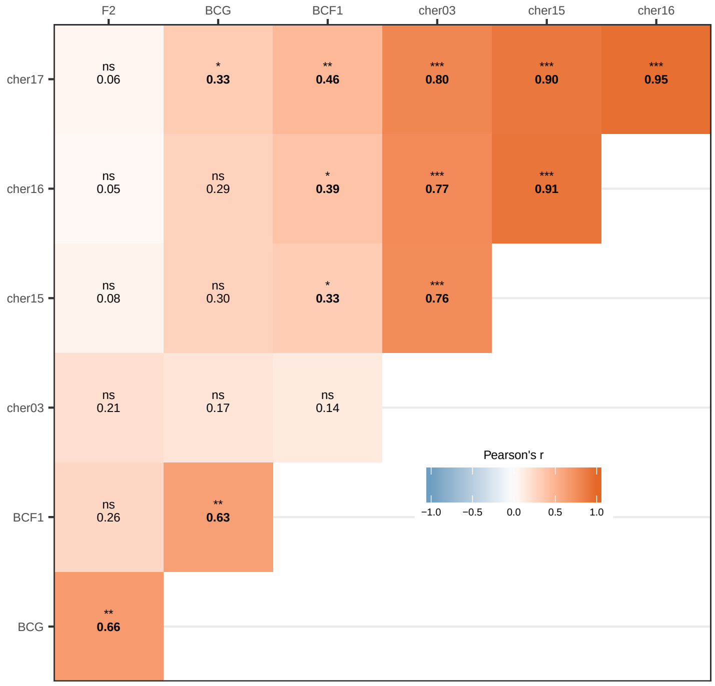

Figure S36: Correlation matrix for temporal variation in Cherbourg.

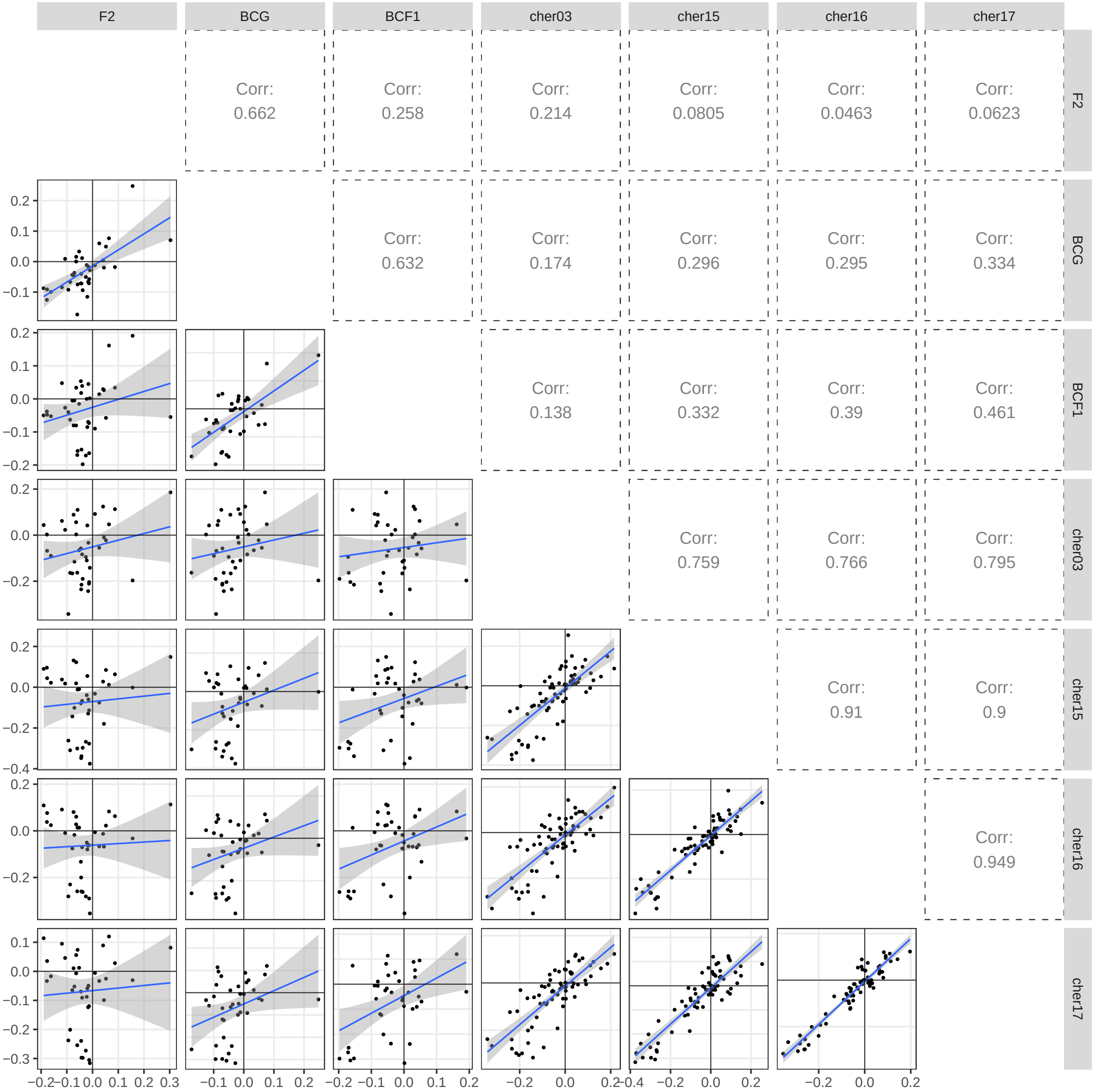

Figure S37: All correlations for Cherbourg.

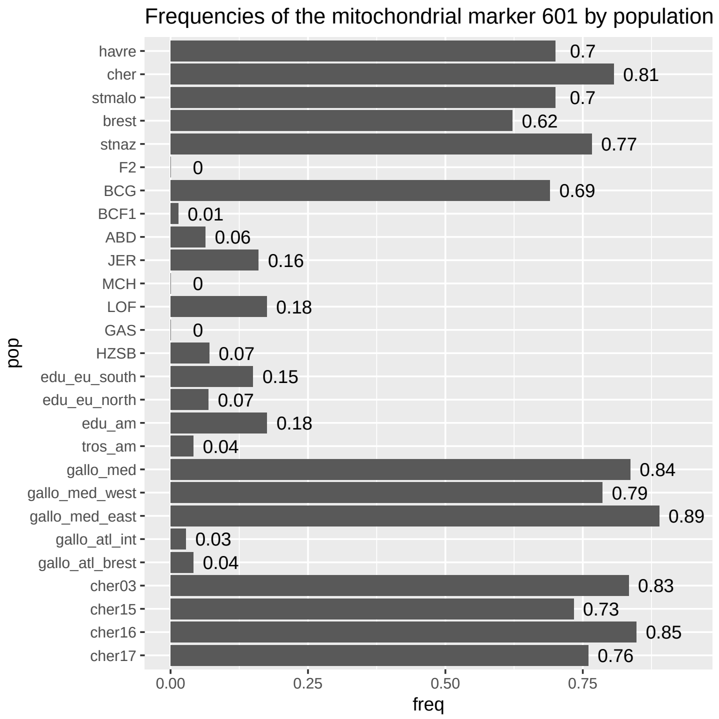

Figure S38: Allele frequencies of the female mitochondrial marker 601.

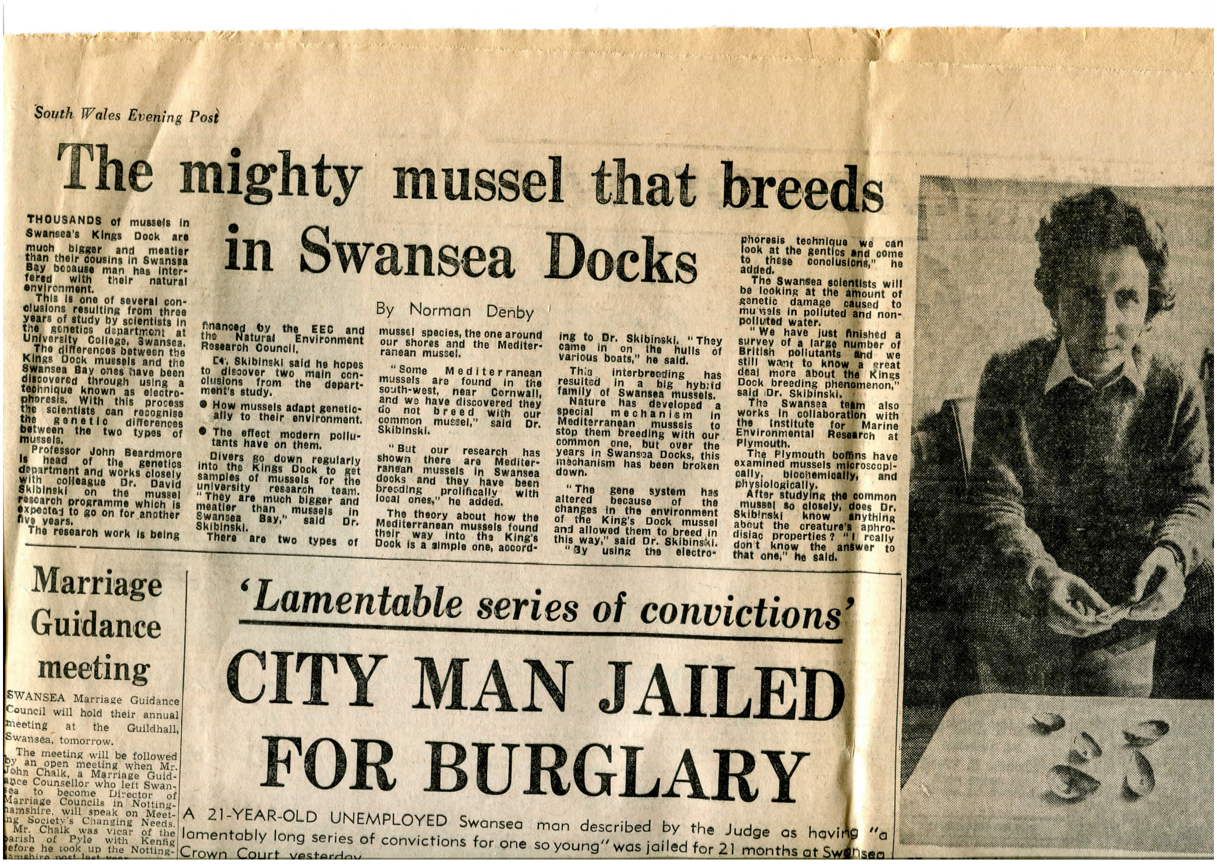

*South Wales Evening Post*

### The mighty mussel that breeds in Swansea Docks

By Norman Denby

THOUSANDS of mussels in Swansea's Kings Dock are much bigger and meatier than their cousins in Swansea Bay because man has interfered with their natural environment.

This is one of several conclusions resulting from three years of study by scientists in the genetics department at University College, Swansea.

The differences between the Kings Dock mussels and the Swansea Bay ones have been discovered through using a technique known as electrophoresis. With this process the scientists can recognise the genetic differences between the two types of mussels.

Professor John Beardmore is head of the genetics department and works closely with colleague Dr. David Skibinski on the mussel research programme which is expected to go on for another five years.

The research work is being financed by the EEC and the Natural Environment Research Council.

Dr. Skibinski said he hopes to discover two main conclusions from the department's study.

- How mussels adapt genetically to their environment.
- The effect modern pollutants have on them.

Divers go down regularly into the Kings Dock to get samples of mussels for the university research team. "They are much bigger and meatier than mussels in Swansea Bay," said Dr. Skibinski.

There are two types of mussel species, the one around our shores and the Mediterranean mussel.

"Some Mediterranean mussels are found in the south-west, near Cornwall, and we have discovered they do not breed with our common mussel," said Dr. Skibinski.

"But our research has shown there are Mediterranean mussels in Swansea docks and they have been breeding prolifically with local ones," he added.

The theory about how the Mediterranean mussels found their way into the King's Dock is a simple one, according to Dr. Skibinski. "They came in on the hulls of various boats," he said.

This interbreeding has resulted in a big hybrid family of Swansea mussels.

Nature has developed a special mechanism in Mediterranean mussels to stop them breeding with our common one, but over the years in Swansea Docks, this mechanism has been broken down.

"The gene system has altered because of the changes in the environment of the King's Dock mussel and allowed them to breed in this way," said Dr. Skibinski.

"By using the electrophoresis technique we can look at the gentics and come to these conclusions," he added.

The Swansea scientists will be looking at the amount of genetic damage caused to mussels in polluted and non-polluted water.

"We have just finished a survey of a large number of British pollutants and we still want to know a great deal more about the Kings Dock breeding phenomenon," said Dr. Skibinski.

The Swansea team also works in collaboration with the Institute for Marine Environmental Research at Plymouth.

The Plymouth boffins have examined mussels microscopically, biochemically, and physiologically.

After studying the common mussel so closely, does Dr. Skibinski know anything about the creature's aphrodisiac properties? "I really don't know the answer to that one," he said.

#### Marriage Guidance meeting

SWANSEA Marriage Guidance Council will hold their annual meeting at the Guildhall, Swansea, tomorrow.

The meeting will be followed by an open meeting when Mr. John Chalk, a Marriage Guidance Counsellor who left Swansea to become Director of Marriage Councils in Nottinghamshire, will speak on Meeting Society's Changing Needs.

Mr. Chalk was vicar of the parish of Pyle with Kenfig before he took up the Nottinghamshire post last year.

#### 'Lamentable series of convictions'

#### CITY MAN JAILED FOR BURGLARY

A 21-YEAR-OLD UNEMPLOYED Swansea man described by the Judge as having "a lamentably long series of convictions for one so young" was jailed for 21 months at Swansea Crown Court yesterday.

Figure S39: The mighty mussels: newspaper article about the Swansea King's dock mussels and the work of Prof. David Skibinski. South Wales Evening Post, 28th of June 1978.

### 3 Supplementary Tables

Table S1: Population information with sample size after filtration and average missing data among population.

| pop | longitude | latitude | locality | date | N | missing data |
|---|---|---|---|---|---|---|
| Aber | -2.07349 | 57.14758 | ABD, Aberdeen, Scotland, UK | | 74 | 0.0254 |
| Ault | 1.42314 | 50.09058 | Ault, France | | 36 | 0.0087 |
| B_amont | 0.14587 | 49.47124 | Bassin Amont, Le Havre, France | | 15 | 0.0346 |
| B_aval | 0.13242 | 49.47447 | Bassin Aval, Le Havre, France | | 15 | 0.0165 |
| Barf | -1.25807 | 49.71015 | Barfleur, France | 2015 | 29 | 0.0152 |
| BCF1 | | | Backcross | | 72 | 0.0253 |
| BCG | | | Backcross | | 72 | 0.0130 |
| BDC | 1.42282 | 50.09060 | Bois-de-Cise, France | 2016 | 30 | 0.0056 |
| Berg | 5.30573 | 60.39560 | Bergen, Norway | | 21 | 0.0235 |
| BFH | -72.46253 | 41.26056 | Brandford Harbor, USA | 2009 | 12 | 0.0162 |
| BIL_001 | -2.49015 | 47.44528 | Le Bile - Penestin, France | 11/01/17 | 25 | 0.0192 |
| BIL1 | -3.08124 | 43.35194 | Zierbena, Bilbao harbor, Spain | 17/07/17 | 12 | 0.0097 |
| BIL2 | -3.07939 | 43.35262 | Zierbena, Bilbao harbor, Spain | 17/07/17 | 12 | 0.0141 |
| BIL3 | -3.03050 | 43.33065 | Santurtzi, Bilbao harbor, Spain | 17/07/17 | 12 | 0.0097 |
| BIL4 | -3.02976 | 43.33044 | Santurtzi, Bilbao harbor, Spain | 17/07/17 | 12 | 0.0152 |
| BIL5 | -3.02063 | 43.32557 | Portugalete, Bilbao harbor, Spain | 17/07/17 | 12 | 0.0249 |
| BIL6 | -3.01302 | 43.33165 | Getxo, Bilbao harbor, Spain | 20/07/17 | 12 | 0.0119 |
| BIL7 | -3.02753 | 43.34187 | Getxo, Bilbao harbor, Spain | 20/07/17 | 12 | 0.0097 |
| Bizerte | 9.86000 | 37.26000 | Bizerte, Tunisia | | 12 | 0.0357 |
| BLYT | -1.49830 | 55.12580 | Blythe, North Sea | 14/11/14 | 5 | 0.0961 |
| Bodo | 14.99857 | 67.21838 | Bodø, Norway | 16/11/16 | 38 | 0.0308 |
| Bost | -71.05000 | 42.36000 | Boston, USA | 2009 | 14 | 0.0427 |
| Bouf | -0.72468 | 49.34802 | Le Bouffay, France | 2015 | 30 | 0.0156 |
| Brest-1 | -4.28628 | 48.31669 | Lagonna Daoulas, France | 08/07/17 | 12 | 0.0238 |
| Brest-10 | -4.58798 | 48.27521 | Fishing dock, Camaret, France | 09/07/17 | 12 | 0.0130 |
| Brest-11 | -4.59713 | 48.27972 | Camaret marina, France | 09/07/17 | 12 | 0.0162 |
| Brest-12 | -4.61044 | 48.26152 | Veryac'h, France | 09/07/17 | 12 | 0.0195 |
| Brest-13 | -4.49673 | 48.22357 | Morgat marina, France | 09/07/17 | 12 | 0.0087 |
| Brest-14 | -4.46343 | 48.23806 | Postolonnec beach, France | 09/07/17 | 12 | 0.0087 |
| Brest-15 | -4.36913 | 48.33790 | Porz Tinduff, France | 24/07/17 | 12 | 0.0173 |

*Continued on next page*

Table S1 – *Continued from previous page*

| pop | longitude | latitude | locality | date | N | missing data |
|---|---|---|---|---|---|---|
| Brest-16 | -4.41397 | 48.33658 | Lauberlac'h, France | 24/07/17 | 12 | 0.0206 |
| Brest-17 | -4.44185 | 48.34370 | Anse du Caro, France | 24/07/17 | 12 | 0.0184 |
| Brest-18 | -4.43723 | 48.35935 | Pointe Marloux, France | 24/07/17 | 12 | 0.0314 |
| Brest-19 | -4.43067 | 48.39313 | Marina Moulin Blanc Nord, France | 24/07/17 | 12 | 0.0130 |
| Brest-2 | -4.45367 | 48.29334 | Lanvéoc, France | 08/07/17 | 12 | 0.0216 |
| Brest-20 | -4.43248 | 48.39002 | Marina Moulin Blanc Sud, France | 24/07/17 | 11 | 0.0519 |
| Brest-21 | -4.47392 | 48.37994 | Brest commerce A, France | 25/07/17 | 12 | 0.0335 |
| Brest-22 | -4.47321 | 48.38280 | Brest commerce B, France | 25/07/17 | 12 | 0.0216 |
| Brest-23 | -4.48110 | 48.37762 | Brest commerce C, France | 25/07/17 | 12 | 0.0325 |
| Brest-24 | -4.45348 | 48.32235 | Pointe de l'Armorique, France | 25/07/17 | 11 | 0.0413 |
| Brest-25 | -4.55102 | 48.35945 | Anse de Sainte-Anne, France | 25/07/17 | 12 | 0.0152 |
| Brest-26 | -4.52923 | 48.36012 | Pointe du Porzic Est, France | 25/07/17 | 12 | 0.0130 |
| Brest-27 | -4.49093 | 48.37732 | Marina du Chateau A, France | 25/07/17 | 12 | 0.0152 |
| Brest-28 | -4.48892 | 48.38018 | Marina du Chateau B, France | 25/07/17 | 11 | 0.0519 |
| Brest-29 | -4.45585 | 48.37470 | Brest commerce D, France | 25/07/17 | 12 | 0.0206 |
| Brest-3 | -4.50891 | 48.28444 | Le Fret, France | 08/07/17 | 12 | 0.0076 |
| Brest-30 | -4.78104 | 48.35904 | Le Conquet, France | 27/07/17 | 12 | 0.0206 |
| Brest-31 | -4.77214 | 48.34305 | Penzer, France | 27/07/17 | 12 | 0.0087 |
| Brest-32 | -4.76820 | 48.32847 | Pointe Saint-Mathieu, France | 27/07/17 | 12 | 0.0108 |
| Brest-33 | -4.69895 | 48.33888 | Plougonvelin, France | 27/07/17 | 12 | 0.0292 |
| Brest-34 | -4.61538 | 48.33758 | Petit Minou, France | 27/07/17 | 12 | 0.0152 |
| Brest-35 | -4.58137 | 48.34809 | Le Mengant, France | 27/07/17 | 12 | 0.0119 |
| Brest-36 | -4.26814 | 48.27982 | Térénez, France | 05/08/17 | 12 | 0.0184 |
| Brest-37 | -4.51477 | 48.20070 | île Vierge, France | 06/08/17 | 12 | 0.0141 |
| Brest-38 | -4.34048 | 48.40575 | Elorn 2, France | 21/09/17 | 7 | 0.0148 |
| Brest-39 | -4.34317 | 48.40582 | Elorn 2B, France | 21/09/17 | 12 | 0.0357 |
| Brest-4 | -4.53997 | 48.29194 | L'île du Renard, France | 08/07/17 | 12 | 0.0065 |
| Brest-40 | -4.34898 | 48.40385 | Elorn St Jean, France | 21/09/17 | 11 | 0.0153 |
| Brest-41 | -4.35612 | 48.40648 | Elorn 3, France | 21/09/17 | 9 | 0.0216 |
| Brest-42 | -4.37208 | 48.39705 | Elorn 6, France | 21/09/17 | 12 | 0.0249 |
| Brest-43 | -4.38280 | 48.39553 | Elorn 9, France | 21/09/17 | 12 | 0.0216 |
| Brest-44 | -4.38205 | 48.39300 | Elorn 11, France | 21/09/17 | 12 | 0.0238 |
| Brest-45 | -4.38670 | 48.39298 | Elorn 14, France | 21/09/17 | 12 | 0.0173 |
| Brest-46 | -4.39465 | 48.39028 | Elorn 15, France | 21/09/17 | 12 | 0.0184 |

*Continued on next page*

Table S1 – *Continued from previous page*

| pop | longitude | latitude | locality | date | N | missing data |
|---|---|---|---|---|---|---|
| Brest-5 | -4.54507 | 48.31460 | Roscanvel, France | 08/07/17 | 12 | 0.0249 |
| Brest-6 | -4.57300 | 48.30710 | La Fraternité, France | 08/07/17 | 12 | 0.0173 |
| Brest-7 | -4.55452 | 48.29658 | Quelern, France | 09/07/17 | 12 | 0.0141 |
| Brest-8 | -4.53207 | 48.33974 | Pointe des Espagnols, France | 09/07/17 | 8 | 0.0568 |
| Brest-9 | -4.58254 | 48.31834 | Îlot des Capucins, France | 09/07/17 | 12 | 0.0141 |
| BretN_g1 | -3.71707 | 48.70014 | venizella, Guimaec, France | | 10 | 0.0104 |
| BretN_g2 | -3.69819 | 48.69000 | poul roudou, Locquirec, France | | 10 | 0.0078 |
| BretN_g3 | -3.67690 | 48.68796 | lezingar, Locquirec, France | | 10 | 0.0104 |
| BretN_g4 | -3.64643 | 48.69481 | pors ar villiec, Locquirec | | 10 | 0.0052 |
| BretN_i1 | -3.54638 | 48.72863 | Léguer estuary, South point, France | | 9 | 0.0072 |
| BretN_i2 | -3.54230 | 48.72895 | Banc du Guer sud, France | | 10 | 0.0117 |
| BretN_i3 | -3.58156 | 48.72139 | Locquémo, France | | 10 | 0.0078 |
| BretN_i4 | -3.58371 | 48.72608 | Locquemo, France | | 10 | 0.0065 |
| BretN_i5q1 | -3.58283 | 48.68886 | Beg ar form, Saint-Michel en Grève, France | | 10 | 0.0091 |
| BretN_i5q2 | -3.58271 | 48.68914 | Beg ar form, Saint-Michel en Grève, France | | 10 | 0.0039 |
| BretN_i6 | -3.61713 | 48.68507 | Beg Douar, Plestin Les Grèves, France | | 10 | 0.0078 |
| BretN_i8 | -3.80425 | 48.72304 | Roc'h Goalen, Plougasnou, France | | 10 | 0.0312 |
| CAD | -6.29842 | 36.52719 | Cadix, Spain | 2016 | 15 | 0.0294 |
| Can | 3.69184 | 43.41226 | Sète, canal, France | | 3 | 0.0909 |
| Cart | -1.80842 | 49.37165 | Carteret, France | 2015 | 29 | 0.0331 |
| CER_001 | -4.44887 | 48.38265 | Brest port, France | 14/12/16 | 25 | 0.0135 |
| Cha | -1.08059 | 46.03229 | Charente, France | | 12 | 0.0119 |
| CHE | -2.29965 | 47.23232 | Pointe de Chemoulin, France | 2016 | 30 | 0.0242 |
| CHE_001 | -1.62336 | 49.64574 | Cherbourg, France | 02/12/16 | 25 | 0.0156 |
| Cher-A | -1.61953 | 49.64530 | Cherbourg A, France | 01/08/17 | 12 | 0.0227 |
| Cher-B | -1.61981 | 49.64956 | Cherbourg B, France | 01/08/17 | 12 | 0.0249 |
| Cher-C | -1.63244 | 49.67307 | Cherbourg C, France | 01/08/17 | 12 | 0.0390 |
| Cher-D | -1.62020 | 49.64818 | Cherbourg D, France | 01/08/17 | 12 | 0.0487 |
| Cher-E | -1.62080 | 49.64208 | Cherbourg E, France | 01/08/17 | 12 | 0.0433 |
| Cher-F | -1.62342 | 49.64566 | Cherbourg F, France | 01/08/17 | 11 | 0.0342 |
| Cher-G | -1.61964 | 49.63846 | Cherbourg G, France | 01/08/17 | 11 | 0.0614 |
| Cher03 | -1.61994 | 49.63942 | Cherbourg, France | | 30 | 0.0108 |
| Cher15 | -1.62332 | 49.64557 | Cherbourg, France | 2015 | 30 | 0.0212 |
| Cher16 | -1.62332 | 49.64557 | Cherbourg, France | 2016 | 99 | 0.0365 |

*Continued on next page*

Table S1 – *Continued from previous page*

| pop | longitude | latitude | locality | date | N | missing data |
|---|---|---|---|---|---|---|
| CHR | -71.51917 | 41.38692 | Charlestown, Snug Harbor Marina, USA | 2009 | 12 | 0.0206 |
| COR | -8.41393 | 43.37011 | Coruña, Galicia, Spain | 2016 | 15 | 0.0069 |
| CRE | -8.97390 | 38.48047 | Praia do Creio, Arrábida, Spain | 2016 | 15 | 0.0078 |
| Croa | 13.62091 | 45.07155 | Croatia | | 14 | 0.0547 |
| D15A | 2.93460 | 54.32470 | D15-A, North Sea | 03/10/15 | 8 | 0.1445 |
| Dieppe | 1.09670 | 49.93556 | Dieppe, France | | 48 | 0.0095 |
| Din | -2.04937 | 48.63913 | Dinard, France | | 13 | 0.0649 |
| DKL | -15.93000 | 23.72000 | Dahkla, Morocco | | 10 | 0.1013 |
| Dun | 2.42178 | 51.10313 | Dunkerque, France | 2016 | 30 | 0.0030 |
| Ena11 | -2.07507 | 48.64035 | Saint-Enogat, Dinard, France | | 7 | 0.0816 |
| Engl | -0.93377 | 49.39478 | Englesqueville, France | 2015 | 30 | 0.0177 |
| EST_001 | -2.53370 | 47.30370 | Le Croisic, France | | 25 | 0.0088 |
| F1 | | | F1 | | 6 | 0.0195 |
| F2 | | | F2 | | 132 | 0.0364 |
| Far | -7.93884 | 37.01568 | Faro, Portugal | | 8 | 0.0633 |
| FAR | -7.93884 | 37.01568 | Faro, Portugal | 2016 | 15 | 0.0190 |
| Fer11 | -3.96309 | 48.72045 | ballast ferry | 2011 | 38 | 0.0123 |
| Fer13 | -3.96309 | 48.72045 | ballast ferry Armorique | 16/05/13 | 27 | 0.0077 |
| FINO | 7.15830 | 55.19500 | FINO 3, North Sea | 23/09/15 | 6 | 0.2186 |
| GA | 6.24164 | 62.46399 | GAS, Gaseid, Alesund, Norway | 2015 | 36 | 0.0400 |
| Gdansk | 18.52794 | 54.60956 | Gulf of Gdansk, Poland | 23/08/12 | 12 | 0.1223 |
| Gran | -1.59947 | 48.84003 | Granville, France | 2015 | 31 | 0.0201 |
| GrCa | -1.03633 | 49.40803 | Grancamp, France | 2015 | 29 | 0.0233 |
| Groix | -3.45911 | 47.64701 | Groix, France | | 48 | 0.0249 |
| HARW | 1.28130 | 51.93480 | Harwich, North Sea | 15/11/15 | 11 | 0.1181 |
| Havre_A | 0.11327 | 49.48586 | Le Havre, France | 2016 | 15 | 0.0069 |
| Havre_B | 0.11612 | 49.47316 | Le Havre, France | 2016 | 15 | 0.0303 |
| Havre_C | 0.12268 | 49.47000 | Le Havre, France | 2016 | 15 | 0.0130 |
| Havre_D | 0.13867 | 49.46841 | Le Havre, France | 2016 | 15 | 0.0139 |
| Havre_E | 0.15604 | 49.46831 | Le Havre, France | 2016 | 15 | 0.0104 |
| Havre_F | 0.15841 | 49.47585 | Le Havre, France | 2016 | 15 | 0.0095 |
| Havre_G | 0.17341 | 49.47242 | Le Havre, France | 2016 | 15 | 0.0087 |
| Havre_H | 0.09767 | 49.48713 | Le Havre, France | 2016 | 15 | 0.0052 |
| Havre_I | 0.16510 | 49.45401 | Le Havre, France | 2016 | 15 | 0.0173 |

*Continued on next page*

Table S1 – *Continued from previous page*

| pop | longitude | latitude | locality | date | N | missing data |
|---|---|---|---|---|---|---|
| Havre-J | 0.11832 | 49.48893 | Le Havre, France | 02/07/17 | 15 | 0.0502 |
| Her | 25.14420 | 35.33800 | Heraklion, Greece | | 35 | 0.0171 |
| Holl | 5.42400 | 53.31000 | Wadden Sea, The Netherlands | | 23 | 0.0056 |
| HORN | 7.81100 | 55.47890 | Horns Rev, North Sea | 10/06/15 | 2 | 0.1299 |
| HOU_001 | -2.93723 | 47.41967 | Ile de Houat, France | 10/01/17 | 25 | 0.0192 |
| JapSea | 135.09238 | 38.73579 | Katolikova, Japan Sea | | 16 | 0.1631 |
| Jer | -2.02038 | 49.17308 | JER, Jersey, France | 2016 | 30 | 0.0143 |
| JUM | -3.90044 | 47.83397 | La Jument, France | 2016 | 30 | 0.0104 |
| K10B | 3.25390 | 53.36260 | K10-B, North Sea | 01/10/14 | 8 | 0.1591 |
| KOR | 128.44707 | 34.83393 | South Korea | | 10 | 0.1169 |
| L4 | -4.57230 | 48.28220 | bay of Brest (Camaret), France | 24/04/17 | 42 | 0.0247 |
| L5 | -4.48815 | 48.37733 | bay of Brest (port), France | 24/04/17 | 49 | 0.0358 |
| LaRochA | -1.22359 | 46.14988 | La Rochelle, France | 24/06/17 | 7 | 0.1410 |
| LaRochB | -1.22449 | 46.15045 | La Rochelle, France | 25/06/17 | 29 | 0.0636 |
| LeHaCM | 0.11380 | 49.45725 | Le Havre, France | 2017 | 14 | 0.0241 |
| LeHaP1 | 0.10579 | 49.46466 | Le Havre P1, France | 2017 | 15 | 0.0277 |
| LeHaP10 | 0.17332 | 49.45352 | Le Havre P10, France | 2017 | 14 | 0.0473 |
| LeHaP11 | 0.17807 | 49.45456 | Le Havre P11, France | 2017 | 15 | 0.0857 |
| LeHaP2 | 0.10996 | 49.46040 | Le Havre P2, France | 2017 | 15 | 0.0199 |
| LeHaP3 | 0.11473 | 49.45825 | Le Havre P3, France | 2017 | 15 | 0.0156 |
| LeHaP4 | 0.12137 | 49.45734 | Le Havre P4, France | 2017 | 15 | 0.0216 |
| LeHaP5 | 0.13330 | 49.45634 | Le Havre P5, France | 2017 | 15 | 0.0727 |
| LeHaP6 | 0.14413 | 49.45546 | Le Havre P6, France | 2017 | 15 | 0.0355 |
| LeHaP7 | 0.15224 | 49.45185 | Le Havre P7, France | 2017 | 15 | 0.0268 |
| LeHaP8 | 0.16015 | 49.45441 | Le Havre P8, France | 2017 | 15 | 0.0268 |
| LeHaP9 | 0.16602 | 49.45376 | Le Havre P9, France | 2017 | 15 | 0.0502 |
| LISB | -9.09260 | 38.76350 | Lisbon, Portugal | 14/02/15 | 5 | 0.1818 |
| LochEti | -5.18317 | 56.45735 | Loch Etive, Scotland, UK | | 11 | 0.1145 |
| Locq | -3.59720 | 48.73691 | Locquemeau, France | | 37 | 0.0190 |
| LOF | 13.87800 | 68.33800 | Lofoten, Norway | 2014 | 43 | 0.0417 |
| MAB_001 | -2.21610 | 46.99760 | Maison Blanche, Noirmoutier, France | | 25 | 0.0088 |
| MAK | 22.61232 | 40.41618 | Makrigiallos, Greece | | 6 | 0.1017 |
| MOU | -4.04187 | 47.84282 | Pointe de Mousterlin, France | 2016 | 30 | 0.0242 |
| Murch | 1.75000 | 61.40000 | MCH, Murchison oil station, Norwegian Sea | | 12 | 0.0465 |

*Continued on next page*

Table S1 – *Continued from previous page*

| pop | longitude | latitude | locality | date | N | missing data |
|---|---|---|---|---|---|---|
| NUS | -51.71040 | 64.19680 | Nuuk, Greenland | 2014 | 28 | 0.0756 |
| NY-harb-A | -73.79327 | 40.79521 | NYC Throgs Neck Bridge, USA | 09/06/04 | 9 | 0.0534 |
| NY-harb-B | -71.94682 | 41.02913 | Montauk, USA | 23/02/14 | 7 | 0.0686 |
| NY-harb-C | -74.01726 | 40.65532 | Brooklyn piers, USA | | 10 | 0.0260 |
| ORT | -7.80508 | 43.72246 | Ortigueira, Galicia, Spain | 2016 | 15 | 0.0139 |
| OST | -72.38419 | 41.26328 | Old Saybrook town, Hartland drive, USA | 2009 | 24 | 0.0211 |
| Ostende | 2.91815 | 51.23815 | Ostende, Belgium | | 29 | 0.0090 |
| Ouis | -0.45892 | 49.33829 | Ouistreham, France | 2015 | 30 | 0.0203 |
| Palice-A | -1.21576 | 46.15909 | La Palice, Bassin à flot, France | 23/02/18 | 18 | 0.0065 |
| Palice-B | -1.21843 | 46.15854 | La Palice, lock, France | 23/02/18 | 76 | 0.0106 |
| PEN | -2.51383 | 47.30632 | Barres de Pen Bron, France | 2016 | 30 | 0.0212 |
| PEN_001 | -2.49363 | 47.49712 | Camaret plage-Penestin, France | 11/01/17 | 25 | 0.0223 |
| Peniche | -9.37944 | 39.36844 | Peniche, Portugal | 2016 | 15 | 0.0121 |
| PET_001 | -4.47146 | 48.38119 | Brest port, France | 14/12/16 | 25 | 0.0151 |
| Pir | -1.60083 | 49.16588 | Pirou, France | 2016 | 30 | 0.0074 |
| POU_001 | -2.41596 | 47.26029 | Le Pouliguen - La Baule, France | 12/01/17 | 25 | 0.0119 |
| Prim | -3.82080 | 48.72122 | Primel, France | | 36 | 0.0177 |
| PtArm97 | -4.45355 | 48.32563 | Pointe de l'Armorique, plage des ducs d'albes, France | | 19 | 0.0444 |
| Q13A | 4.13610 | 52.19110 | Q13-A, North Sea | 28/05/14 | 12 | 0.0617 |
| QAS | -69.24030 | 77.46500 | Qaanaaq, Greenland | 2014 | 30 | 0.1364 |
| RadeBrest_R1 | -4.39973 | 48.39014 | Pont d'Iroise, France | | 10 | 0.0195 |
| RadeBrest_R2 | -4.36882 | 48.33810 | Port du Tinduff, France | | 10 | 0.0169 |
| RadeBrest_R3 | -4.50680 | 48.28679 | Le Fret, France | | 10 | 0.0091 |
| RadeBrest_R4 | -4.55083 | 48.30609 | Quélern, Roscanvel, France | | 10 | 0.0104 |
| RadeBrest_R5 | -4.57194 | 48.28199 | Pte Ste Barbe, Camaret, France | | 10 | 0.0104 |
| Rave | -1.21023 | 49.47333 | Ravenoville, France | 2015 | 30 | 0.0130 |
| Revi | -1.22933 | 49.57467 | Réville, France | 2015 | 30 | 0.0160 |
| ROC_VER | -1.40556 | 45.98358 | Rocher Vert Chaucre cote ouest Oléron, France | 05/10/16 | 25 | 0.0151 |
| RoRo | -3.38314 | 48.81866 | Roc-Rouge, France | | 32 | 0.0114 |
| Roth | -1.96213 | 48.68703 | Rothéneuf, France | | 32 | 0.0288 |
| SB1 | -2.71614 | 48.55568 | Pointe du Roselier, France | | 10 | 0.0039 |
| SB2 | -2.78885 | 48.58458 | Pordic, plage du petit havre, France | | 10 | 0.0052 |
| SCHV | 4.25820 | 52.09870 | Scheveningen, North Sea | 08/07/14 | 1 | 0.0779 |
| SoRo | -1.55248 | 48.73239 | Sol-Roc, France | | 30 | 0.0472 |

*Continued on next page*

Table S1 – *Continued from previous page*

| pop | longitude | latitude | locality | date | N | missing data |
|---|---|---|---|---|---|---|
| StAnd | 0.07980 | 49.54744 | Saint-Andrieux, France | | 15 | 0.0087 |
| StJo | 0.15264 | 49.64395 | Saint-Jouin, France | 2015 | 27 | 0.0106 |
| StLau-CBD | -69.46615 | 48.26941 | Saint Lawrence, Cap de Bon Désir, Canada | | 9 | 0.1010 |
| StLau-TD | -69.69614 | 48.13441 | Saint Lawrence, Tadoussac, Canada | | 12 | 0.1180 |
| StMalo | -2.02165 | 48.64855 | Saint-Malo port, France | 16/07/17 | 30 | 0.0156 |
| StNaz-I | -2.19894 | 47.27074 | Saint-Nazaire, France | 30/01/18 | 26 | 0.0080 |
| StNaz-II | -2.20027 | 47.27391 | Saint-Nazaire, France | 30/01/18 | 25 | 0.0348 |
| StNaz-III | -2.20259 | 47.27312 | Saint-Nazaire, France | 30/01/18 | 25 | 0.0358 |
| StNaz-IV | -2.19813 | 47.28964 | Saint-Nazaire, France | 30/01/18 | 26 | 0.0330 |
| StNaz-V | -2.19880 | 47.27941 | Saint-Nazaire, France | 30/01/18 | 26 | 0.0280 |
| SV1 | 11.13620 | 79.11230 | Svalbard – Kongsfjorden, Norway | 2012 | 6 | 0.0455 |
| SV2 | 11.13620 | 79.11230 | Svalbard – Kongsfjorden, Norway | 2013 | 9 | 0.0505 |
| SV3 | 11.13620 | 79.11230 | Svalbard – Kongsfjorden, Norway | 2014 | 12 | 0.0649 |
| SV4 | 15.60260 | 78.23810 | Svalbard – Adventfjorden, Norway | 2014 | 11 | 0.0106 |
| Tatihou | -1.23627 | 49.58673 | Tatihou, France | 09/07/05 | 24 | 0.0076 |
| Th16 | 3.69179 | 43.41227 | Thau, France | 2016 | 23 | 0.0169 |
| TH16B | 3.68782 | 43.41474 | Thau, ponton station, France | 2016 | 18 | 0.0043 |
| VHO | -8.82312 | 37.38853 | Vale dos Homens, Aljezur, Spain | 2016 | 15 | 0.0078 |
| VIG | -8.70214 | 42.26008 | Vigo, Spain | 2016 | 15 | 0.0104 |
| Vill | 0.11605 | 49.39935 | Villerville, France | 2015 | 30 | 0.0251 |
| VILL | 0.12383 | 49.40374 | Villerville, France | 2016 | 30 | 0.0108 |
| Vill16 | 0.12351 | 49.40369 | Villerville, France | 2016 | 15 | 0.0035 |
| VLR | 0.79465 | 49.87812 | Veules-les-Roses, France | 2016 | 30 | 0.0052 |
| Wim | 1.60307 | 50.77263 | Wimereux, France | 2016 | 29 | 0.0049 |

Table S2: Softwares and R packages used in this study

| software | version | reference | link |
|---|---|---|---|
| R | 3.5.2 | R Core Team, 2019 | https://www.r-project.org/ |
| Python | 3.7.1 | - | https://www.python.org/ |
| Structure | 2.3.4 | Falush et al., 2003; Pritchard, Stephens, and Donnelly, 2000 | https://web.stanford.edu/group/pritchardlab/structure.html |
| Admixture | 1.3.0 | Alexander and Lange, 2011 | http://software.genetics.ucla.edu/admixture/ |
| CLUMPAK | 1.1 | Kopelman, Mayzel, Jakobsson, Rosenberg, and Mayrose, 2015 | http://clumpak.tau.ac.il/ |
| QGIS | 3.4.2 | - | https://qgis.org/fr/site/ |
| Newhybrids | 1.1 beta 3 | Anderson and Thompson, 2002 | http://ib.berkeley.edu/labs/slatkin/eriq/software/software.htm |

| R package | version | reference |
|---|---|---|
| tidyverse | 1.2.1 | Wickham, 2017 |
| reshape2 | 1.4.3 | Wickham, 2007 |
| adegenet | 2.1.1 | T. Jombart and Ahmed, 2011; Thibaut Jombart, 2008 |
| pegas | 0.11 | Paradis, 2010 |
| hierfstat | 0.04-22 | Goudet and Jombart, 2015 |
| introgress | 1.2.3 | Gompert and Buerkle, 2010 |
| qtl | 1.44-9 | Broman, Wu, Sen, and Churchill, 2003 |
| LDheatmap | 0.99-5 | Shin, Blay, McNeney, and Graham, 2006 |
| genetics | 1.3.8.1 | Warnes, Gorjanc, Leisch, and Man, 2013 |
| hzar | 0.2-5 | Derryberry, Derryberry, Maley, and Brumfield, 2014 |
| ggpubr | 0.2 | Kassambara, 2018 |
| psych | 1.8.12 | Revelle, 2018 |
| EmpiricalBrownsMethod | 1.10.0 | Poole, Gibbs, Shmulevich, Bernard, and Knijnenburg, 2016 |
| ggstatsplots | 0.0.9 | Patil, 2019 |
| foreach | 1.4.4 | Microsoft and Weston, 2017 |
| doMC | 1.3.5 | Analytics and Weston, 2017 |
| gdistance | 1.2-2 | van Etten, 2017 |

Table S3: Across all analyses multiple selection threshold have been used to select either individuals or markers for specific analyses. Decision thresholds are presented in table S3. For statistical tests we chose $\alpha = 0.05$ as the significant threshold.

| Variable used | for what? | with what? | kept if |
|---|---|---|---|
| Missing data of individuals | full dataset | genotypes | $< 30\%$ |
| | genetic map F2 | genotypes | $< 10\%$ |
| Missing data of markers | full dataset | genotypes | $< 10\%$ |
| | $F_{ST}$ between reference populations | genotypes of *M. trossulus* pop. only | $< 30\%$ |
| $Q$ value or prob. of membership ($C$) | reference groups filtration | `Admixture` $K = 3$ (fig. S4) | $Q > 85\%$ from putative cluster |
| | *M. trossulus* individuals | full `Structure` $K = 5$ (fig. S5) | $Q_{tros} > 0.1$ |
| | list of dock mussels | `Structure` local w/o admixture $K = 3$ (fig. S20) | $\in (\text{Port}) \wedge \in (C_{dock} > 0.9)$ |
| | list of *M. edulis* in ports | `Structure` local w/o admixture $K = 3$ | $\in (\text{Port}) \wedge \in (C_{edu} > 0.5)$ |
| | list of gallo_atl in ports | `Structure` local w/o admixture $K = 3$ | $\in (\text{Port}) \wedge \in (C_{gallo\_atl} > 0.5$ |
| | list of admix gallo_atl | `Structure` w/o admixture $K = 5$ (fig. S19) | $\in (\text{focal pop.}) \wedge \in (C_{gallo\_atl} + C_{admix\_atl} > 0.9)$ |

Table S4: Reference panel groups and populations

| L1 | L2 | L3 | pop | N |
|---|---|---|---|---|
| edu | edu_am | edu_am | Bost | 14 |
| | | | CHR | 12 |
| | | | OST | 24 |
| | edu_eu | edu_eu_south | Dun (ext) | 29 |
| | | | Holl (ext) | 23 |
| | | | Ostende (ext) | 28 |
| | | | Wim (ext) | 29 |
| | | | Cha (int) | 12 |
| | | edu_eu_north | Bodo | 38 |
| | | | Berg | 38 |
| gallo | gallo_atl | gallo_atl_iber | CAD | 15 |
| | | | COR | 15 |
| | | | CRE | 15 |
| | | | DKL | 10 |
| | | | Far | 8 |
| | | | FAR | 14 |
| | | | ORT | 15 |
| | | | Peniche | 15 |
| | | | VHO | 15 |
| | | | VIG | 15 |
| | | gallo_atl_brit | Prim | 36 |
| | gallo_med | gallo_med_east | Croa | 14 |
| | | | Her | 35 |
| | | | MAK | 6 |
| | | gallo_med_west | Can | 3 |
| | | | Th16 | 23 |
| | | | TH16B | 18 |
| | | | Bizerte | 12 |
| tros | tros_am | tros_am | LochEti | 11 |
| | | | StLau-CBD | 9 |
| | | | StLau-TD | 11 |
| | tros_eu | tros_eu_baltic | Gdansk | 12 |
| | tros_pac | tros_pac_west | JapSea | 16 |

Table S5: Linkage map – (i) genetic map produced from lab F2 crosses, (ii) markers on the same contig and (iii) left markers with unknown relation.

| | locus | linkage group | position (cM) | unlinked set |
|---|---|---|---|---|
| (i) | 063 | 1 | 0.00 | • |
| | 067 | 1 | 22.88 | |
| | 607 | 1 | 24.23 | |
| | 062 | 1 | 25.11 | • |
| | 210 | 1 | 28.33 | |
| | 055 | 1 | 28.77 | |
| | 034 | 1 | 31.99 | |
| | 115 | 1 | 32.44 | |
| | 064 | 2 | 0.00 | |
| | 026 | 2 | 1.36 | |
| | 047 | 2 | 1.36 | |
| | 160 | 2 | 1.36 | • |
| | 161 | 2 | 3.65 | |
| | 052 | 3 | 0.00 | • |
| | 007 | 3 | 9.70 | |
| | 610 | 3 | 27.41 | |
| | 015 | 4 | 0.00 | |
| | 155 | 4 | 2.29 | • |
| | 900 | 4 | 14.32 | |
| | 142 | 5 | 0.00 | |
| | 148 | 5 | 3.22 | |
| | 184 | 5 | 3.22 | • |
| | 061 | 6 | 0.00 | |
| | 080 | 6 | 10.00 | • |
| | 082 | 7 | 0.00 | • |
| | 802 | 7 | 10.00 | |
| | 144 | 8 | 0.00 | |
| | 145 | 8 | 10.00 | • |
| | 001 | 9 | 0.00 | • |
| | 022 | 10 | 0.00 | • |
| | 073 | 11 | 0.00 | • |
| | 094 | 12 | 0.00 | • |
| | 164 | 13 | 0.00 | • |
| | 180 | 14 | 0.00 | • |
| | 211 | 15 | 0.00 | • |
| | 617 | 16 | 0.00 | • |
| (ii) | 202 | R_L02_mira_Contig1 | 2420 (bp) | • |
| | 602 | R_L02_mira_Contig1 | 3684 (bp) | |
| | 604 | R_L02_mira_Contig1 | 6232 (bp) | |
| (iii) All other markers not in the linkage map | | | | • |

General Note

| $p$ value | symbol |
|---|---|
| $p < 0.001$ | *** |
| $0.001 < p < 0.01$ | ** |
| $0.01 < p < 0.05$ | * |
| $p > 0.05$ | ns |

When statistically significant, results are presented in bold.

Table S6: Fst values between reference groups.

| Level | group 1 | group 2 | $F_{ST}$ |
|---|---|---|---|
| L1 | edu | gallo | **0.8127*** ** |
| | edu | tros | **0.7226*** ** |
| | gallo | tros | **0.8086*** ** |
| L2 | gallo_med | gallo_atl | **0.3823*** ** |
| | edu_eu | edu_am | **0.4845*** ** |
| L3 | gallo_med_west | gallo_med_east | **0.0586*** ** |
| | edu_eu_south_int | edu_eu_south_ext | 0.0024 (ns) |
| | edu_eu_south_int | edu_eu_north | **0.2086*** ** |
| | edu_eu_south_ext | edu_eu_north | **0.3129*** ** |
| | gallo_atl_ext | gallo_atl_int | **0.1555*** ** |

Table S7: Fst values and significance between ports.

| | cher | havre | brest | stmalo |
|---|---|---|---|---|
| havre | **0.0235*** ** | | | |
| brest | **0.0046*** ** | **0.0245*** ** | | |
| stmalo | 0.0078 (ns) | **0.0390*** ** | 0.0058 (ns) | |
| stnaz | 0.0071 (ns) | **0.0304*** ** | **0.0067** ** | 0.0043 (ns) |

Table S8: Fst values and significance between Cherbourg years.

| | cher03 | cher15 | cher16 |
|---|---|---|---|
| cher15 | **0.0066** ** | | |
| cher16 | **0.0097** * | 0.0009 (ns) | |
| cher17 | 0.0032 (ns) | -0.0017 (ns) | -0.0007 (ns) |

Table S9: Fst values and significance between naturally and Norway admixed.

| | ABD | HZSB | GAS | JER | LOF |
|---|---|---|---|---|---|
| HZSB | **0.0429*** ** | | | | |
| GAS | **0.0736** * | **0.1021** * | | | |
| JER | **0.0974*** ** | **0.1439*** ** | 0.0335 (ns) | | |
| LOF | **0.0372*** ** | **0.0743*** ** | 0.0047 (ns) | 0.0414 (ns) | |
| MCH | 0.0346 (ns) | 0.0462 (ns) | 0.1795 (ns) | **0.2330** * | 0.1163 (ns) |

Table S10: Statistical tests of the differences in ancestry estimates between populations of implicating Atl. *M. galloprovincialis* and *M. edulis* (i.e. Norway admixed and naturally admixed). The Dwass-Steel-Crichtlow-Fligner test is used to compute post-hoc pairwise comparisons, the statistic (W) and corrected $p$-values for multiple tests are reported for each type of ancestry.

| group1 | group2 | edu_eu_south | | gallo_atl | | gallo_med | | edu_am | |
|---|---|---|---|---|---|---|---|---|---|
| | | W | p.value | W | p.value | W | p.value | W | p.value |
| JER | ABD | -8.0014855 | 0.0000003 | 8.093009 | 0.0000003 | -1.9301499 | 1.0000000 | 0.4648148 | 1.0000000 |
| JER | MCH | -6.3869294 | 0.0000784 | 6.386518 | 0.0000392 | -3.3998864 | 0.8073574 | -1.2188032 | 1.0000000 |
| JER | HZSB | -8.2758887 | 0.0000002 | 8.291331 | 0.0000002 | -2.6247521 | 1.0000000 | 1.2462346 | 1.0000000 |
| ABD | MCH | -5.7267576 | 0.0004261 | 5.920113 | 0.0001286 | -2.5251363 | 1.0000000 | -1.6665649 | 1.0000000 |
| ABD | HZSB | -4.2334401 | 0.0152819 | 4.159408 | 0.0135876 | -0.5622536 | 1.0000000 | 1.5813721 | 1.0000000 |
| MCH | HZSB | 1.5074295 | 1.0000000 | -1.750562 | 0.7669834 | 2.3974014 | 1.0000000 | 2.9963772 | 0.1888831 |
| GAS | LOF | -1.3324952 | 1.0000000 | 1.675714 | 0.7832573 | -0.4038931 | 1.0000000 | 1.4839329 | 1.0000000 |
| GAS | JER | 5.8227527 | 0.0003818 | -3.283418 | 0.0776347 | 0.9194070 | 1.0000000 | -5.9576512 | 0.0001795 |
| GAS | ABD | -1.2763083 | 1.0000000 | 6.491894 | 0.0000315 | -0.3242880 | 1.0000000 | -6.6249430 | 0.0000233 |
| GAS | MCH | -4.5553253 | 0.0090960 | 6.585699 | 0.0000296 | -2.0567182 | 1.0000000 | -4.8952169 | 0.0033482 |
| GAS | HZSB | -3.0020824 | 0.1682154 | 6.555299 | 0.0000296 | -0.6873292 | 1.0000000 | -7.1242722 | 0.0000059 |
| LOF | JER | 7.9519649 | 0.0000003 | -5.949607 | 0.0001286 | 1.9547265 | 1.0000000 | -8.4598018 | 0.0000000 |
| LOF | ABD | 0.7416394 | 1.0000000 | 6.047916 | 0.0001051 | 0.4258784 | 1.0000000 | -10.4083896 | 0.0000000 |
| LOF | MCH | -4.4370894 | 0.0106137 | 7.028091 | 0.0000090 | -2.0570260 | 1.0000000 | -6.6559353 | 0.0000233 |
| LOF | HZSB | -2.5689731 | 0.3136611 | 7.008418 | 0.0000090 | -0.0492258 | 1.0000000 | -12.0687157 | 0.0000000 |

Table S11: Comparisons of ancestry estimates in dock mussel populations. See Table S10 for details.

| group1 | group2 | edu_eu_south | | gallo_atl | | gallo_med | | edu_am | |
|---|---|---|---|---|---|---|---|---|---|
| | | W | p.value | W | p.value | W | p.value | W | p.value |
| brest | havre | 10.7655111 | 0.0000000 | -15.6453266 | 0.0000000 | -3.341636 | 0.0714903 | 5.979967 | 0.0006899 |
| brest | cher | 0.5032010 | 1.0000000 | -10.9995502 | 0.0000000 | 6.803901 | 0.0000110 | 2.996186 | 0.1429073 |
| brest | stnaz | 0.9072883 | 1.0000000 | -5.6694271 | 0.0003583 | 3.482114 | 0.0674667 | -1.141112 | 1.0000000 |
| brest | stmalo | -2.9745824 | 0.1482498 | -3.4396767 | 0.0627953 | 5.240644 | 0.0012351 | -2.577176 | 0.2506348 |
| havre | cher | -13.4998443 | 0.0000000 | 6.6257054 | 0.0000222 | 13.215188 | 0.0000000 | -3.266336 | 0.1084834 |
| havre | stnaz | -7.7970358 | 0.0000003 | 6.6026763 | 0.0000222 | 7.137681 | 0.0000044 | -5.056126 | 0.0038002 |
| havre | stmalo | -8.2353098 | 0.0000001 | 4.5053667 | 0.0070604 | 7.675113 | 0.0000008 | -5.016890 | 0.0038002 |
| cher | stnaz | 0.8986060 | 1.0000000 | 2.4963584 | 0.2841525 | -1.389627 | 0.9541097 | -3.234122 | 0.1084834 |
| cher | stmalo | -3.6887206 | 0.0511747 | 1.8065668 | 0.6554651 | 3.202155 | 0.0767377 | -3.980020 | 0.0358256 |
| stnaz | stmalo | -3.6202551 | 0.0511747 | 0.3570753 | 1.0000000 | 3.302863 | 0.0714903 | -1.562515 | 0.8759137 |

Table S12: Comparisons of ancestry estimates in Cherbourg temporal sampling. See Table S10 for details.

| group1 | group2 | edu_eu_south | | gallo_atl | | gallo_med | | edu_am | |
|---|---|---|---|---|---|---|---|---|---|
| | | W | p.value | W | p.value | W | p.value | W | p.value |
| cher17 | cher16 | -2.6860867 | 0.4230421 | 2.3094012 | 0.5026203 | 1.8889989 | 0.6677609 | 1.5834060 | 1 |
| cher17 | cher03 | 1.6852922 | 0.8582597 | 1.6552149 | 0.8892385 | -2.6432869 | 0.3065542 | 0.3310927 | 1 |
| cher17 | cher15 | 0.4714781 | 1.0000000 | -1.4395540 | 0.9074246 | 0.4614467 | 1.0000000 | 0.2006680 | 1 |
| cher16 | cher03 | 3.7937987 | 0.1074177 | 0.4877319 | 1.0000000 | -4.2160701 | 0.0422489 | -0.8576754 | 1 |
| cher16 | cher15 | 2.4189968 | 0.4273574 | -3.1784465 | 0.3622642 | -0.8510679 | 1.0000000 | -0.7791183 | 1 |
| cher03 | cher15 | -1.1499584 | 1.0000000 | -2.4254679 | 0.5026203 | 2.6344868 | 0.3065542 | -0.0209115 | 1 |

Table S13: Comparisons of ancestry estimates in Atl. *M. galloprovincialis* from Brittany. See Table S10 for details.

| group1 | group2 | edu_eu_south | | gallo_atl | | gallo_med | | edu_am | |
|---|---|---|---|---|---|---|---|---|---|
| | | W | p.value | W | p.value | W | p.value | W | p.value |
| gallo_atl_brest_bay | gallo_atl_brest | -4.540429 | 0.0024335 | 4.074363 | 0.0072794 | 2.8645769 | 0.2354177 | -1.909974 | 0.3243082 |
| gallo_atl_brest_bay | gallo_atl_int | -12.819450 | 0.0000000 | 12.408811 | 0.0000000 | -0.3787987 | 1.0000000 | -8.974752 | 0.0000000 |
| gallo_atl_brest | gallo_atl_int | -10.709842 | 0.0000000 | 10.484107 | 0.0000000 | -2.4195419 | 0.2394843 | -7.325735 | 0.0000006 |

Table S14: Estimation of the number of generations since admixture in ports.

| port | average r | sd(r) | generations |
|---|---|---|---|
| brest | 0.05873 | 0.00007 | 5.87 |
| cher | 0.07082 | 0.00006 | 7.08 |
| cher03 | 0.12713 | 0.00061 | 12.71 |
| cher15 | 0.07670 | 0.00031 | 7.67 |
| cher16 | 0.06290 | 0.00008 | 6.29 |
| cher17 | 0.06472 | 0.00013 | 6.47 |
| havre | 0.13902 | 0.00018 | 13.90 |
| stmalo | 0.08277 | 0.00066 | 8.28 |
| stnaz | 0.03472 | 0.00005 | 3.47 |
